# Age-dependent disease tolerance to SARS-CoV-2 infection

**DOI:** 10.64898/2026.06.27.734955

**Authors:** Sonia Trikha Rastogi, Miguel Mesquita, Diogo Martins Fonseca, Sara Salazar, Silvia Cardoso, Pedro Faisca, Bernhard Drotleff, Marta Alenquer, Jean-Christophe Lone, Veronica Miguel, David Sancho, Laura Herrero, Tiago Paixão, Maria João Amorim, Elisa Jentho, Luis Graça, Jamil Zola Kitoko, Miguel P. Soares

## Abstract

Disease tolerance limits infectious disease severity through tissue damage control mechanisms that do not target pathogens directly. Here we demonstrate that age-dependent decline in adipose tissue lipolysis compromises disease tolerance to SARS-CoV-2 infection. Young adult mice exhibited robust adipocyte lipolysis and 80% survival, whereas old mice showed impaired adipocyte lipolysis and only 20% survival. Genetic repression of adipocyte lipolysis eliminated this age-dependent survival advantage without affecting viral titers, revealing that adipocyte lipolysis is essential for disease tolerance to SARS-CoV-2 in young adults. Impaired adipocyte lipolysis in aged mice was associated with a plasma lipidomic signature that predicts COVID-19 severity and mortality in three independent human cohorts. Mechanistically, adipocyte lipolysis provides free fatty acids (FFA) to support bone marrow emergency myelopoiesis, through CD36- and CPT1-dependent FFA cellular uptake and mitochondrial import, respectively. Bone marrow derived monocytes migrate to the lung via CCL2/CCR2-dependent mechanism where they enforce an immune-metabolic communication network with parenchymal cells to sustain lung structure and function. This circuit is not required to confer protection against influenza infection, revealing pathogen-specific disease tolerance mechanisms. These findings reveal adipose tissue catabolism as a central age-dependent factor responsible for exacerbated COVID-19 mortality in aged populations.

**One-Sentence Summary:** SARS-CoV-2 infection induces adipose tissue lipolysis to release fatty acids that drive myelopoiesis and monocyte production for lung protection and COVID-19 disease tolerance, but this protective circuit declines with age, increasing disease severity in the elderly.

## INTRODUCTION

Host response to infection encompasses two distinct defense strategies: resistance, which targets and reduces pathogen burden and disease tolerance, which limits tissue damage without targeting pathogens directly^1,2^. Although immune-mediated resistance mechanisms have been extensively studied, how the immune system contributes to tissue damage control^3^ and disease tolerance to infection remains poorly understood^4^.

Severe acute respiratory syndrome coronavirus 2 (SARS-CoV-2) infections can progress towards acute respiratory distress syndrome (ARDS) underlying the lethal outcome of the coronavirus disease 2019 (COVID-19)^5,6^. While viral load correlates with ARDS onset and severity as well as COVID-19 mortality^7^, this association is confounded by aging^8–10^, among other independent risk factors^11^. This suggests that age-dependent COVID-19 susceptibility results not only from defective viral clearance but also from compromised disease tolerance^12,13^. In keeping with this notion, aging has been recently associated with a decline of disease tolerance to infection^14,15^.

Disease tolerance relies on reprogramming of host energy metabolism^16–19^, via integrated mechanisms that control glucose production via gluconeogenesis^20,21^ as well as free fatty acids (FFA) and glycerol mobilization via adipocyte lipolysis^22^. Aging is associated with a pronounced decline of adipocyte lipolysis^23–25^, suggesting that this age-dependent vulnerability might compromise disease tolerance to severe infections in the elderly^26^. Here we tested this hypothesis for COVID-19 and found that age-dependent decline of adipocyte lipolysis enhances SARS-CoV-2 lethality in the elderly, a hallmark of the COVID-19 pandemic^8–10^.

## RESULTS

### Adipose tissue lipolysis confers age-dependent disease tolerance to MA-SARS-CoV-2

To examine age-dependent effects in COVID-19 pathogenesis, we infected young adult (10-16 weeks) and elderly (>20 months old) C57BL/6J mice, corresponding to ∼20-30 and >65 year-old humans, respectively, with mouse-adapted (MA)-SARS-CoV-2^27^. The viral inoculum was calibrated to cause 20-40% lethality in young mice (LD20), while resulting in 80% lethality in old mice (LD80) (Fig. 1A,B). Lung viral titers were slightly increased (0.5 log_10_) in old *vs.* young mice, as assessed 5 days post-infection (dpi) (Fig. 1C), before the onset of lethality in old mice (Fig. 1B). This suggests that age-dependent increase in mortality is not readily explained by an overt viral titer increase, consistent with SARS-CoV-2 load being only modestly dependent on the age^28^ and not fully explaining the increased severity and mortality of elderly patients^29^.

**Figure 1.**
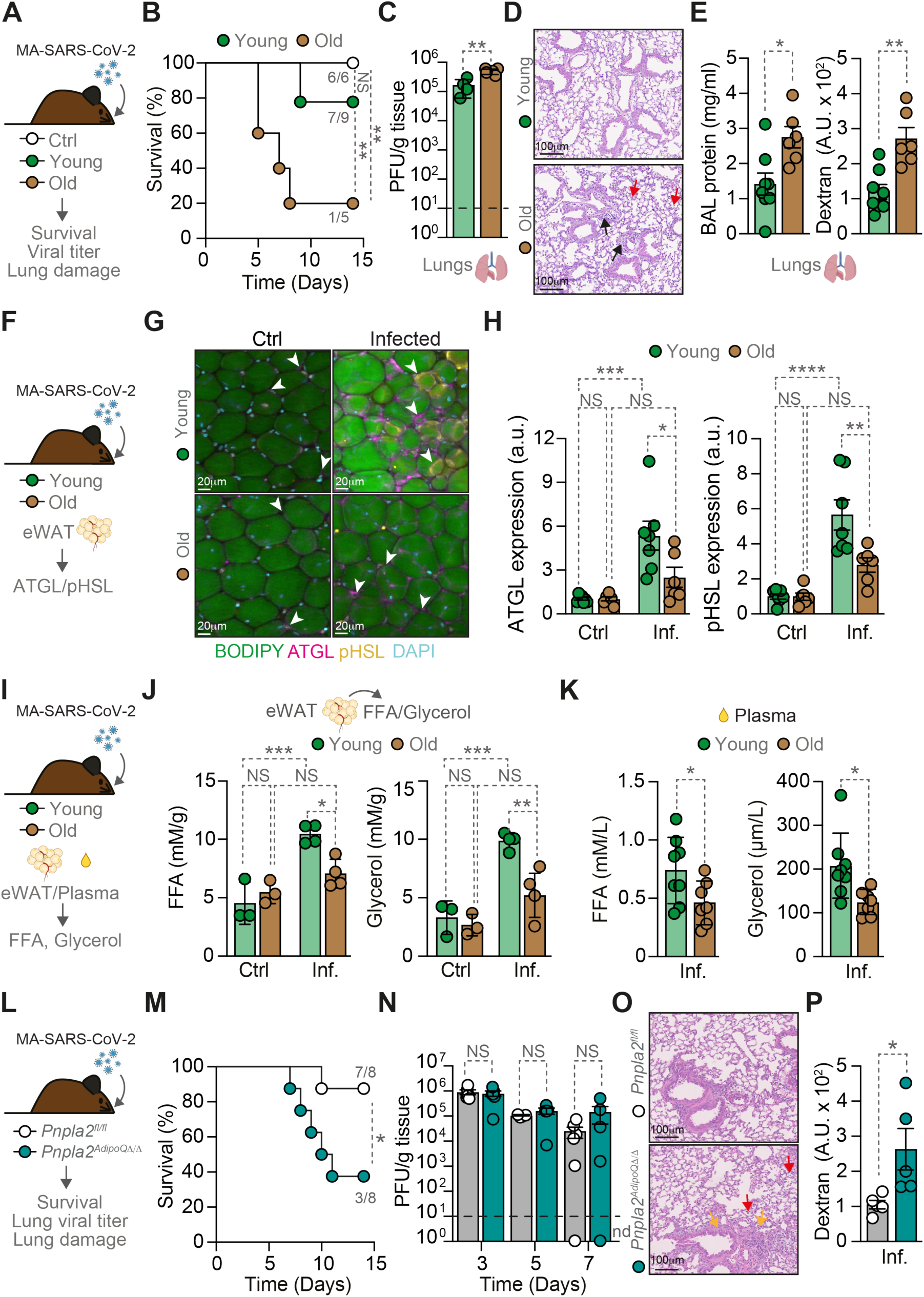
Adipose tissue lipolysis confers age-dependent disease tolerance to MA-SARS-CoV-2 infection. (**A**) Schematic of young (10-16 weeks) and old (>20 months) C57BL/6J mice infected intranasally with mouse-adapted (MA)-SARS-CoV-2 strain. (**B**) Survival curves of old control as well as young *vs*. old infected (inf.) mice (n=5-9 mice *per* group, males). (**C**) Lung viral titers measured as plaque forming units (PFU) *per* g of tissue at 5 days post-infection (dpi) (n=4-5 mice *per* group, males). Dashed line indicates limit of detection. (**D**) Representative H&E-stained lung sections at 5 dpi showing alveolar (red arrows) and perivascular infiltrates (black arrows). (**E**) Protein concentration in the bronchoalveolar lavage (BAL) and dextran-rhodamine leakage to the blood after intranasal instillation at 5 dpi (n=6-8 mice *per* group, males & females). (**F**) Schematic of lipolytic enzymes expression analysis in the epididymal white adipose tissue (eWAT) from young and old infected mice at 5 dpi. (**G**) Representative immunofluorescence images of eWAT from young *vs*. old control and infected mice, stained for BODIPY 493/503 (lipid droplets, green), ATGL (magenta), pHSL (yellow), and DAPI (nuclei, cyan). White arrowheads highlight positive staining of lipolytic enzymes. (**H**) Quantification of ATGL and pHSL from (G) (n=6-7 mice *per* group, males, a.u.; arbitrary unit). (**I**) Quantification of free fatty acids (FFA) and glycerol (**J**) released from eWAT explants (n=3-4 mice *per* group, males) and (**K**) in plasma (n = 7-8 mice *per* group, males) from young *vs*. old control and infected mice at 5 dpi. (**L**) Adipocyte-specific ATGL knockout (*Pnpla2^AdipoQΔ/Δ^*) mice and littermate (*Pnpla2^fl/fl^*) controls infected with MA-SARS-CoV-2 and monitored for (**M**) Survival (n=8 mice *per* group, males & females) and (**N**) Lung viral titers at 3, 5, and 7 dpi (n=4-6 mice *per* group *per* timepoint, males & females). Dashed line indicates limit of detection, nd: not determined. (**O**) Representative H&E-stained lung sections at 5 dpi showing alveolar (red arrows) and interstitial infiltrates (yellow arrows). **(P)** Dextran-rhodamine leakage to the blood after intranasal instillation at 5 dpi (n=4-5 mice *per* group, males & females) Data represented as mean *±* SD (J, K) or SEM (C, E, H, N, P). Numbers indicate surviving mice *per* total mice at day 14 (B, M). Circles represent individual mice (C, E, H, J, K, N, P). Survival curves were compared using the Cox proportional hazards model adjusted for experiment, with pairwise log-rank tests between-group comparisons (B, M). *p* values were determined using Log-rank test (B, M), unpaired t-test (C, E, K, P) and two-way ANOVA (H, J, N). NS, not significant; *p < 0.05; **p < 0.01; ***p < 0.001; ****p < 0.0001. Data pooled from three independent experiments (B) or two independent experiments with a similar trend (C, E, H, K, M, N, P) or from one experiment (D, J, O).

Histological analysis of the lungs confirmed a more prominent inflammation (Fig. 1D) and ARDS development (Fig. 1D,E) in old *vs.* young infected-mice at 5 dpi, as demonstrated by bronchoalveolar lavage (BAL) protein content and dextran extravasation from airspaces due to lung alveolar-capillary barrier disruption (Fig. 1E). This suggests that age-dependent COVID-19 mortality is attributed to defective tissue (lung) damage control^3^.

We hypothesized that age-dependent decrease in adipocyte lipolysis^23–25^ limits FFA mobilization, an energy substrate we have recently found to be essential to support tissue damage control and disease tolerance to bacterial infection^22^. MA-SARS-CoV-2 infection was associated with the upregulation of the rate-limiting lipolytic enzymes, adipose triglyceride lipase (ATGL) and phosphorylated hormone-sensitive lipase (pHSL) in epididymal white adipose tissue (eWAT) from young but not old mice, as assessed by immunofluorescence analysis (Fig. 1F-H; Fig. S1) and confirmed by western blot analysis (Fig. S2). This was corroborated by an increase in FFA and glycerol release from the eWAT of young *vs.* old infected-mice (Fig. 1I,J) and with FFA and glycerol accumulation in plasma from young *vs.* old infected-mice (Fig. 1K). This suggests that young mice mobilize FFA via a robust lipolytic response to MA-SARS-CoV-2 infection, while old mice fail to do so, consistent with age-dependent decline in adipocyte lipolysis^23–25^.

To test whether age-dependent decline in adipocyte lipolysis^23–25^ compromises disease tolerance to MA-SARS-CoV-2 infection, we generated adipocyte-specific *Pnpla2* (ATGL) deficient mice, *Pnpla2^AdipoQΔ/Δ^* mice. ATGL deletion was confirmed in eWAT, under steady state and after infection in young *Pnpla2^AdipoQΔ/Δ^ vs. Pnpla2^fl/fl^* mice (Fig. S3).

MA-SARS-CoV-2 infection was associated with a reduction of eWAT and ScWAT (subcutaneous WAT) mass in *Pnpla2^fl/fl^* mice but not in young *Pnpla2^AdipoQΔ/Δ^* mice (Fig. S4A,B). This was corroborated by failure to accumulate circulating FFA in young *Pnpla2^AdipoQΔ/Δ^ vs. Pnpla2^fl/fl^* infected mice (Fig. S4C,D). Moreover, infected *Pnpla2^AdipoQΔ/Δ^* mice exhibited lower circulating glycerol, compared to *Pnpla2^fl/fl^* mice (Fig. S4C,D). This confirms that ATGL is essential for the induction of adipose tissue lipolysis and FFA mobilization in response to SARS-CoV-2 infection.

Young *Pnpla2^AdipoQΔ/Δ^* mice developed severe disease (Fig. S5A,B) and succumbed (Fig. 1L,M) to MA-SARS-CoV-2 infection, while *Pnpla2^fl/fl^* controls remained minimally affected (Fig. S5A,B; 1L,M). Lung viral titers were similar between genotypes, as assessed across multiple timepoints (Fig. 1N). Moreover, viral titers remained below the detectable level in eWAT, as assessed 5 dpi (Fig. S5C,D). This confirming that young *Pnpla2^AdipoQΔ/Δ^* mice fail to establish disease tolerance to SARS-CoV-2 infection.

The lethal outcome of MA-SARS-CoV-2 infection in *Pnpla2^AdipoQΔ/Δ^*mice was attributed to ARDS, as illustrated histologically by prominent alveolar and interstitial infiltrates (Fig. 1O, S5E,F) and disruption of alveolar-capillary barrier (Fig. S5G, 1P). Critically, liver enzymes (ALT, AST) and kidney function (urea) remained normal in both genotypes (Fig. S5H-K), suggesting that lethality of MA-SARS-CoV-2 infection in young *Pnpla2^AdipoQΔ/Δ^*mice is not attributed to multi-organ failure.

Notably, influenza A virus (PR8 strain) virulence was not increased in young *Pnpla2^AdipoQΔ/Δ^ vs.* controls *Pnpla2^fl/fl^*mice, as assessed by disease severity, survival and body weight loss in response to an increasing range of infectious inocula (Fig. S6). This suggests that adipocyte lipolysis is dispensable to survive influenza.

Taken together these findings suggest that adipocyte lipolysis is essential to confer disease tolerance to SARS-CoV-2 infection. It also suggests that age-dependent decline in adipocyte lipolysis^23–25^ is sufficient to compromise disease tolerance to SARS-CoV-2 but not influenza A virus infection.

### Age-dependent impairment of adipocyte lipolysis is associated with COVID-19 severity

To validate whether age-dependent impairment of adipocyte lipolysis in mice impacts on COVID-19 severity, we performed untargeted plasma lipidomics in old *vs.* young, *Pnpla2^fl/fl^ vs. Pnpla2^AdipoQΔ/Δ^* mice infected with human specific SARS-CoV-2 B.1.351 (β) variant. The plasma lipidome (905 lipid metabolites) of old *vs.* young *Pnpla2^fl/fl^*mice revealed an age-dependent reduction of 124 lipid species (log₂FC ≥ 1, *p* ≤ 0.05) (Fig. 2A,B). This age dependent reduction was restricted to 45 lipid species in *Pnpla2^AdipoQΔ/Δ^*mice (Fig. 2C,D).

**Figure 2.**
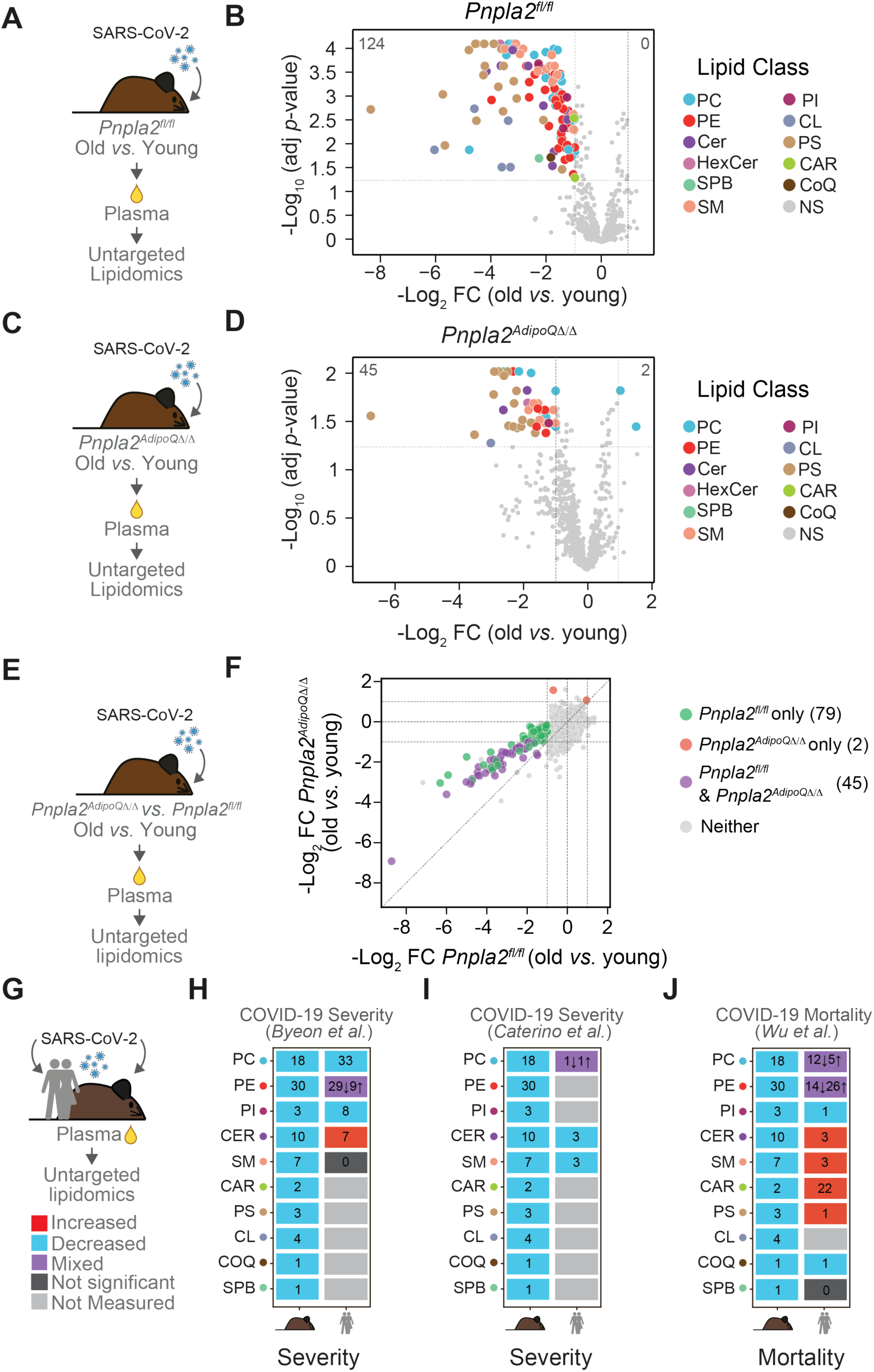
Age- and adipocyte lipolysis- dependent alterations in the plasma lipidome during SARS-CoV-2 infection are conserved in humans. (**A**) Young (3 months) and old (16 months) *Pnpla2^fl/fl^*mice were infected with the human specific SARS-CoV-2 B.1.351 (β) variant, plasma was collected at 3 dpi, and untargeted lipidomics was performed (n = 3-4 mice *per* group, males). Total lipids analyzed: 905. (**B**) Volcano plot showing differential lipid abundance in plasma from old *vs.* young *Pnpla2^fl/fl^* mice. Each dot represents an individual lipid species colored by lipid class. Numbers indicate regulated lipids in old mice. (**C**) Young and old *Pnpla2^AdipoQΔ/Δ^*mice were infected and analyzed as in (A). (**D**) Volcano plot showing differential lipid abundance in plasma from old *vs*. young *Pnpla2^AdipoQΔ/Δ^* mice. (**E**) Comparative analysis of age-dependent lipid changes between *Pnpla2^AdipoQΔ/Δ^*and *Pnpla2^fl/fl^* mice. (**F**) Scatter plot comparing log fold changes (old *vs*. young) between genotypes. Colored dots indicate lipids significantly altered in *Pnpla2^fl/fl^* mice (green, n=79), *Pnpla2^AdipoQΔ/Δ^* mice only (red, n=2), both genotypes (purple, n=45), or neither (grey). (**G**) Comparison of age-dependent and COVID-19-associated lipid changes between human^30–32^ and mouse plasma. (**H**) Heatmap comparing COVID-19 severity-associated lipid changes between humans^30^ and mice. (**I**) Heatmap showing conservation of COVID-19 severity-associated lipid changes between humans^31^ and mice. (**J**) Heatmap showing conservation of COVID-19 mortality-associated lipid changes between humans^32^ and mice. Numbers (H, I, J) indicate individual lipid species significantly altered within each lipid class; ↑ upregulated, ↓ downregulated. Colors (H, I, J) indicate regulation status: red, increased; blue, decreased; purple, mixed; dark gray, not significant; light gray, not measured. Abbreviations: PC, phosphatidylcholine; PE, phosphatidylethanolamine; Cer, ceramides; HexCer, hexosylceramide; SPB, sphingoid base; SM, sphingomyelin; PI, phosphatidylinositol; CL, cardiolipin; PS, phosphatidylserine; CAR, carnitine; CoQ, coenzyme Q; NS, not significant.

A direct comparison of the plasma lipidomic profiles from infected *Pnpla2^fl/fl^ vs. Pnpla2^AdipoQΔ/Δ^* mice revealed 79 lipid species reduced in an age- and adipocyte lipolysis-dependent manner (Fig. 2E,F). This suggests that defective adipocyte lipolysis accounts for ∼64% (79/124 lipid metabolites) of the age-dependent alterations in the plasma lipidome of SARS-CoV-2 infected mice.

We then compared the adipocyte-dependent lipidomic signature (79 lipids) of old infected mice with plasma lipidomic signatures associated with COVID-19 severity in three independent cohorts^30–32^. The adipocyte lipolysis-dependent reduction of phosphatidylcholine (PC), phosphatidylethanolamines (PE) and phosphatidylinositol (PI) in old infected mice was consistent with that of critical COVID-19 patients, reported by *Byeon et al.*^30^, where PE species showed mixed directionality (29↓9↑) (Fig. 2G,H). The reduction of PC, ceramide and sphingomyelin (SM) in old infected mice corroborated with that of severe COVID-19 patients, reported by *Caterino et al.*^31^, where PC (1↓1↑) displayed mixed directionality (Fig. 2G,I). The reduction of PC, PE, PI and coenzyme Q (COQ) was consistent with that of fatal COVID-19 cases reported by *Wu et al.*^32^, where PC (12↓5↑) and PE (14↓26↑) displayed mixed directionality (Fig. 2G,J). Data integration confirmed the age-dependent phospholipid depletion (PC, PE, PI) driven by impaired adipocyte lipolysis in mice as a conserved feature of COVID-19 severity and fatal outcomes in these human cohorts^30–32^, with ceramide and SM showing mixed directionality across datasets. This suggests that age-dependent compromise of adipocyte lipolysis contributes critically to explain the increase in COVID-19 severity and lethality in elderly individuals^8–10^.

Of note, severe COVID-19 is associated with an increase in circulating carnitines (CAR) ^32^, while old infected mice presented an adipocyte lipolysis-dependent reduction of CAR (Fig. 2G-J)^30–32^.

### Ketogenesis is dispensable to survive SARS-CoV-2 infection

We reasoned that impaired FFA mobilization may be responsible for the impaired ketogenesis associated with severe COVID-19^33^. In keeping with this notion, MA-SARS-CoV-2 infection induced the accumulation of circulating ketone bodies (Fig. S7A,B)^33^ in an adipocyte lipolysis-dependent manner as revealed by the failure of infected *Pnpla2^AdipoQΔ/Δ^* mice to accumulate ketone bodies in the circulation in response to infection, when compared to control infected *Pnpla2^fl/fl^* mice (Fig. S7A,B).

To address whether the protective effect of adipocyte lipolysis is mediated via a mechanism that relies on ketogenesis, we generated *Hmgcs2^AlbΔ/Δ^* mice carrying a deletion of 3-hydroxy-3-methylglutaryl-CoA synthase 2 (*Hmgcs2*), the rate limiting mitochondrial ketogenesis enzyme^34^, specifically in hepatocytes. As expected, *Hmgcs2^AlbΔ/Δ^* mice failed to accumulate circulating ketone bodies in response to MA-SARS-CoV-2 infection (Fig. S7C,D). However, this was not associated with increased disease severity (Fig. S7E), mortality (Fig. S7F), body weight loss (Fig. S7G) or thermoregulation (Fig. S7H), compared to control *Hmgcs2^fl/fl^* mice. Together, these findings suggest that hepatic ketogenesis is dispensable to survive MA-SARS-CoV-2 infection.

### Adipocyte lipolysis regulates monocyte lung recruitment in SARS-CoV-2 infection

We next explored other possible mechanisms whereby adipose tissue lipolysis counters the pathogenesis of ARDS to confer disease tolerance to MA-SARS-CoV-2 infection. Flow cytometry analysis of lung immune cells revealed that infected *Pnpla2^AdipoQΔ/Δ^*mice exhibited marked reductions in both Ly6C^+^MHCII^+^ and Ly6C^+^MHCII^−^ monocyte populations, compared to control *Pnpla2^fl/fl^* mice, as assessed at 5 dpi (Fig. 3A-C, S8A,B). While alveolar macrophages (AM; F4/80^+^CD11c^+^CD11b^−^) were reduced upon infection, there was no difference between genotypes (Fig. S8C,D). Similarly, interstitial macrophages (IM; F4/80^+^CD11c^low^CD11b^+^), neutrophils (CD11b^+^Ly6G^+^) and dendritic cells (DCs; CD11c^+^MHCII^+^) showed comparable numbers between genotypes under both control and infected conditions (Fig S8E-I). Although infection led to an increase in NK cells (CD3⁻NK1.1⁺) and NKT cells (CD3⁺NK1.1⁺), no significant differences were observed between genotypes under both control and infected conditions (Fig. S8J-M). This was also the case for CD8^+^ T cells (CD3^+^CD8^+^) and CD4^+^ T cells (CD3^+^CD4^+^) (Fig. S9A-D), including activated CD8^+^ T cells (CD8^+^CD44^+^) (Fig. S9E,F) and activated CD4^+^ T cells (CD4^+^CD44^+^) (Fig. S9G,H), as well as for total B cells (CD11b^−^CD19^+^) (Fig. S9I,J). This suggests that adipose tissue lipolysis selectively supports the recruitment of monocytes into the lungs without affecting other immune populations.

**Figure 3.**
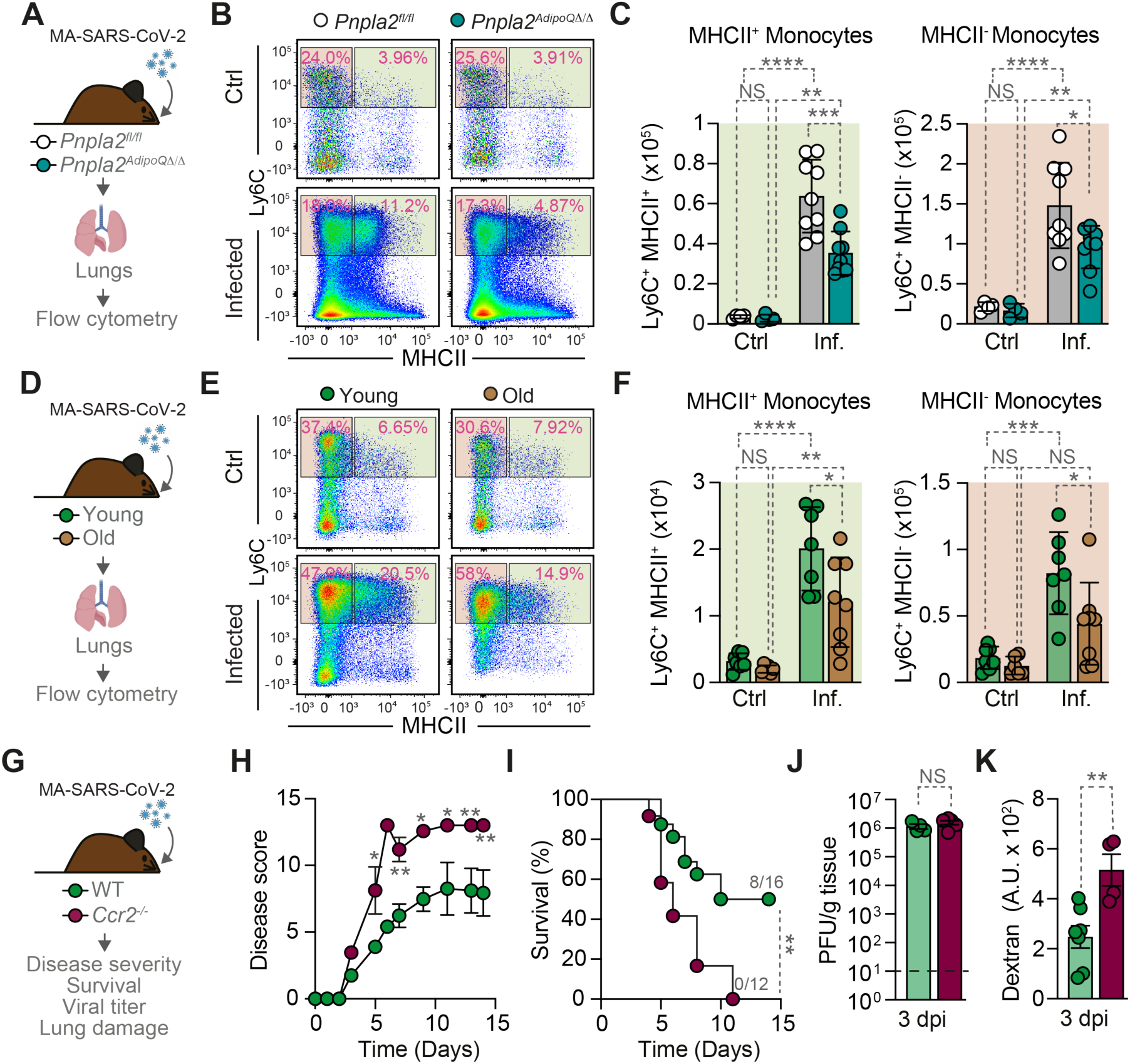
Adipose tissue lipolysis regulates monocyte lung infiltration during MA-SARS-CoV-2 infection. (**A**) Young *Pnpla2^fl/fl^* and *Pnpla2^AdipoQΔ/Δ^* mice were infected with MA-SARS-CoV-2 and lung monocyte populations were analyzed by flow cytometry at 5 dpi. (**B**) Representative flow cytometry plots for Ly6C^+^MHCII^+^ and MHCII^−^ monocyte subsets from control (Ctrl) and infected (Inf.) *Pnpla2^fl/fl^ vs. Pnpla2^AdipoQΔ/Δ^* mice. (**C**) Quantification of lung Ly6C^+^MHCII^+^ and MHCII^−^ monocytes from (B) (n=4-9 mice *per* group, males & females). (**D**) Young (10-16 weeks) and old (>20 months) C57BL/6J mice infected with MA-SARS-CoV-2 analyzed by flow cytometry at 5 dpi for lung monocyte populations. (**E**) Representative flow cytometry plots showing Ly6C^+^MHCII^+^ and MHCII^−^monocyte. (**F**) Quantification of Ly6C^+^MHCII^+^ and MHCII^−^ monocyte from (E) (n=6-8 mice *per* group, males). (**G**) Wild-type (WT) and *Ccr2^−/-^* mice infected with MA-SARS-CoV-2 and monitored for (**H**) disease severity and (**I**) survival (n=12-16 mice *per* group, males & females). (**J**) Lung viral titers at 3 dpi (n=4-5 mice *per* group, males & females). Dashed line indicates limit of detection. (**K**) Rhodamine fluorescence intensity at 3 dpi (n=4-7 mice *per* group, males and females). Data represented as mean *±* SD (C, F) or SEM (H, J, K). Numbers indicate surviving mice *per* total mice at day 14 (I). Circles represent individual mice (C, F, I, J, K). Survival curves were compared using the Cox proportional hazards model adjusted for experiment, with pairwise log-rank tests between-group comparisons (I). *p* values determined using Log-rank test (I) and two-way ANOVA (C, F, H) or unpaired t-test (J, K). NS, not significant; *p < 0.05; **p < 0.01; ***p < 0.001; ****p < 0.0001. Data pooled from one (J), two (A-F, K) or six (H, I) independent experiments with a similar trend.

The impaired recruitment of Ly6C^+^MHCII^+^ and Ly6C^+^MHCII^−^ monocytes to the lungs was also a hallmark of the severe outcome of MA-SARS-CoV-2 infection in old *vs.* young mice at 5 dpi (Fig. 3D-F, S10A,B). AM were reduced in old *vs.* young mice at steady state but remained equivalent upon infection in both age groups (Fig. S10C,D). IM showed a marked reduction in old *vs.* young mice after infection (Fig. S10E). Neutrophils, DCs, and NKT cells showed largely comparable numbers between young and old mice (Fig. S10F-I,L,M). NK cells showed a modest infection-induced increase in young but not old mice (Fig. S10J,K). This suggests that while some immune populations show minor age-related differences, the primary defect in old mice is impairment in monocyte lung recruitment (Fig. 3E,F). This indicates that monocyte recruitment might be involved in establishing disease tolerance to MA-SARS-CoV-2 infection.

### Monocytes recruitment to the lung is essential to survive SARS-CoV-2 infection

To probe whether monocyte lung recruitment is necessary for the establishment of disease tolerance to SARS-CoV-2 infection, we used young adult CCL2 chemokine receptor-deficient (*Ccr2^−/-^*) mice, which fail to mobilize monocytes from bone marrow (BM) into peripheral tissue^35^. *Ccr2^−/-^*mice developed severe disease (Fig. 3G,H) and succumbed to MA-SARS-CoV-2 infection, compared to wild-type controls (Fig. 3I), as previously shown^36^. Similar viral titers between genotypes (Fig. 3J) confirms that monocyte lung recruitment is essential to establish disease tolerance to SARS-CoV-2 infection.

The lethal outcome of MA-SARS-CoV-2 infection in *Ccr2^−/-^*mice was associated with a near-complete absence of lung MHCII^+^ and MHCII^−^ monocytes (Ly6C^+^), compared to wild-type littermate controls (Fig. S11A-D). The lethal outcome of SARS-CoV-2 infection in *Ccr2^−/-^* mice was characterized by the development of ARDS (Fig. 3K), showing that monocyte lung recruitment establishes disease tolerance to SARS-CoV-2 infection via a mechanism that prevents the development of ARDS.

Of note, the number of lung AM were comparable (Fig. S11E,F) while IM showed a modest decrease in absolute numbers (Fig. S11G). Neutrophil lung infiltration was exacerbated in infected *Ccr2^−/-^ vs.* wild-type controls (Fig. S11H,I). Dendritic cells showed a modest decrease in both percentage and absolute number (Fig. S11J,K), while NK cells and NKT cells remained comparable between genotypes (Fig. S11L-O).

To test whether monocyte recruitment into the lungs is sufficient *per se* to counter the pathogenesis of ARDS, we adoptively transferred bone marrow-derived monocytes (BMDMo) intranasally to young *Pnpla2^AdipoQΔ/Δ^*mice (Fig. 4A). To trace BMDMo, we used *LysM^Cre/WT^ R26^tdT/WT^* (*tdT^LysM^*) mice^37^, in which myeloid-lineage specific Cre (*LysM^Cre/WT^*) induces the expression of a *Rosa26*-Flox-stop-Flox tandem dimer (td) Tomato allele (*R26^tdT/WT^*) (Fig. S12A,B). Adoptive transfer of BMDMo 3, 5 and 7 dpi rescued *Pnpla2^AdipoQΔ/Δ^*mice, preventing disease progression (Fig. 4B) and lethality to 33%, compared to 84% lethality in vehicle-treated controls (Fig. 4C, S12C-E). This shows when naïve monocytes access the lung, these can prevent the onset of lethal ARDS caused by SARS-CoV-2 infection.

**Figure 4.**
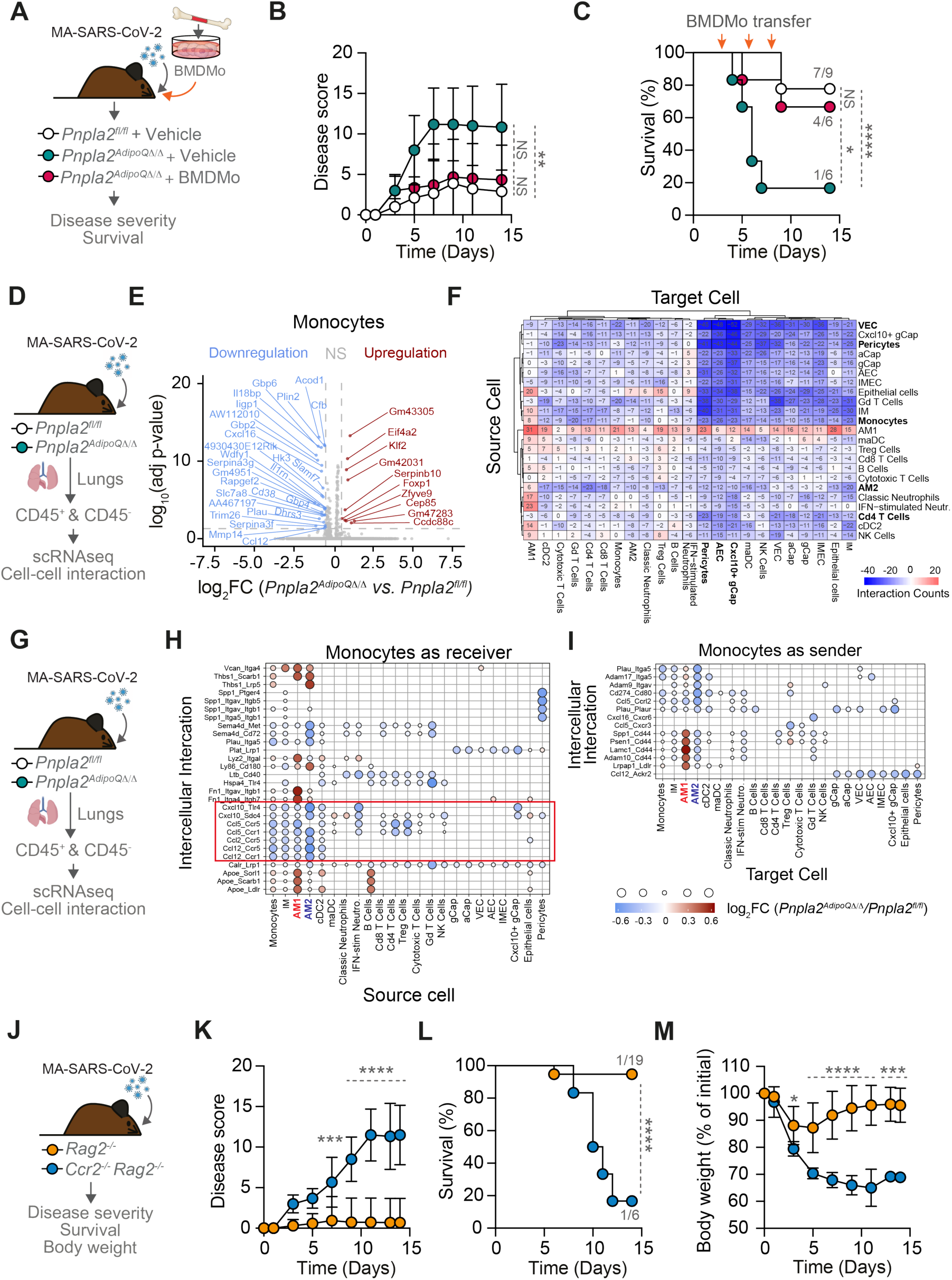
Monocyte lung infiltration is essential to establish disease tolerance to MA-SARS-CoV-2 infection. (**A**) *Pnpla2^fl/fl^*and *Pnpla2^AdipoQΔ/Δ^* mice were infected with MA-SARS-CoV-2. *Pnpla2^AdipoQΔ/Δ^* mice received either vehicle or adoptive transfer of BM derived monocytes (BMDMo) from C57BL6/J mice and monitored for (**B**) disease severity and (**C**) Survival (n=6-9 mice *per* group, males). (**D**) Young *Pnpla2^fl/fl^* and *Pnpla2^AdipoQΔ/Δ^* mice were infected with MA-SARS-CoV-2 and CD45^+^ and CD45^−^ populations were harvested at 5 dpi for scRNA-seq analysis and cell-cell interaction (n=3 mice *per* group, males). (**E**) Volcano plot showing differential gene expression (log₂FC ≥ 0.5, *p*_adj_ ≤ 0.05) in monocytes from *Pnpla2^AdipoQΔ/Δ^ vs. Pnpla2^fl/fl^* infected mice. Differentially expressed genes (DEG) are highlighted in red (upregulated) or blue (downregulated), non-DEG (NS) are shown in grey. Each dot represents an individual gene. (**F**) Heatmap showing differential ligand-receptor interaction counts between structural and immune lung populations from *Pnpla2^AdipoQΔ/Δ^ vs. Pnpla2^fl/fl^* infected mice. Blue indicates decreased and red indicates increased interaction counts in *Pnpla2^AdipoQΔ/Δ^ vs. Pnpla2^fl/fl^*infected mice. (**G)** CD45^+^ and CD45^−^ populations from infected *Pnpla2^fl/fl^* and *Pnpla2^AdipoQΔ/Δ^* mice were analyzed for cell-cell interaction as in (D). **(H)** Dot plot showing differential ligand-receptor interactions with monocytes as receivers (express receptor) and **(I)** as senders (express ligand) across all cell types from *Pnpla2^AdipoQΔ/Δ^ vs. Pnpla2^fl/fl^* infected mice. Red box (H) highlights reduced chemokine signaling. Only statistically significant interactions (*p* < 0.01) are shown (H, I). Dot color and size (H, I) indicate the log_2_FC in ligand-receptor mean expression score (lr.mean) between *Pnpla2^AdipoQΔ/Δ^* and *Pnpla2^fl/fl^* interaction strength (red: increased; blue: decreased in *Pnpla2^AdipoQΔ/Δ^*). (**J**) *Rag2^−/-^* and *Ccr2^−/-^ Rag2^−/-^* mice infected with MA-SARS-CoV-2 and monitored for (**K**) disease severity, (**L**) survival and (**M**) body weight loss (n=6-19 mice *per* group, males & females). Data represented as mean *±* SD (B, K, M). Numbers indicate surviving mice *per* total mice at day 14 (C, L). Survival curves were compared using the Cox proportional hazards model adjusted for experiment, with pairwise log-rank tests between-group comparisons (C, L). *p* values determined using Log-rank test (C, L) or two-way ANOVA (B, K, M). NS, not significant; *p < 0.05; **p < 0.01; ***p < 0.001; ****p < 0.0001. Data from one (E-I), two (B, C), or five (K-M) independent experiments with a similar trend.

### Adipocyte lipolysis regulates lung gene expression during MA-SARS-CoV-2 infection

We then explored the mechanism whereby monocytes prevent the onset of lethal ARDS caused by SARS-CoV-2 infection. To do so, we performed single-cell RNA sequencing (scRNAseq) of CD45^+^ and CD45^−^ fractions from lungs of infected *Pnpla2^fl/fl^* and *Pnpla2^AdipoQΔ/Δ^* mice at 5 dpi. Analysis of CD45^+^ scRNAseq, focusing on the myeloid cells, identified monocytes, IM, AM (AM1 and AM2), DCs (cDC1, cDC2, Mo-DC, and pDC), and neutrophils [classical and interferon (IFN)-stimulated] visualized by uniform manifold approximation and projection (UMAP) (Fig. S13A,B).

Differential gene expression (DGE) analysis revealed predominant downregulation of genes in monocyte clusters from *Pnpla2^AdipoQΔ/Δ^ vs. Pnpla2^fl/fl^* infected mice (log₂FC ≥ 0.5, *p*_adj_ ≤ 0.05). These included IFN-stimulated genes (ISG; *Gbp2, Gbp4, Gbp6, Iigp1, Il18bp, Wdfy1, Gm4951*) (Fig. 4D,E), consistent with impairment of IFN signaling in monocytes from severe COVID-19 patients^38^. Critically, *Acod1* (aconitate decarboxylase 1), which produces itaconate, an immunometabolite with potent tissue-protective properties^39,40^, was among the most significantly downregulated genes (Fig. 4E). Minor changes were also observed in classic and IFN-stimulated neutrophils (Fig. S13C) and IM (Fig. S13D), while no DGE was detected in AM (Fig. S13E) and DCs (Fig. S13F). Of note, there was a negligible DGE in adaptive immune cells between *Pnpla2^AdipoQΔ/Δ^* and *Pnpla2^fl/fl^*mice, essentially absent in CD8^+^ cytotoxic T cells, CD4^+^ T helper cells CD8^+^ cytotoxic T cells and regulatory T cells (Fig S13G-I). This monocyte-specific transcriptional signature suggests that adipocyte lipolysis controls monocyte recruitment and/or effector function, underlying ARDS in severe COVID-19^41^.

### Adipocyte lipolysis regulates lung cell-cell communication during MA-SARS-CoV-2 infection

To determine whether adipocyte lipolysis regulates intercellular lung communication, we combined scRNAseq analysis (LIANA)^42^ from sorted lung CD45^+^ and CD45^−^ 5 dpi. Intercellular communication counts were reduced in *Pnpla2^AdipoQΔ/Δ^ vs. Pnpla2^fl/fl^*mice (Fig. 4F). This was particularly pronounced for structural cell populations, including vascular endothelial cells (VEC), capillary endothelial cells (Cap), pericytes, and epithelial cells (Fig. 4F). Of note there was a marked reduction of intercellular interactions among endothelial cells and among epithelial cells (Fig 4F), consistent with the development of ARDS in *Pnpla2^AdipoQΔ/Δ^* infected mice.

Of 48,508 putative ligand-receptor interactions analyzed, 309 showed significant alterations (log₂FC ≥ 0.5, *p* ≤ 0.01). These included ligand-receptor interactions associated with tissue damage control (173 reduced *vs.* 136 increased) as well as impaired AM interaction with epithelial and endothelial cells via oncostatin M, associated with loss of lung epithelial barrier integrity^43^ (Table S1).

Focusing on monocytes, the reduction in ligand-receptor interactions was bidirectional (Fig. 4G-I). There was a reduced interaction strength of chemokine signals received by lung-infiltrating monocytes from AM and IM in *Pnpla2^AdipoQΔ/Δ^ vs. Pnpla2^fl/fl^* mice (Fig. 4H). These monocytes also showed diminished relay of signals to cell populations predicted to regulate tissue remodeling, immune regulation, and inflammatory resolution (Fig. 4I). This is illustrated by reduced urokinase (uPA/*Plau*) signaling to its receptor uPAR (*Plaur*) on Cxcl10⁺ gCap (general capillary) endothelial cells (Fig. 4I), consistent with impaired extracellular-matrix remodeling and vascular repair in the infected lung^44,45^. In contrast, AM1 showed increased interaction strength with incoming monocytes in *Pnpla2^AdipoQΔ/Δ^ vs. Pnpla2^fl/fl^* infected mice, notably through pro-inflammatory (e.g. *Il1b*, *Tnf*, *Cxcl10*) and CD44-mediated ligand-receptor pairs (Fig. 4I). Together, these findings suggest that adipocyte lipolysis sustains the capacity of lung-infiltrating monocytes to establish intercellular communication with parenchyma cells, presumably supporting tissue repair and inflammatory resolution during SARS-CoV-2 infection.

### Monocytes establish disease tolerance to SARS-CoV2 infection independently of adaptive immunity

Adaptive immune responses can aggravate SARS-CoV-2 immunopathology, as demonstrated in *Rag*-deficient mice lacking mature T and B cells^46^. To test whether monocytes confer protection against SARS-CoV-2 infection via a mechanism that regulates adaptive immunity, we generated *Ccr2*^−/-^*Rag2*^−/-^ mice. Consistent with previous findings^46^, *Rag2*^−/-^ mice showed residual disease severity (Fig. 4J,K) and mortality (Fig. 4L) preserving body weight (Fig. 4M). In contrast, *Ccr2*^−/-^*Rag2*^−/-^ mice developed severe disease (Fig. 4K) and succumbed (Fig. 4L) to MA-SARS-CoV-2 infection with pronounced body weight loss (Fig. 4M). This demonstrated that the protective effect of monocytes against SARS-CoV-2 infection acts independently of adaptive immunity.

### Adipose tissue lipolysis supports emergency myelopoiesis in SARS-CoV2 infection

We reasoned that failure of *Pnpla2^AdipoQΔ/Δ^* mice to recruit monocytes to the lungs in response to SARS-CoV-2 infection may be caused by a defect in emergency myelopoiesis^47,48^. In support of this notion, MA-SARS-CoV-2 infection induced BM emergency myelopoiesis in *Pnpla2^fl/fl^* mice, which was impaired in *Pnpla2^AdipoQΔ/Δ^*mice (Fig. 5A,B). This is revealed by a marked reduction of long-term hematopoietic stem cells (LT-HSC), short-term HSC (ST-HSC), multipotent progenitor 3 (MPP3), common myeloid progenitors (CMP), bulk granulocyte-monocyte progenitors (GMP), and GMP numbers in infected *Pnpla2^AdipoQΔ/Δ^ vs. Pnpla2^fl/fl^* mice (Fig. 5A,B). Of note, most individual stem cell populations involved in hematopoiesis and myelopoiesis remained intact during MA-SARS-CoV-2 infection (Fig. S14, S15). This reveals a functional adipocyte lipolysis-dependent mechanism supporting emergency myelopoiesis during COVID-19^41^.

**Figure 5.**
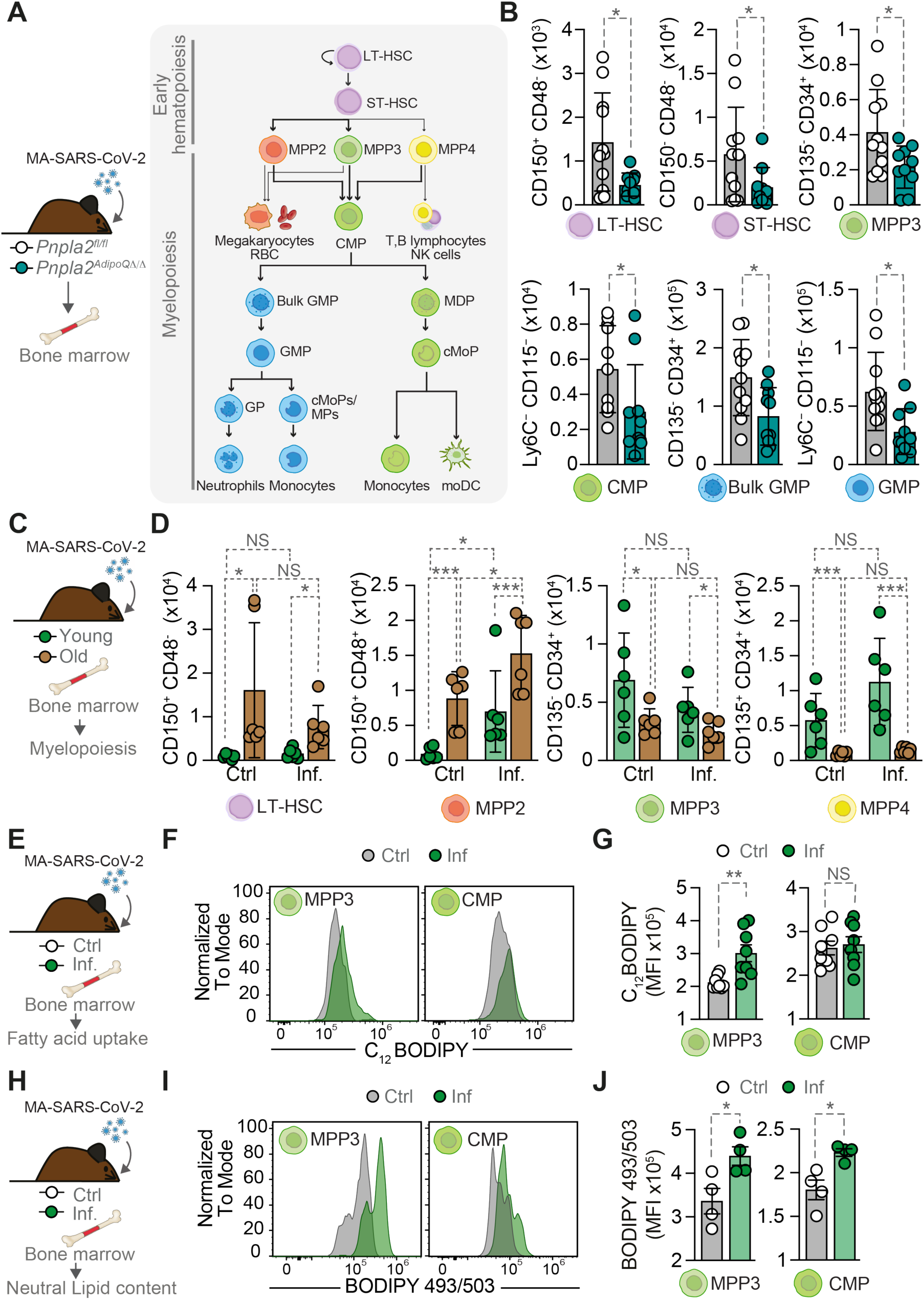
Adipose tissue lipolysis regulates bone marrow hematopoiesis during MA-SARS-CoV-2 infection. (**A**) Young *Pnpla2^fl/fl^* and *Pnpla2^AdipoQΔ/Δ^* mice infected with MA-SARS-CoV-2 were monitored for BM hematopoietic stem and progenitor cells (HSPCs) and myeloid progenitors by flow cytometry at 5 dpi. LT-HSC: long-term hematopoietic stem cells; ST-HSC: short-term hematopoietic stem cells; MPP: multipotent progenitors; CMP: common myeloid progenitors; GMP: granulocyte-monocyte progenitors; MDP: monocyte-dendritic cell progenitors; cMoP: common monocyte progenitors, GP: Granulocyte progenitors; MPs (monocyte progenitor), moDCs (monocyte-derived dendritic cells). (**B**) Quantification of LT-HSC, ST-HSC, MPP3, CMP, Bulk GMP, and GMP populations in BM from *Pnpla2^fl/fl^*and *Pnpla2^AdipoQΔ/Δ^* mice (n=10-11 mice *per* group, males & females). (**C**) Young (10-16 weeks) and old (>20 months) C57BL/6J mice were infected with MA-SARS-CoV-2 and BM myelopoiesis was analyzed at 5 dpi. (**D**) Quantification of LT-HSC, MPP2, MPP3, and MPP4 populations in BM from young *vs*. old control (Ctrl) and infected (Inf.) mice (n=6 mice *per* group, males). (**E**) Control and infected C57BL/6J young mice were analyzed for FFA uptake by BM progenitor cells using BODIPY C_12_ at 5 dpi. (**F**) Representative flow cytometry histograms showing BODIPY C_12_ staining (fatty acid uptake) in MPP3 and CMP populations from control and infected mice. (**G)** Quantification of BODIPY C_12_ mean fluorescence intensity (MFI) from (F) (n=8 mice *per* group, males). (**H**) Control and infected C57BL/6J young mice were analyzed for lipid content in BM progenitor cells using BODIPY 493/503 at 5 dpi. (**I**) Representative flow cytometry histograms showing BODIPY 493/503 staining (lipid content) in MPP3 and CMP populations from control and infected mice. **(J)** Quantification of BODIPY 493/503 MFI from (I) (n=4 mice *per* group from one out of two independent experiments with similar trend, males). Data represented as mean *±* SD (B, D) or SEM (G, J). Circles represent individual mice (B, D, G, J). *p* values were determined using unpaired t-test (B, G, J) and two-way ANOVA (D). NS, not significant; *p < 0.05; **p < 0.01; ***p < 0.001. Data pooled from three (B) or two (D, G) independent experiments with a similar trend or from one experiment representative of two independent experiments (J).

Old mice also presented clear signs for impaired BM myelopoiesis at steady state (Fig. 5C,D, S16, S17) as well as emergency myelopoiesis in response to MA-SARS-CoV-2 infection (Fig. 5C,D). This is revealed by baseline changes in hematopoietic progenitor cells across LT-HSC and MPP populations (Fig. 5D, S16), consistent with previously described^49–51^. Upon MA-SARS-CoV-2 infection, old mice presented a reduction of bone marrow MPP3 and MPP4 cells, compared to the young, infected mice (Fig. 5D), despite a baseline myeloid bias^51^. This suggest that age-dependent impaired adipocyte lipolysis compromises MA-SARS-CoV-2 infection-induced emergency myelopoiesis in old mice. Of note, most individual stem cell populations remained intact during MA-SARS-CoV-2 infection in old *vs.* young C57BL/6J mice (Fig. S16, S17).

Consistent with adipocyte lipolysis-derived FFA fueling myelopoiesis, MA-SARS-CoV-2 infection increased intracellular FFA (BODIPY™ FL C_12_) uptake in a subset of hematopoietic progenitors including LKS, ST-HSC (Fig. S18) and myeloid progenitor MPP3 but not in others, such as the downstream progenitor CMP (Fig. 5E-G, S18). Moreover, MA-SARS-CoV-2 infection was also associated with an accumulation of neutral lipids (BODIPY 493/503) in most of the hematopoietic and myeloid progenitor including both MPP3 and CMP populations (Fig. 5H-J, S19).

Taken together these findings suggest that FFA mobilization via adipocyte lipolysis provides the essential metabolic fuel to support BM stress myelopoiesis and drive the expansion of myeloid progenitors including MPP3 and CMP. This metabolic circuit fails in old mice impairing the induction of BM stress myelopoiesis in response to SARS-CoV-2 infection.

### FFA uptake/oxidation is required for disease tolerance to MA-SARS-CoV-2 infection

To test whether adipocyte lipolysis delivers FFA to hematopoietic cells, FFA cellular uptake via CD36 was inhibited pharmacologically (sulfo-N-succinimidyl oleate; SSO). CD36 inhibition compromised survival to MA-SARS-CoV-2 infection, increasing mortality to 67% *vs.* 25% in SSO *vs.* vehicle-treated controls (Fig. 6A,B), without affecting viral titer in the lungs (Fig. 6C). This was associated with a marked reduction in monocyte lung recruitment (Fig. 6D,E), without affecting AM, IM, neutrophils, DCs, NK cells, and NKT cells (Fig. S20). This is consistent with a selective impairment of monocyte recruitment similar to infected *Pnpla2^AdipoQΔ/Δ^* mice and old mice (Fig. 3C and F).

**Figure 6.**
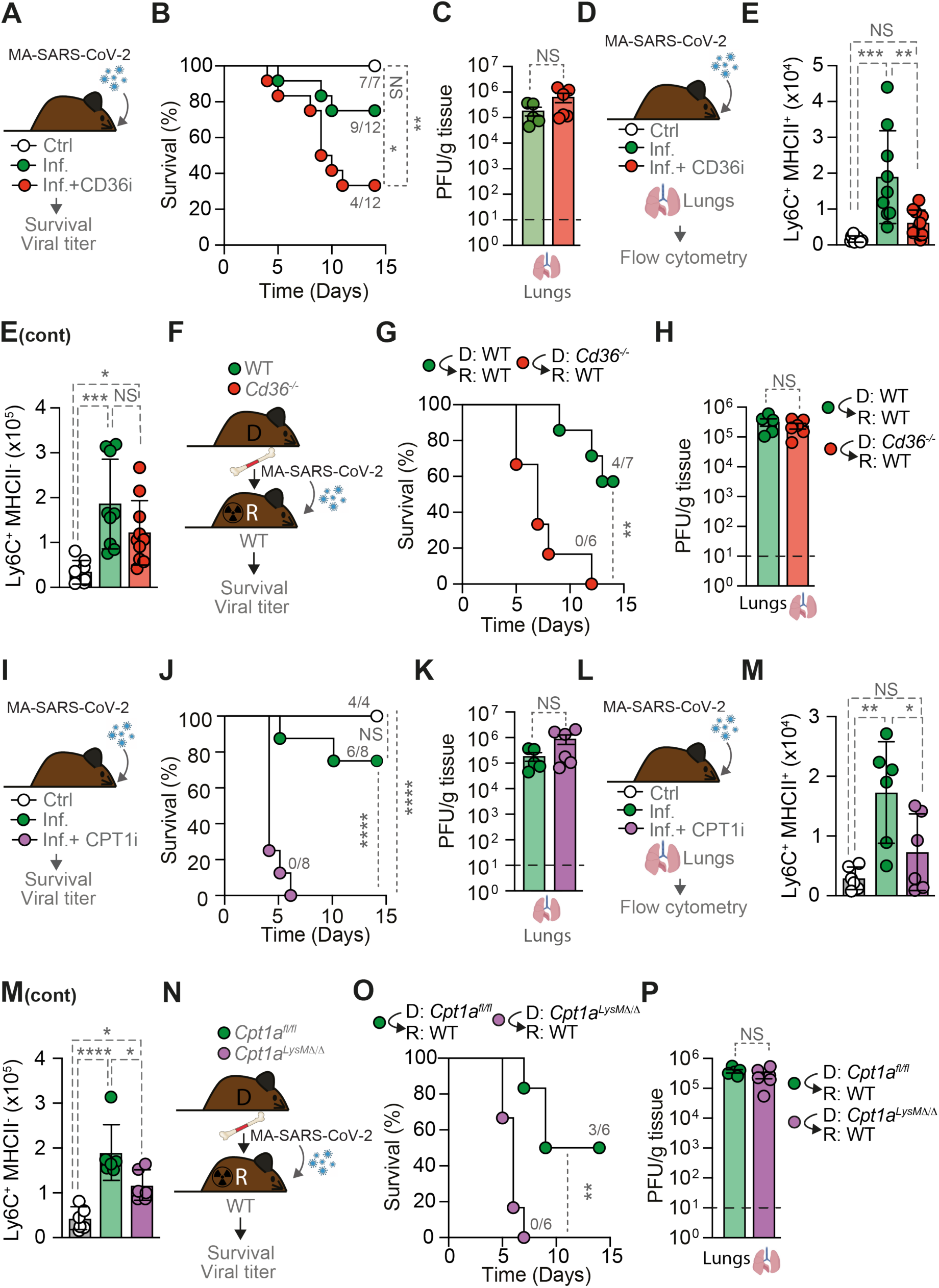
FFA uptake and oxidation are required for disease tolerance to MA-SARS-CoV-2 infection. (**A**) C57BL/6J mice were infected with MA-SARS-CoV-2 and treated with vehicle or CD36 inhibitor, sulfo-N-succinimidyl oleate, (CD36i), and monitored for (**B**) Survival (n=7-12 mice *per* group, males) and (**C**) Lung viral titers (PFU/g tissue) at 3 dpi (n=5-6 mice *per* group, males). (**D**) C57BL/6J mice were infected and treated as in (A) and lung monocyte populations were analyzed by flow cytometry at 3 dpi. Quantification of (**E**) Ly6C^+^ MHCII^+^ and Ly6C^+^ MHCII^−^ monocytes (n=8-10 mice *per* group, males). (**F**) Lethally irradiated C57BL/6J recipient (R) mice were transplanted with BM from WT or *Cd36^−/-^* donor (D) mice, and mice were allowed to reconstitute for 6-8 weeks before being infected with MA-SARS-CoV-2. (**G**) Survival curves (n=6-7 mice *per* group, males & females) and (**H**) lung viral titers (n=5-6 mice *per* group, males & females). (**I**) C57BL/6J mice were infected with MA-SARS-CoV-2 and treated with vehicle or CPT1 inhibitor, etomoxir (CPT1i), and monitored for (**J**) Survival (n=4-8 mice *per* group, males) and (**K**) lung viral titers (n=5-6 mice *per* group, males). (**L**) C57BL/6J mice were infected and treated as in (J) and lung monocyte populations were analyzed by flow cytometry at 3 dpi. (**M**) Quantification of Ly6C^+^ MHCII^+^ and Ly6C^+^ MHCII^−^ monocytes (n=6 mice *per* group, males). (**N**) Lethally irradiated C57BL/6J recipient (R) mice were transplanted with BM from *Cpt1a^fl/fl^*or *Cpt1a^LysMΔ/^*^Δ^ donor (D) mice and infected with MA-SARS-CoV-2 (6-8 weeks thereafter). (**O**) Survival curves (n=6 mice *per* group, males & females) and (**P**) lung viral titers of bone marrow chimeras at 5 dpi (n=5-6 mice *per* group, males & females). Dashed line indicates limit of detection (C, H, K, P). Data represented as mean *±* SD (E, M) or SEM (C, H, K, P). Circles represent individual mice (B, C, E, G, H, J, K, M, O, P). Numbers in B, G, J and O indicate surviving mice *per* total mice at day 14. *p* values determined using Log-rank test (B, G, J, O), unpaired t-test (C, H, K, P) and one-way ANOVA (E, M). NS, not significant; *p < 0.05; **p < 0.01; ****p < 0.0001. Data pooled from two (C, G, H, J, K, M, O, P) or three (B, E) independent experiments with a similar trend.

To probe the involvement of CD36 genetically, lethally irradiated wild-type mice received BM from wild-type (*Cd36^+/+^*) *vs. Cd36-*deficient (*Cd36^−/-^*) mice (Fig. 6F). All bone marrow chimeric mice carrying *Cd36^−/-^* hematopoietic cells succumbed to MA-SARS-CoV-2 (100%), compared to 40% lethality in control bone marrow chimeric mice carrying *Cd36^+/+^* hematopoietic cells (Fig. 6G). This was not associated with changes in viral titers (Fig. 6H), suggesting that hematopoietic CD36 expression is essential to establish disease tolerance to MA-SARS-CoV-2 infection.

To test whether adipocyte lipolysis is required to support hematopoietic β-oxidation during MA-SARS-CoV-2 infection, mitochondrial FFA import by carnitine palmitoyltransferase-1 (CPT1)^52^ was inhibited pharmacologically (etomoxir) (Fig. 6I). CPT1 inhibition compromised survival to MA-SARS-CoV-2 infection, increasing mortality to 100% *vs.* 25% in Etomoxir *vs.* vehicle-treated controls (Fig. 6J), without affecting viral titers in the lungs (Fig. 6K). Mortality was associated with a marked reduction in monocyte lung recruitment (Fig. 6L,M), without affecting major lung immune cell populations (Fig. S21). This suggests that the selective impairment of monocyte recruitment in infected *Pnpla2^AdipoQΔ/Δ^* mice and old mice (Fig. 3C and F) is due to failure of adipocyte lipolysis to deliver FFA required to support β-oxidation.

MA-SARS-CoV-2 infection was associated with a marked increase of protein synthesis across most hematopoietic stem cell populations, including myeloid progenitors such as MPP3 and CMP (Fig. S22), as assessed *ex vivo* by puromycin incorporation^53^. This was fully inhibited by etomoxir, suggesting that stress myelopoiesis relies on β-oxidation, explaining why inhibition of β-oxidation compromises survival to MA-SARS-CoV-2 infection.

To probe the involvement of CPT1 genetically, we used *Cpt1a^LysMΔ/Δ^*mice carrying a myeloid-specific deletion of CPT1A, the predominant CPT1 isoform expressed in myeloid cells, to generate bone marrow chimeric mice carrying hematopoietic cells from *Cpt1a^LysMΔ/Δ^ vs.* control (*Cpt1a^fl/fl^*) mice (Fig. 6N). All bone marrow chimeric mice carrying *Cpt1a^LysMΔ/Δ^* hematopoietic cells succumbed to MA-SARS-CoV-2 (100%), compared to 50% lethality in control bone marrow chimeric mice carrying *Cpt1a^fl/fl^* hematopoietic cells (Fig. 6O). This was not associated with changes in viral titers (Fig. 6P), suggesting that myeloid CPT1A expression is essential to establish disease tolerance to MA-SARS-CoV-2 infection. Together, these results establish that both CD36-mediated FFA uptake and CPT1A-dependent mitochondrial β-oxidation are required to confer disease tolerance to SARS-CoV-2 infection.

## DISCUSSION

We identified adipose tissue lipolysis as a rate-limiting metabolic requirement for emergency myelopoiesis that confers disease tolerance to SARS-CoV-2 infection. However, this protective effect is not required to establish disease tolerance to influenza virus infection, illustrating the distinct pathology caused by these two respiratory viruses^54,55^.

Mechanistically, FFA mobilization via adipocyte lipolysis fuels the expansion and functional maturation of monocytes through myelopoiesis that protects from ARDS, the primary cause of COVID-19 mortality^8^. This protective inter-organ communication circuit fails with aging: old mice cannot mount the lipolytic response required to sustain emergency myelopoiesis, lowering monocyte production and lung recruitment, leading to lethal ARDS. These findings establish a metabolic-immune based tissue damage control mechanism^3,4^ that protects against viral ARDS and its age-dependent collapse as a mechanism of COVID-19 susceptibility in elderly individuals^8–10^.

The central role of monocytes in the protective response to SARS-CoV-2 infection is consistent with their capacity to support tissue function^37^ and disease tolerance to SARS-CoV-2 infection^56,57^. Importantly, this does not preclude aberrantly activated monocyte-derived macrophages from driving COVID-19 severity^41,58,59^, a duality reflecting how metabolic heterogeneity impacts on macrophage effector functions^58,60,61^.

FFA deprivation resulting from impaired adipocyte lipolysis due to aging or imposed genetically in young mice creates a pathological cascade that compromises the intercellular network supporting lung function during SARS-CoV-2 infection. This is illustrated for example by impaired signaling of alveolar macrophages with epithelial and endothelial cells (*i.e., via* oncostatin M and CXCL10), predictive of loss of lung epithelial barrier integrity^43^. Loss of these tissue damage control mechanisms^3^ compromises lung function and therefore disease tolerance to SARS-CoV-2 infection.

The dual requirement for CD36 and CPT1A establishes mitochondrial β-oxidation of extracellular FFA, rather than that of endogenous fatty acids alone, as the rate-limiting metabolic program enabling monocytes to execute tissue-protective functions in infected lungs. Whether FFA β-oxidation represents a conserved metabolic axis governing disease tolerance across diverse infections remains to be established.

A critical role of adipose tissue lipolysis in the support of emergency myelopoiesis during SARS-CoV-2 infection resembles what has been described in myocardial infarction^62^. Whether, similar to myocardial infarction^62^, bone marrow adipose tissue lipolysis is the source of FFA supporting emergency myelopoiesis in SARS-CoV-2 infection warrants investigation.

Our findings are consistent with FFA β-oxidation being a hallmark of tissue-reparative monocyte-derived macrophages, coupling efferocytosis to the production of protective mediators: such as IL-10^63^, specialized pro-resolving lipid mediators derived from FFA substrates^64,65^ and growth factors such as amphiregulin^66^, that promote epithelial repair and maintain alveolar-capillary barrier integrity^67^. Additional energy-demanding mechanisms deployed by tissue-reparative macrophages include the disposal and recycling of damaged mitochondria^68–70^, via transmitophagy^37^ and possibly the transfer of functional mitochondria to damaged cells^71^.

We provide a mechanistic framework for understanding age-related COVID-19 susceptibility. Aging is associated with a progressive decline in adipose tissue lipolytic capacity^23–25^, which coupled with age-dependent changes in myeloid cell function^72^, creates a metabolic-immune vulnerability that predisposes older individuals to severe COVID-19. This suggests that therapeutic strategies to enhance FFA availability or restore monocyte metabolic function could be used as targeted interventions for elderly COVID-19 patients, with potential applicability to other age-sensitive infectious diseases.

## MATERIAL AND METHODS

### Mice

Mice (male and female) were bred and maintained under specific pathogen-free (SPF) conditions at the Gulbenkian Institute of Molecular Medicine (GIMM), housed at standard *vivarium* temperature (22°C) in a 12-hour light/dark cycle lights on from 8:00 am (ZT 0) to 8:00 pm (ZT 12). Mice were maintained with free access to water and standard chow pellets (*ad libitum*). All experimental protocols were approved in a two-step procedure, by the Animal Welfare Body of the GIMM and by the Portuguese National Entity that regulates the use of laboratory animals in research (Direção Geral de Alimentação e Veterinária; DGAV). Experimental procedures followed the Portuguese (Decreto-Lei n° 113/2013) and European (Directive 2010/63/EU) legislation. C57BL/6J mice were obtained from the GIMM animal facility. C57BL/6J *AdipoQ^Cre/Wt^Pnpla2^fl/fl^* (*i.e., Pnpla2^AdipoQ^*^Δ/Δ^) mice and littermate control *Pnpla2^fl/fl^* mice were previously described^73^. C57BL/6J *Ccr2^−/-^* were previously described^74^. C57BL/6J *Rag2^−/-^*were previously described^75^. C57BL/6J *Ccr2^−/-^ Rag2^−/-^* were generated by crossing C57BL/6J *Ccr2^−/-^* with C57BL/6J *Rag2^−/-^*. C57BL/6J *Cd36^−/-^* were previously described^76^. *Hmgcs2^fl/fl^* mice were previously described^77^. *Alb^Cre/WT^ Hmgcs2^fl/fl^*were generated by crossing *Alb^Cre/WT^* mice (#003574-JAX) with *Hmgcs2^fl/fl^* mice. *Cpt1a^fl/fl^* mice were obtained from Prof. Laura Herrero (University of Barcelona, Spain)^78^. C57BL/6J *LysM^Cre/WT^ Cpt1a^fl/fl^*mice were generated by crossing *LysM^Cre/WT^* with *Cpt1a^fl/fl^* mice by Prof. David Sancho (Nacional de Investigaciones Cardiovasculares Carlos III (CNIC), Madrid, Spain). *LysM^Cre/WT^ R26^tdT/WT^* mice, in which a *Rosa26*-tandem dimer (td) Tomato-Flox-stop-Flox allele driving the expression of td Tomato (tdT) by Cre-driven excision, under the control of *Rosa26* promoter (*tdT^R^*^26^) mice were crossed with *LysM^Cre/WT^* mice to enable tdTomato expression specifically in myeloid-lineage cells (*tdT^LysM^)*^37^.

### Cells and viruses

To generate SARS-CoV-2 viral stocks, VeroE6 cells were inoculated with mouse adapted (MA)-SARS-CoV-2 isolate hCoV-19/mouse/USA/IA-N501Y-MA30/2021 (Lineage B) (BEI Resources #NR-56222) or human specific SARS-CoV-2 USA/MD-HP01542/2021 (Lineage B.1.351) (BEI Resources #NR-55282) to generate the P1 stocks. Supernatant was clarified by centrifugation (700 x g × 5min) and then aliquoted for storage at − 80°C. Virus titer was determined by plaque assay using Vero E6 cells.

### Plaque assay

For viral titration, Vero E6 cells were seeded for 24 h, infected with different dilutions of SARS-CoV-2 virus in DMEM supplemented with 2 mM L-glutamine and 1% (v/v) penicillin/streptomycin and devoid of serum for 1h at 37°C and 5% CO2. After 1h, the cells were overlayed by 1% carboxymethyl cellulose supplemented with 2.5% serum and incubated for 5 days at 37°C and 5% CO_2_. The media was aspirated and the cells were fixed and stained with 0.1% toluidine blue in 4% formaldehyde O/N. The medium was aspirated and the cells were washed with PBS and viral titer was determined.

### Viral infection and disease assessment

Mice were maintained at the GIMM ABSL-3/ABSL-2 facility. Briefly, mice were anesthetized with isoflurane and intranasally inoculated with SARS-CoV-2 virus or influenza A virus. For SARS-CoV-2 virus, mice were intranasally inoculated with either 1.75×10^5^ PFU of SARS-CoV-2 (Lineage B.1.351, 50μL per nostril) or 0.2 × 10^4^ PFU of MA-SARS-CoV-2 (25μL per nostril). For influenza infections, mice were inoculated with 50, 75 or 150 PFU of influenza A virus strain A/Puerto Rico/8/34 (PR8, 30 µL). Mice were monitored daily from day 0 (day of infection) onward for clinical severity score (CSS, including activity, level of consciousness, response to stimulus and respiration), body weight (Ohaus® CS200 scaler, Sigma Aldrich), rectal temperature (Rodent thermometer; BIO-TK8851, Bioset) and blood ketone bodies (FreeStyle Precision Neo; Abbott) for 14 days.

### Lung viral titers

Viral titers were determined at day 3, 5 and 7 post-infections from the right upper lobes of MA-SARS-CoV-2 mice. Samples were homogenized in serum-free DMEM (Gibco) using tungsten carbide beads (Qiagen) in a Tissue Lyzer II (1600 MiniG) for 2 min. The supernatant was clarified by centrifugation at 2,000 x g for 5 min. Viral titer was determined by plaque assay using Vero E6 cells.

### Histopathological analysis

Mice were sacrificed by CO_2_ inhalation, transcardially perfused *in toto* with ice-cold PBS (1X, 20 mL). For histological analysis, lungs were fixed in 10% formalin for 24h, embedded in paraffin, sectioned (3 μm) and stained with hematoxylin & eosin (H&E). Whole sections from formalin-fixed and H&E-stained tissues were examined using a DM2000 LED microscope (Leica Microsystems), and digital images were acquired with a NanoZoomer-SQ digital slide scanner (Hamamatsu Photonics). Lung sections were evaluated using a semi-quantitative histopathological scoring system^79^. The assessment encompassed multiple parameters including alveolar edema, emphysema, atelectasis, and inflammatory infiltrates at alveolar, interstitial, peribronchial and perivascular locations. Each parameter was graded based on the percentage of affected area per lobe or severity of involvement, with predominant inflammatory cell types identified. Vascular lesions, including endothelitis, vasculitis, and thrombi formation, were scored for presence and severity. A composite total score was calculated by summing individual parameter scores to quantify overall lung pathology.

### Lung permeability assay

Lung permeability was measured using fluorescently labeled dextran (10 kDa), essentially as described^80^. Briefly, 25 μL of 50 mg/mL rhodamine-dextran (Sigma #R8881) was dissolved in sterile PBS and instilled intranasally into the airways. Mice were sacrificed 1h thereafter (CO₂ inhalation), blood was collected (EDTA tubes) and centrifuged at 3,000 x g, 10 min. Plasma was harvested, and virus was inactivated by the addition of NP-40 (10 μL in 200 μL plasma). Fluorescence intensity was measured at an excitation wavelength of 570 nm and an emission wavelength of 594 nm using a GloMax reader.

### Protein determination in BAL

To measure protein concentration in the BAL, as a proxy of increase in lung capillary leakage, mice were sacrificed (CO₂ inhalation), and the thorax was opened to expose the trachea. The lungs were washed by intratracheal instillation of sterile PBS (20G catheter, 3 × 0.5 mL) followed by gentle aspiration. The recovered fluid was centrifuged at 300 x g, 10 min and the cell-free supernatant was collected and incubated at 70°C for 10 min to inactivate the virus. Protein concentration was determined using Bradford reagent by measuring the absorbance at 595 nm.

### Adipose tissue mass

Mice were euthanized, perfused *in toto* with ice-cold PBS. The eWAT and scWAT fat pads were harvested and weighed (Entris^®^ II, Satorius).

### Adipose tissue immunofluorescence

Mice were sacrificed by CO_2_ inhalation, transcardially perfused *in toto* with ice-cold PBS (1X, 20 mL). eWAT was harvested and fixed with 4% PFA with gentle shaking, overnight (O/N) at 4°C. Next day, tissues were washed thrice with 1X PBS (30 min each) and permeabilized (B1N buffer: 0.1% Triton X-100, 0.3 M Glycine, 0.01% Sodium Azide in distilled water) O/N with gentle shaking at RT. Samples were blocked using PtxwH buffer (1x PBS, 0.1% Triton X-100, 0.05% Tween-20, 2mg/mL Heparin, 0.01% Sodium Azide, 3% donkey serum) O/N at RT. Samples were incubated with primary antibodies: anti-ATGL (PNPLA2/ATGL Antibody, #AF5365 Novus biologicals) and anti-pHSL (Phospho-HSL Ser660, Cell Signaling Technology, #45804,) prepared in blocking buffer for 3 days with gentle shaking at RT. The samples were then washed with PtxwH buffer, 5 times (1h each). After washing, the samples were incubated with secondary antibody, donkey anti-sheep IgG H&L (Alexa Fluor® 647) (Abcam, #ab150179) and donkey anti-rabbit IgG H&L (Alexa Fluor® 568) (Abcam, #ab175470) prepared in blocking buffer O/N with gentle shaking at RT. The samples were then washed with PtxwH buffer, 5 times (30 min each). After washing, the samples were incubated with BODIPY 493/503 (Cayman, #25892) and DAPI (in PBS) for 2h at 37°C. The samples were then washed with PBS, 3 times (30 min each) and imaged using a laser scanning confocal microscope (LSM980, Zeiss) and fluorescence intensity was measured using ImageJ 1.53k software (Rasband, W.S., ImageJ, U.S. NIH, Bethesda, Maryland, USA, https://imagej.nih.gov/ij/,1997-2014).

### Western Blot

Mice were sacrificed and perfused as described before, eWAT was collected and snap frozen in liquid nitrogen. The tissue was homogenized in NP40 extraction buffer (100 mL of extraction buffer containing 3 mL of 5M NaCl, 10 mL of 10% NP-40, 5 mL of 1M Tris, 82 mL of H2O, protease inhibitor: complete Mini, EDTA-free, Sigma #11836170001) using a tissue lyzer (Qiagen) with tungsten carbide beads (Qiagen). Samples were incubated 1h on ice and then centrifuged (20,000 x g, 15 min, 4°C). The middle layer (without the fat) was transferred to another tube and centrifuged again, as before. The supernatant was transferred and protein was quantified via Bradford assay (BioRad, #5000006). Protein (30 µg) was resolved on a 10% SDS-PAGE and transferred to Polyvinylidene fluoride (PVDF) membranes. The membranes were blocked (5% bovine serum albumin: BSA in 1XT-TBS) and incubated with primary antibodies overnight (4°C): Anti-mouse phospho-HSL (Ser660) (#45804, Rabbit, 1:1000), Anti-mouse HSL (#4107, Rabbit, 1:1000), Anti-mouse ATGL (#2439, Rabbit, 1:1000) and Anti-mouse β-Actin (#4967, Rabbit, 1:1000) all from Cell Signaling Technology. Membranes were washed (3x, 1X T-TBS) and incubated (1h, RT) with corresponding peroxidase-conjugated secondary antibodies (goat anti-rabbit IgG (H+L) HRP #31460). Peroxidase activity was detected using SuperSignal West Pico PLUS Chemiluminescent Substrate (ThermoFisher Scientific). Blots were developed using Amersham Imager 680 (GE Healthcare), equipped with a Peltier cooled Fujifilm Super CCD. Densitometry analysis was performed with ImageJ (Rasband, W.S., ImageJ, U.S. NIH, Bethesda, Maryland, USA, https://imagej.nih.gov/ij/, 1997-2014), using only images without saturated pixels.

### FFA measurement from the explants

Mice were sacrificed and perfused as described before. The eWAT was harvested and weighed. eWAT (15mg) was taken from each mouse and incubated in a 24 well plate containing 1 mL of pre-warmed Krebs-Ringer Solution, HEPES-buffered (Thermofisher, #J67795.AP) with 2% FA free BSA (AlbumiNZ™ Bovine Albumin Low Free Fatty Acid #02199899). The plate was then incubated at 37 °C for 2h. After 2h, incubation media was collected in 1.5 mL tubes. The tubes were incubated at 65°C for 10 min. For FFA measurement, Free FA Quantitation Kit (Sigma, #MAK044) was used. Briefly 2, 4, 6, 8, and 10 nmol/well standards were prepared in a 96 well plate. 15 μL of sample were added to the well and volume was made up to 50 μL. 2 μL of acyl-CoA synthetase reagent was added to each sample and standard well, and incubated at 37 °C for 30 min. The master reaction mix was prepared according to the manufacture instruction and 50 μl was added to each well and the plate was incubated at 37 °C for 30 min. The absorbance was measured at 570 nm and the FFA concentration of unknown samples were determined.

### Glycerol measurement from the explants

Mice were sacrificed and perfused as described before. The epidydimal (eWAT) white adipose tissue fat pads were harvested and weighed. eWAT (15mg) was taken from each mouse and incubated in a 24 well plate containing 1 mL of pre-warmed Krebs-Ringer Solution, HEPES-buffered (Thermofisher, #J67795.AP) with 2% FA free BSA (AlbumiNZ™ Bovine Albumin Low Free Fatty Acid #02199899). The plate was then incubated at 37 °C for 2h. After 2h, incubation media was collected in 1.5 mL tubes. The tubes were incubated at 65°C for 10 min. For glycerol measurement, glycerol standard solution (Sigma #G7793, Conc: 0,26 mg/mL of glycerol) was used. Briefly standards were prepared by diluting Glycerol Standard Solution 2-fold in PBS. 50 μl of standard or sample was added to the 96 well plate. 100 μL of Free Glycerol Reagent (Sigma, #F6428) was added to each well and incubate for 15 min at RT. Absorbance was measured at 540nm and the glycerol levels of unknown samples were determined.

### Blood analysis

Mice were sacrificed by CO_2_ inhalation, and the blood was collected by cardiac puncture in EDTA tubes. The blood was centrifuged (1,500 x g, 10 min, 4 °C) and plasma was collected. To inactivate the virus, plasma was treated with RNase A (sigma #10109169001, 1:1000) and Triton X 100 (sigma #515958069, 1%). Serological analyses were outsourced by DNATech (Portugal; http://www.dnatech.pt/web/). Blood levels of FFA, glycerol, ALT, AST and urea were assessed.

### Untargeted Lipidomics

Young (3-4 months) and old (16 months) *Pnpla2^fl/fl^* and *Pnpla2^AdipoQΔ/Δ^* mice were infected with SARS-CoV-2 (Lineage B.1.351). Mice were sacrificed at 3 days post infection and blood was collected through intracardial puncture (23 G needle pre-coated with 0.5M EDTA), plasma was extracted by centrifugation (1500 x g; 10 min.) and 50 µL was used for lipid extraction. Isopropanol (450 µL containing internal standards) was added and after thorough vortexing and incubation (-20 °C; 20 min.) samples were centrifuged (10 min.; 14,000 x g, 4 °C). Supernatants were collected and used for LC-MS/MS analysis, performed on a Vanquish UHPLC system coupled to an Orbitrap Exploris 240 high-resolution mass spectrometer (Thermo Scientific, MA, USA) in negative and positive ESI (electrospray ionization) mode. Chromatographic separation was carried out on an ACQUITY Premier CSH C18 column (Waters; 2.1 mm × 100 mm, 1.7 µm) at a flow rate of 0.3 mL/min. The mobile phase consisted of water:ACN (40:60, v/v; mobile phase A) and IPA:ACN (9:1, v/v; mobile phase B), which were modified with a total buffer concentration of 10 mM ammonium acetate + 0.1 % acetic acid (negative mode) and 10mM ammonium formate + 0.1% formic acid (positive mode), respectively. The following gradient (23 min. total run time including re-equilibration) was applied (min/%B): 0/15, 2.5/30, 3.2/48, 15/82, 17.5/99, 19.5/99, 20/15, 23/15. Column temperature was maintained at 65°C, the autosampler was set to 4°C and sample injection volume was set to 3 µL (positive mode) and 5 µL (negative mode). Analytes were recorded via a full scan with a mass resolving power of 120,000 over a mass range from 200 – 1700 m/z (scan time: 100 ms, RF lens: 70%). To obtain MS/MS fragment spectra, data-dependent acquisition was carried out (resolving power: 15,000; scan time: 54 ms; stepped collision energies [%]: 25/35/50; cycle time: 600 ms). Ion source parameters were set to the following values: spray voltage: 3250 V / 3000 V, sheath gas: 45 psi, auxiliary gas: 15 psi, sweep gas: 0 psi, ion transfer tube temperature: 300°C, vaporizer temperature: 275°C. Level 1 feature identification was based on the MS-DIAL^81^ LipidBlast V68 library.

All experimental samples were measured in a randomized manner. Pooled quality control (QC) samples were prepared by mixing equal aliquots from each processed sample. Multiple QCs were injected at the beginning of the analysis in order to equilibrate the analytical system. A QC sample was analyzed after every 5th experimental sample to monitor instrument performance throughout the sequence. For determination of background signals and subsequent background filtering, an additional processed blank sample was recorded.

### Lipidomics Analysis

Lipid abundances were normalized using the median Total Ion Current (mTIC)^82^ method and subsequently log₂-transformed prior to statistical analysis. This transformation serves to stabilize variance and render the data approximately normally distributed, which was verified per lipid feature using the Shapiro-Wilk test (scipy.stats, Python). Detections from positive and negative ionization modes were merged into a unified dataset; in cases where the same lipid species was identified in both modes, only the detection with the highest intensity was retained to avoid redundancy. Differential abundance between groups was assessed by fitting an independent linear model fitted by ordinary least squares, for each lipid feature, specified as: lipid_abundance ∼ group, where group defined the mice age and genetic background. All models were implemented using the statsmodels package in Python. Because lipid abundances were modelled on the log₂ scale, the estimated group coefficient directly represents the log₂ fold change (log₂FC) between conditions. Pairwise group contrasts were extracted from each fitted model using the t_test_pairwise function, which evaluates linear combinations of model coefficients and yields nominal p-value per contrast per lipid. To account for the inflated risk of false positives arising from the simultaneous testing of hundreds of lipid species, all nominal p-values were corrected using the Benjamini–Hochberg procedure (statsmodels.stats.multitest.multipletests), controlling the false discovery rate (FDR) at the dataset level. A lipid species was deemed differentially regulated if it met both criteria: an FDR-adjusted p-value < 0.05 and an absolute log₂ fold change greater than 1, corresponding to at least a two-fold difference in abundance between groups. Volcano plots were generated to visualize the joint distribution of effect size and statistical significance (p-value) using the plotly Python library.

The same lipid classes were analyzed in the human datasets^30–32^, where fold changes were calculated from the available intensity data. To maximize consistency across re-analyses and between studies, a t-test was applied to identify individually significant lipid species, using a fold-change threshold of absolute log₂ fold change greater ≥ 0.58 (the smallest threshold across all studies) and a corrected p-value (FDR) < 0.05.

### Flow cytometry

#### Lungs Myeloid and lymphoid panel

Mice were sacrificed and perfused as described before. Lungs were harvested and digested with cRPMI media containing 1mg/mL collagenase IV (sigma #C5138) and 100μg/mL DNase I (sigma #10104159001). Cell suspensions were filtered through a cell strainer (70 µm, Corning) and RBCs were lyzed using ACK lysis buffer. Cells were then stained with Fc block (BD pharmingen, #553142) and fixable live dead yellow dye (LIVE/DEAD™ Fixable Yellow Dead Cell Stain Kit, Invitrogen, #L34959) for 20 min (4°C). The cells were washed with PBS containing 2% FBS and centrifuged (700 x g, 2 min, 4°C). The cells were fixed using 4% paraformaldehyde for 30 min (4°C). Cells were washed with PBS containing 2% FBS and centrifuged (700 x g, 2 min, 4°C). Cells were stained with following antibodies: For lungs, CD45 APCe780 (#47-0451-82), Ly6G AF700 (#56-9668-82) from Invitrogen, CD11b BV785 (#101243), NK1.1 PE/Dazzle 594 (#108748), TCRγδBV605 (#118129), CD3 AF647 (#152320), MHCII BV711 (#107643), F4/80 BV421(#123132), Ly6C PE/Cy7 (#128018), B220 PercP/Cy5.5 (#103236), CD4 PE/Cy7 (#100422), CD19 BV711 (#115555), CD69 PE (#104508), CD44 BV421 (#103039), CD25 AF488 (#102017), all from Biolegend, CD11C PE (BD Pharmingen, #557401), CD8 PerCP/Cy5.5 (Life technology, #45-0081-82) for 30 min (4°C). The cells were washed with PBS containing 2% FBS and centrifuged (700 x g, 2 min, 4°C) and analyzed in a LSRFortessa X-20 (BD Biosciences). FACS data was analyzed with FlowJo V10.8.1.

#### Bone marrow hematopoiesis and myelopoiesis

Mice were sacrificed and perfused as described before. The femur was harvested and cut at both ends and placed in a 0.5 mL tubes with bottom pinched with a hole by 20G needle. The 0.5 mL tube was then placed in a 1.5 mL tube, and the cells were harvested by centrifugation at (5,000 x g, 10 sec, 4 °C). RBCs were lyzed using ACK buffer and cells were then stained with fixable live dead yellow dye for 20 min (4°C). For measuring FA uptake and intracellular lipid content, the cells were stained with BODIPY™ FL C_12_ (Invitrogen #D3822) and BODIPY 493/503 (Cayman, #25892), respectively for 30 min at 37°C. Cells were washed with PBS containing 2% FBS and centrifuged (700 x g, 2 min,4°C). Cells were then stained with CD16/32 BV421 (Biolegend, #101331) for 15 min (4°C). Cells were washed with PBS containing 2% FBS and centrifuged (700 x g, 2 min, 4°C). The cells were stained with the following primary antibodies: Sca-1 FITC (#122505), Ly6C PerCP/Cy5.5 (#128012), CD115 PE (#135505), CD34 PE/Cy5 (#119311), F4/80 AF647 (#123122), CD11C BV605 (#117333), CD3 Biotin (#100243), CD49b Biotin (#103521), CD90.1 Biotin (#202510), all from Biolegend, CD135 PerCPeF710 (#46-1351-82), CD150 eF450 (#48-1502-82), MHCII eF506 (#69-5321-82) from Invitrogen, CD48 PE-Vio 770 (Miltenyi Biotec S. L, #130-102-363), c-KIT APC (BD Pharmingen, #553356), CD19 Biotin (eBioscience, #553784), for 25 min (4°C). The cells were washed with PBS containing 2% FBS and centrifuged (700 x g, 2 min, 4°C). The cells were then stained with secondary antibody PE/Fire700 Streptavidin (Biolegend, #405174) for 15 min (4°C) and washed with PBS containing 2% FBS and centrifuged (700 x g, 2 min, 4°C). The cells were fixed using 4% PFA for 30 min (4°C). The cells were washed with PBS containing 2% FBS and centrifuged (700 x g, 2 min, 4°C) and analyzed in a Cytek Aurora (Cytek Biosciences). FACS data was analyzed with FlowJo V10.8.1.

#### Puromycin incorporation

Mice were sacrificed and femur was harvested as described before. After RBC lysis, 2 × 10^5^ cells were seeded in a 96 well plate and treated with either vehicle (PBS) or etomoxir (4μM, Cayman, #11969-50) for 20 min. Puromycin (final concentration 10 µg/mL, sigma # P7255) was added to cultures for 25 min. Cells were washed with PBS containing 2% FBS and centrifuged (700 x g, 2 min, 4°C). Cells were then stained with fixable live dead yellow dye, CD16/32 BV421 and other antibodies as described before. The cells were fixed and permeabilized using FIX & PERM kit (ThermoFisher #GAS004). After permeabilization, cells were stained with anti-puromycin antibody (Merck # MABE343). The cells were washed with PBS containing 2% FBS and centrifuged (700 x g, 2 min, 4°C) and analyzed in a Cytek Aurora (Cytek Biosciences). FACS data was analyzed with FlowJo V10.8.1. Mean fluorescence intensity (MFI) was used to quantify puromycin incorporation^53^.

### Single-cell RNA-seq sample preparation

Lungs were harvested from *Pnpla2^fl/fl^* and *Pnpla2^AdipoQ^*^Δ/Δ^ infected mice and digested with cRPMI media containing 1mg/mL collagenase IV and 100μg/mL DNase I. Cell suspensions were filtered through a cell strainer (70 µm), and RBCs were lyzed using ACK lysis buffer. Cells were then stained with Fc block (BD pharmingen, #553142) and fixable live dead yellow dye (LIVE/DEAD™ Fixable Yellow Dead Cell Stain Kit, Invitrogen, #L34959) for 20 min (4°C). The cells were washed with PBS containing 2% FBS and RNASE inhibitor (sigma # 3335399001) and centrifuged (700 x g, 2 min, 4°C). Single-cell suspensions were stained with CD45.2 (clone 104, Biolegend # 109808) and TotalSeq-C-tagged anti-mouse Hashtag 5 to 10 (Biolegend #155875, #155877, #155879, #155881, #155883, #155885). The cells were washed twice with PBS containing 2% FBS and RNASE inhibitor. The cells were sorted into CD45^+^ and CD45^−^ populations using Cytek Auroa cell sorter (Cytek Biosciences) and adjusted to 1,333 cells/μL in PBS. Single-cell library preparation was performed according to the Chromium GEM-X Single Cell 5’ Kit v3 (10x Genomics). Library preparations were done based on manufacturer’s instruction. PCR were done using a C1000 Touch Thermal Cycler (Bio-Rad). cDNA and library quality control and quantification were performed with a HS NGS Fragment kit (Agilent Technologies). Gene expression libraries were indexed using the Dual Index Plate TT, Set A and antibody barcode libraries were indexed with Dual Index Kit TN, Set A. All libraries were verified by Fragment Analyzer prior to sequencing on a NextSeq 2000 instrument (Illumina) using the P2 reagent kit with 100-cycle runs.

### sc-RNA analysis

#### RNA expression quantification, demultiplexing and quality control

Gene expression was quantified using Cell Ranger v9.0.1^83^ with default parameters and aligned against the mouse reference transcriptome GRCm39-2024-A. To assign each cell to its respective mouse of origin and identify potential doublets, demultiplexing was performed using the HTODemux function within the Seurat R package. Following quantification, cells were filtered based on quality control metrics to remove low-quality cells. Thresholds were determined by manual inspection of the distributions of the quality control metrics. For CD45^+^ dataset, cells with fewer than 5,000 detected genes (nFeature_RNA), a mitochondrial read fraction (percent.mt) below 25%, and a total unique UMI count (nCount_RNA) below 25,000 were retained. For CD45^−^ cells, following criteria were applied: fewer than 5,000 detected genes, a mitochondrial content under 8%, a total UMI count below 20,000, and a ribosomal gene percentage (percent.rp) below 17%. Only cells meeting these predefined thresholds were retained for downstream analysis.

#### Downstream analysis

Downstream analyses were conducted using the Seurat R package (v5.4.0)^84^. To enable lineage-specific analysis, the CD45^+^ population was partitioned into two distinct datasets, myeloid and lymphoid, while the CD45- population was processed as a single dataset. For all objects, normalization and scaling were performed via the SCTransform workflow^85^, which identified the top 5,000 highly variable features while simultaneously regressing out variation associated with total UMI counts (nCount_RNA) and mitochondrial transcript percentages (percent.mt). Dimensionality reduction was performed through Principal Component Analysis (PCA), with the appropriate number of principal components (PCs) selected based on the inflection point of ElbowPlot visualizations. Specifically, 28, 30, and 40 PCs were chosen for the myeloid, lymphoid, and CD45^−^ datasets, respectively. These components were used as input for UMAP dimensionality reduction using the RunUMAP function. Clustering was executed using the FindClusters function with dataset-specific resolutions: 0.6 for myeloid cells, 0.2 for lymphoid cells, and 0.4 for the CD45^−^ fraction. Differential gene expression analysis was performed using the FindMarkers function, with significant markers defined by a log_2_ fold change (log_2_FC) threshold greater than 0.5 and a p-value below 0.05, as determined by the non-parametric Wilcoxon Rank Sum test. Cell cluster annotation was performed by cross-referencing differentially expressed genes with established markers from the literature^86^.

#### Cell-cell communication

To characterize intercellular signaling dynamics, the LIANA framework (v0.1.14)^42^ was applied independently to the *Pnpla2^AdipoQΔ/Δ^* and *Pnpla2^fl/fl^*mouse datasets. For each condition, the dataset containing myeloid, lymphoid, and CD45^−^ populations served as input. To improve robustness of interaction inference, infrequent cell populations were excluded using the filter_nonabundant_celltypes function with default parameters. Cell-cell communication was then inferred via the liana_wrap function, specifically employing the CellPhoneDB method^87^ against the mouse OmniPath database^88^ as the reference resource. The resulting interaction tables were filtered to retain only significant entries (*p* <0.01). To further prioritize high-confidence signaling events, interactions within the top 80^th^ percentile (percentile > 20) based on the mean ligand-receptor expression (lr.mean) were considered for downstream analysis. To quantify shifts in connectivity between the two experimental groups, the absolute difference in interaction counts for each cell-type pair was computed by subtracting the interaction matrices between conditions. For interactions conserved across both conditions, the log_2_FC of lr.mean values was calculated to assess the relative change in signaling magnitude between the *Pnpla2^AdipoQΔ/Δ^* and *Pnpla2^fl/fl^*.

### Adoptive transfer of bone marrow derived monocytes (BMDMo)

Adoptive transfer of bone marrow-derived monocytes (BMDMo) was performed essentially as described^89^. Briefly, BMDMo cells were freshly harvested from C57BL/6J mice and placed in culture for 5 days in Ultra-low-adherence T75 cell culture flasks (Corning #734-4139) with BMDMo differentiation culture medium (RPMI, 10% FCS, 1% Penicillin/Streptomycin, supplemented with 10% L929 culture supernatant. On day 5, BMDMo were collected, washed in RPMI (10 mL; 300 x g, 5 min, 4 °C) and resuspended in RPMI. On days 3, 5 and 7 post-infection, 50 µL (25μl per nostril) of BMDMo suspension was injected intranasally in each mouse (3 × 10^6^ cells/mouse/).

### Drug treatments

Etomoxir (Cayman, 11969-50) was freshly prepared before administration by diluting into 1X PBS. Mice were administered *i.p.* with 25 mg/kg of etomoxir or vehicle before infection and on days 1, 3 and 5 post infection. Sulfosuccinimidyl oleate sodium (MedChem express, #HY-112847A) was freshly prepared before administration by diluting into 10% DMSO, 30% PEG400 and 60% PBS. Mice were administered *i.p.* with 40 mg/kg of Sulfosuccinimidyl oleate sodium or vehicle before infection and on days 1, 3 and 5 post infection.

### Bone marrow Chimera

All bone marrow chimera combinations were generated by lethally irradiating (8.5 Gy) recipient mice and reconstituted 4 h post-irradiation with freshly isolated or cryo- preserved bone marrow cells isolated from donor mice (retroorbital injection of 2∼3×10^6^ cells in 100 μL RPMI). Successful hematopoietic cell reconstitution was confirmed by concomitant lethal irradiation of control mice that were not reconstituted with bone marrow cells.

### Statistical Analysis

Statistically significant differences between two experimental groups were assessed using two-tailed Unpaired t-test. Comparisons between more than two groups were carried out by one or two-way ANOVA with Tukey’s or Sídák multiple-comparisons test. Survival was represented by Kaplan–Meier plots and difference between the groups was assessed using the log-rank test. All statistical analyses were performed using GraphPad Prism 10 software. Differences were considered significant at a p value <0.05. ns: Not significant, p > 0.05; *p < 0.05, **p < 0.01; ***p < 0.001; ****p < 0.0001. In MA-SARS-CoV-2 experimental studies, mice that failed to exhibit body weight reduction within the first 5 days post-infection were identified as technical outliers and excluded from further analysis.

## Supporting information

Supplementary data for Figure 4F-I

## SUPPLEMENTARY FIGURES & LEGENDS

**Supplementary Figure 1.**
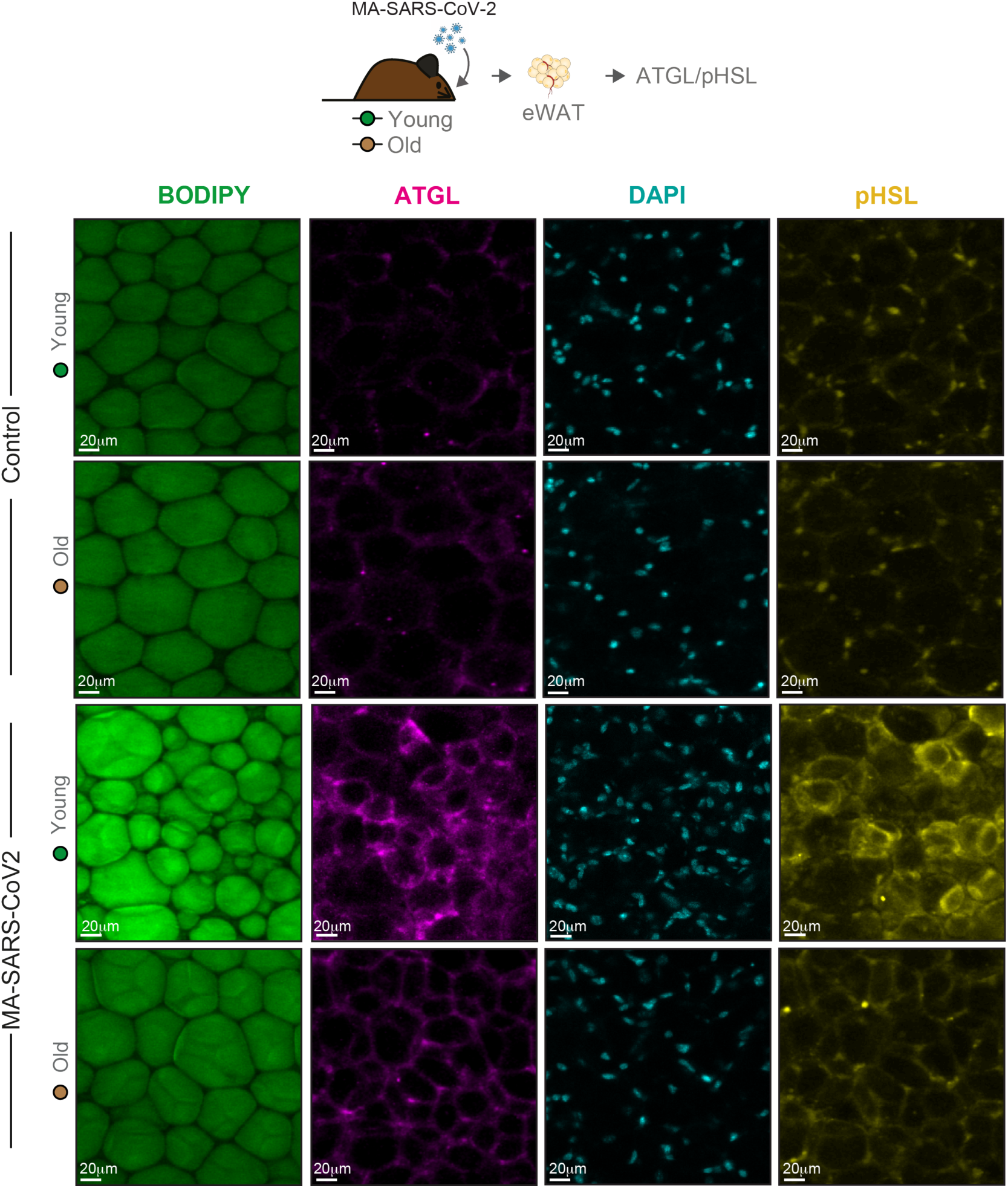
Age-dependent lipolytic responses to MA-SARS-CoV-2 infection. Deconvolution of representative immunofluorescence images of epididymal white adipose tissue (eWAT) from young (10-16 weeks) and old (>20 months) C57BL/6J mice under steady state (control) conditions or 5 days after MA-SARS-CoV-2 infection from figure 1G. eWAT stained for BODIPY 493/503 (lipid droplets, green), ATGL (magenta), pHSL (yellow) and DAPI (nuclei, cyan). Scale bars: 20 μm. Images are representative of n=6-7 mice *per* group (males) from 2 independent experiments.

**Supplementary Figure 2.**
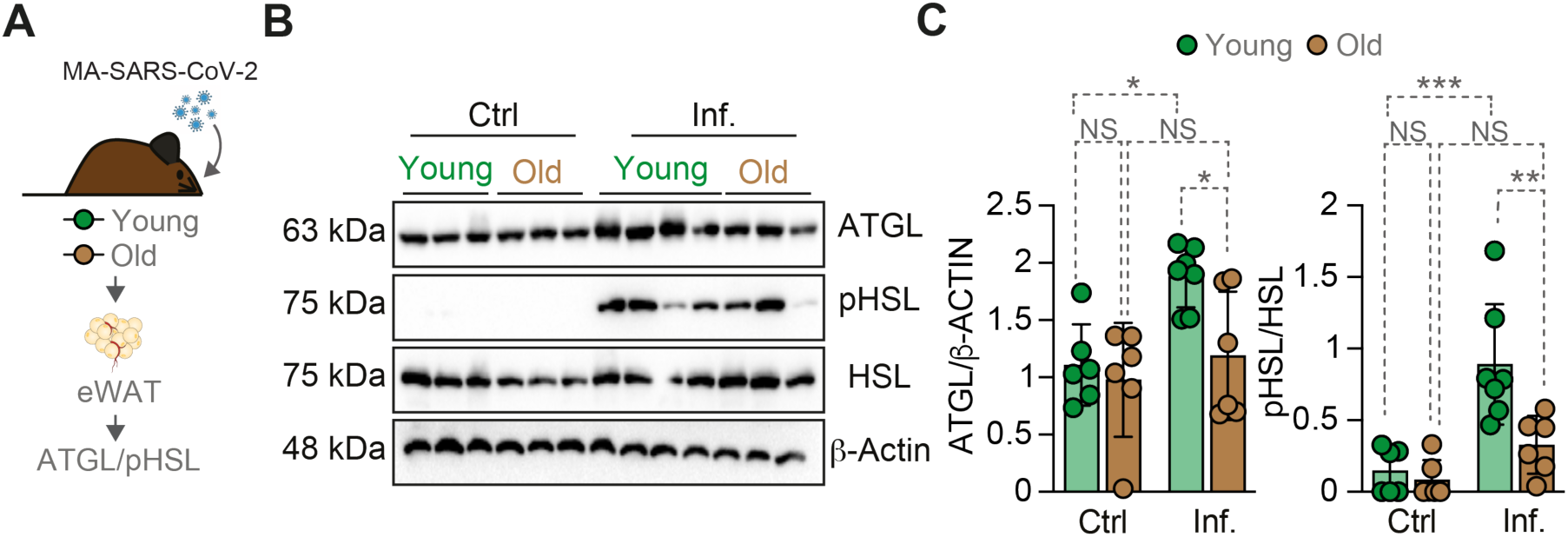
Age-dependent lipolytic responses to MA-SARS-CoV-2 infection. (**A**) Young (10-16 weeks) and Old (>20 months) C57BL/6J mice were infected with MA-SARS-CoV-2 and epididymal white adipose tissue (eWAT) was harvested at 5 dpi for western blot. (**B**) Detection of ATGL, pHSL, HSL and β-ACTIN protein expression from eWAT of C57BL/6J mice under steady state (control; ctrl) conditions or 5 days after MA-SARS-CoV-2 infection (n=6-7 mice *per* group, males & females). (**C**) Relative quantification of ATGL and pHSL/HSL protein expression normalized to β-Actin detected by western blot. Data shown as mean ± SD (C). *p* values determined by two-way ANOVA (C). NS, not significant; *p < 0.05, **p < 0.01; ***p < 0.001. Data pooled from two independent experiments with a similar trend.

**Supplementary Figure 3.**
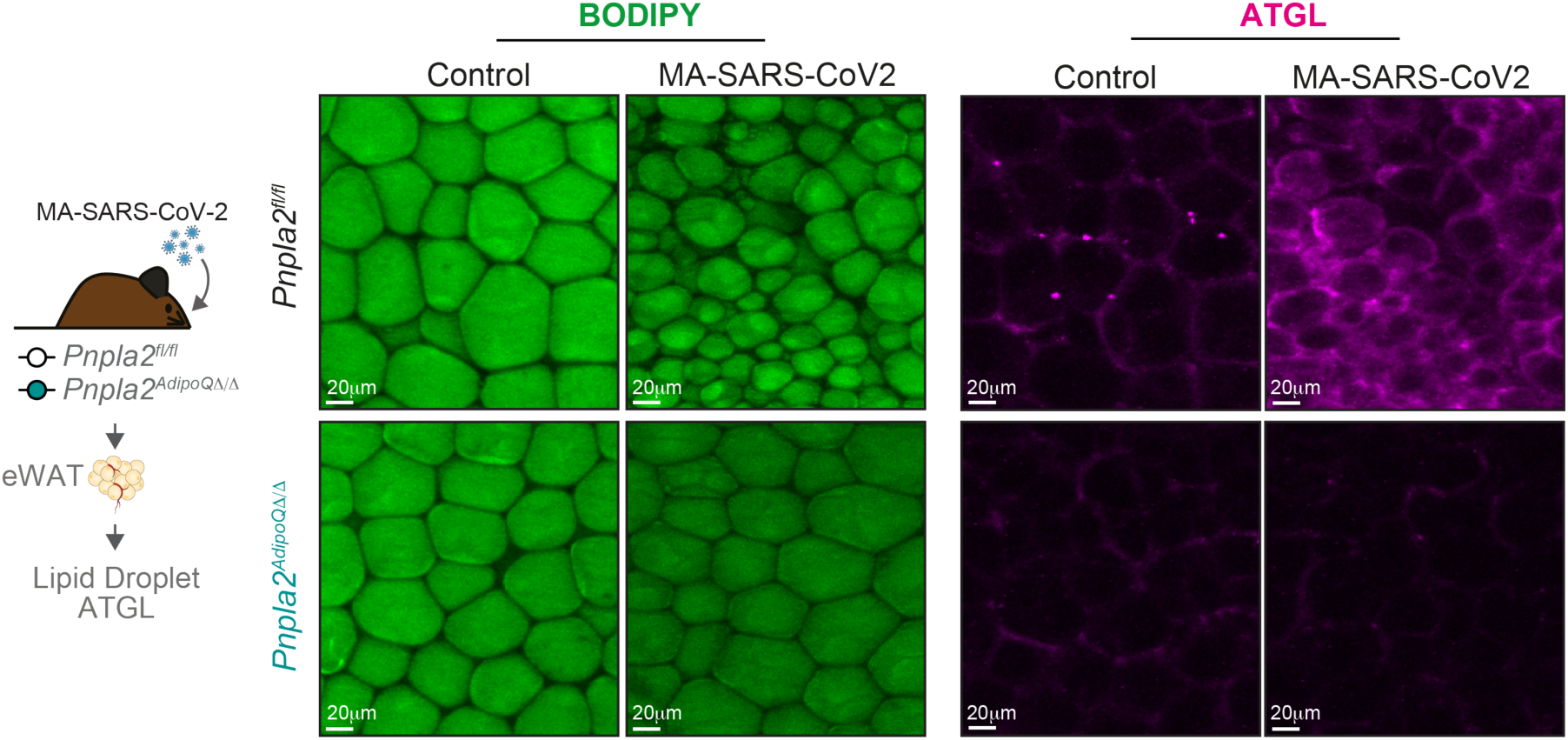
Adipocyte-specific ATGL deletion. Representative immunofluorescence images of epididymal white adipose tissue (eWAT) from Young *Pnpla2^fl/fl^* and *Pnpla2^AdipoQΔ/Δ^* mice under steady state conditions or following MA-SARS-CoV-2 infection at 5 dpi. Left panels show BODIPY 493/503 staining (green) marking lipid droplets; right panels show ATGL immunostaining (magenta). Scale bars: 20 μm. Images are representative of n=3-7 mice *per* group (males & females) from 2 independent experiments.

**Supplementary Figure 4.**
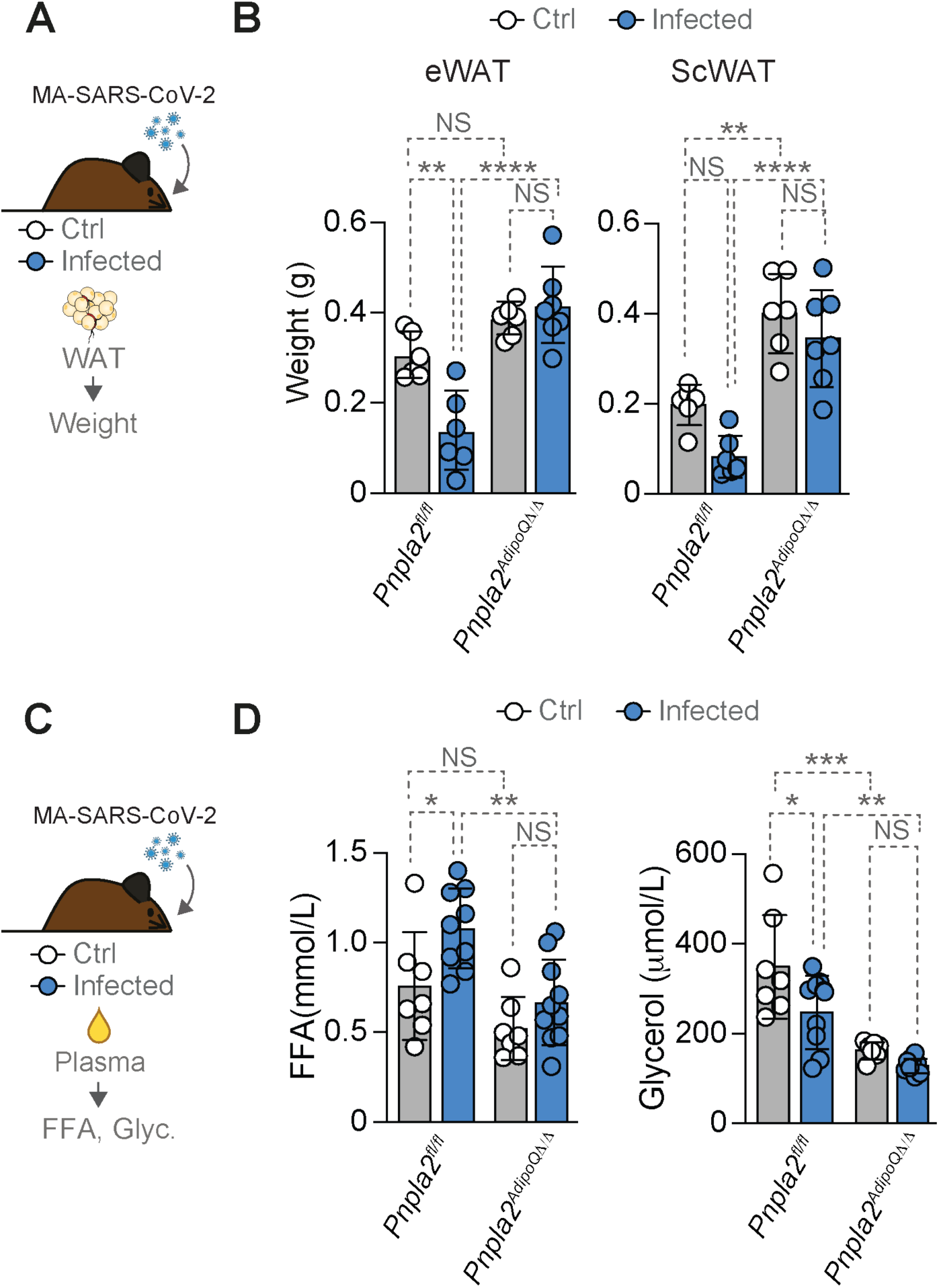
Adipocyte ATGL deletion prevents FFA and glycerol mobilization in response to MA-SARS-CoV-2 infection. (**A**) Young *Pnpla2^fl/fl^*and *Pnpla2^AdipoQΔ/Δ^* mice were infected with MA-SARS-CoV-2 and white adipose tissue (WAT) weights were measured at 5 dpi. (**B**) Epididymal WAT (eWAT) and subcutaneous WAT (ScWAT) weights (n=6-7 mice *per* group, males & females). (**C**) Plasma was collected from young *Pnpla2^fl/fl^* and *Pnpla2^AdipoQΔ/Δ^* mice at steady state or 5 dpi to measure (**D**) free fatty acids (FFA) and glycerol (n=7-10 mice *per* group, males & females). Data represented as mean *±* SD (B, D). Circles represent individual mice (B, D). *p* values were determined using two-way ANOVA (B, D). NS, not significant; *p < 0.05; **p < 0.01; ***p < 0.001, ****p < 0.0001. Data pooled from two (B) or three (D) independent experiments with a similar trend.

**Supplementary Figure 5.**
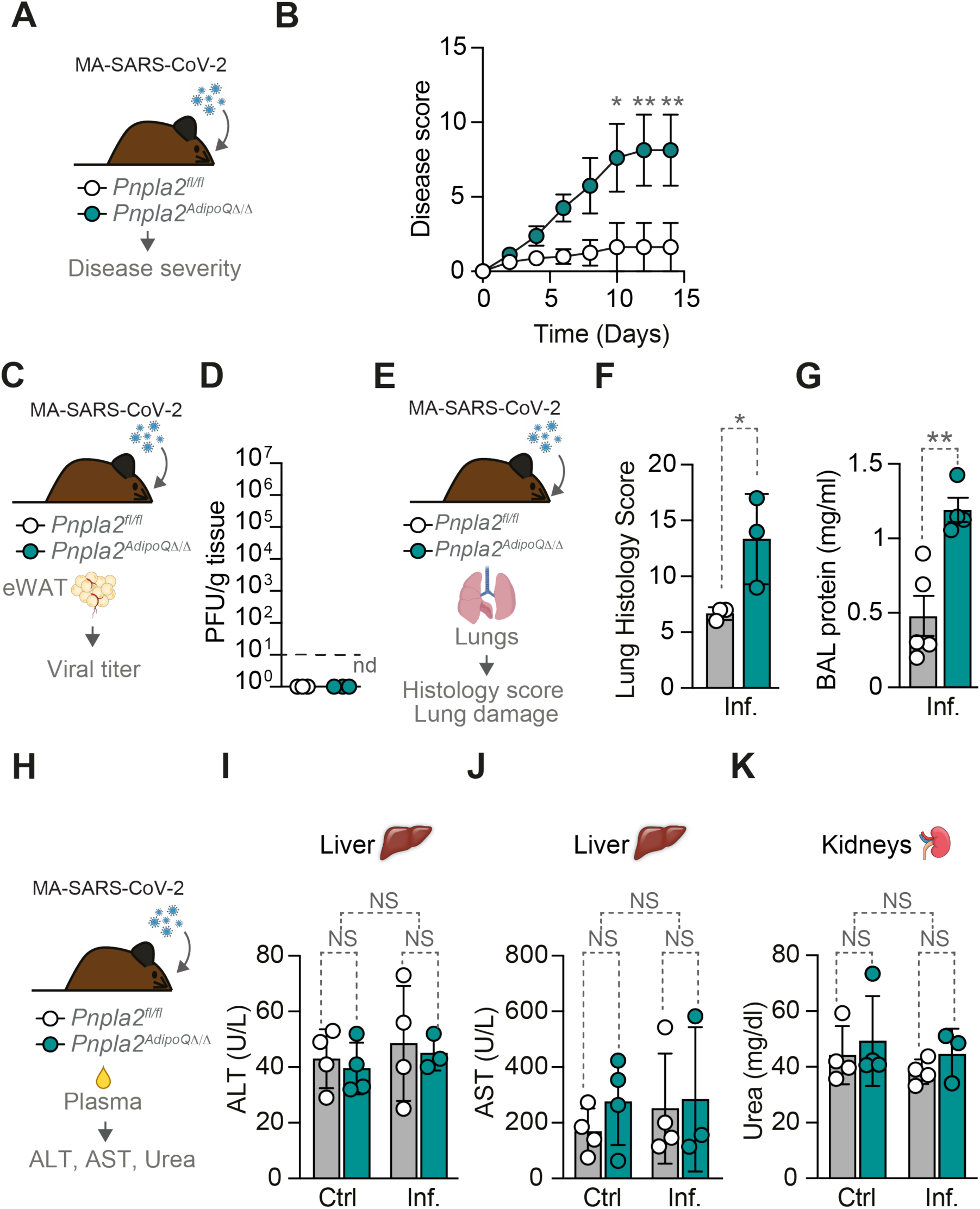
ATGL deletion causes progressive lung-specific pathology during MA-SARS-CoV-2 infection without systemic organ dysfunction. (**A**) Young *Pnpla2^fl/fl^* and *Pnpla2^AdipoQΔ/Δ^* mice were infected with MA-SARS-CoV-2 and monitored for (**B**) disease severity (n=8 mice per group, males & females). (**C**) eWAT was collected from young *Pnpla2^fl/fl^* and *Pnpla2^AdipoQΔ/Δ^* mice at 5 dpi to assess (**D**) viral titers (n=3 mice *per* group, males). Dashed line indicates limit of detection, nd: not determined. (**E**) Lung damage at 5 dpi, shown as (**F**) histology scores (n=3 mice per group, males) and (**G**) bronchoalveolar lavage (BAL) protein concentration (n=4-5 mice per group, males & females). (**H**) Plasma was collected from young *Pnpla2^fl/fl^*and *Pnpla2^AdipoQΔ/Δ^* mice at steady state or 5 dpi to assess (**I-K**) liver (ALT, AST) and kidney (urea) damage markers (n=3-4 mice *per* group, males & females). Data represented as mean *±* SD (B, F, I, J, K) or SEM (D, G). Circles represent individual mice (D, F, G, I, J, K). *p* values determined using two-way ANOVA (B, I, J, K) and unpaired t test (D, F, G). NS, not significant; *p < 0.05; **p < 0.01. Data pooled from two independent experiments with a similar trend (B, G) or one experiment (D, F, I, J, K).

**Supplementary Figure 6.**
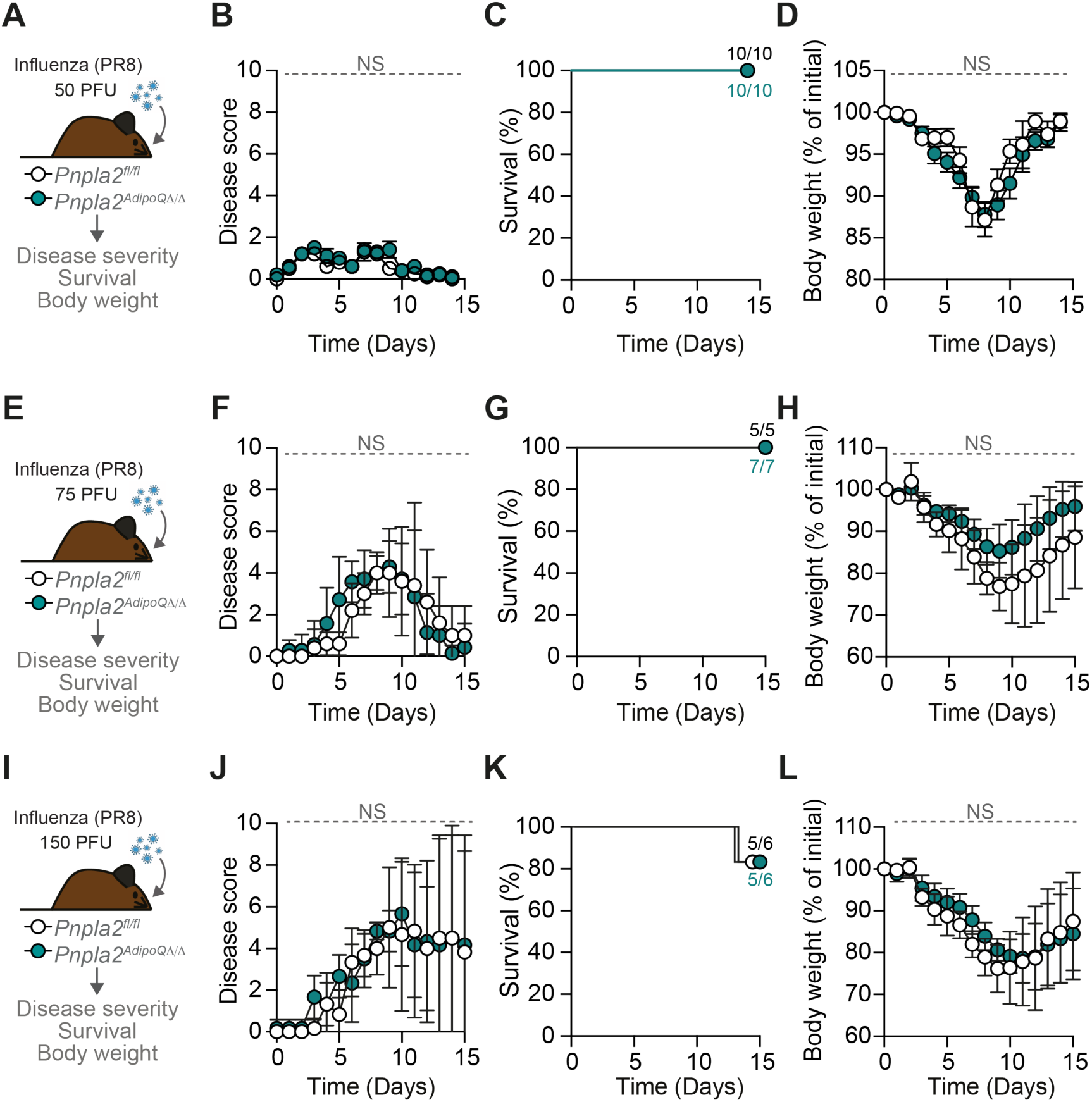
Adipocyte lipolysis is dispensable for disease tolerance to influenza A virus infection. (**A**) Young *Pnpla2^fl/fl^* and *Pnpla2^AdipoQΔ/Δ^* mice were infected with 50 PFU of influenza A virus (PR8 strain) and monitored for (**B**) disease severity, (**C**) survival and (**D**) body weight (n= 10 mice *per* group, males & females). (**E**) Young *Pnpla2^fl/fl^* and *Pnpla2^AdipoQΔ/Δ^*mice were infected with 75 PFU of influenza A virus and monitored for (**F**) disease severity, (**G**) survival and (**H**) body weight (n= 5-7 mice *per* group, females). (**I**) Young *Pnpla2^fl/fl^* and *Pnpla2^AdipoQΔ/Δ^* mice were infected with 150 PFU of influenza A virus and monitored for (**J**) disease severity, (**K**) survival and (**L**) body weight (n= 5-6 mice *per* group, females). Numbers in (C, G, K) indicate surviving mice per total mice at day 14. Data shown as mean ± SD. *p* values determined using two-way ANOVA (B, D, F, H, J, L) or Log-rank test (C, G, K). NS, not significant. Data pooled from two independent experiments with a similar trend.

**Supplementary Figure 7.**
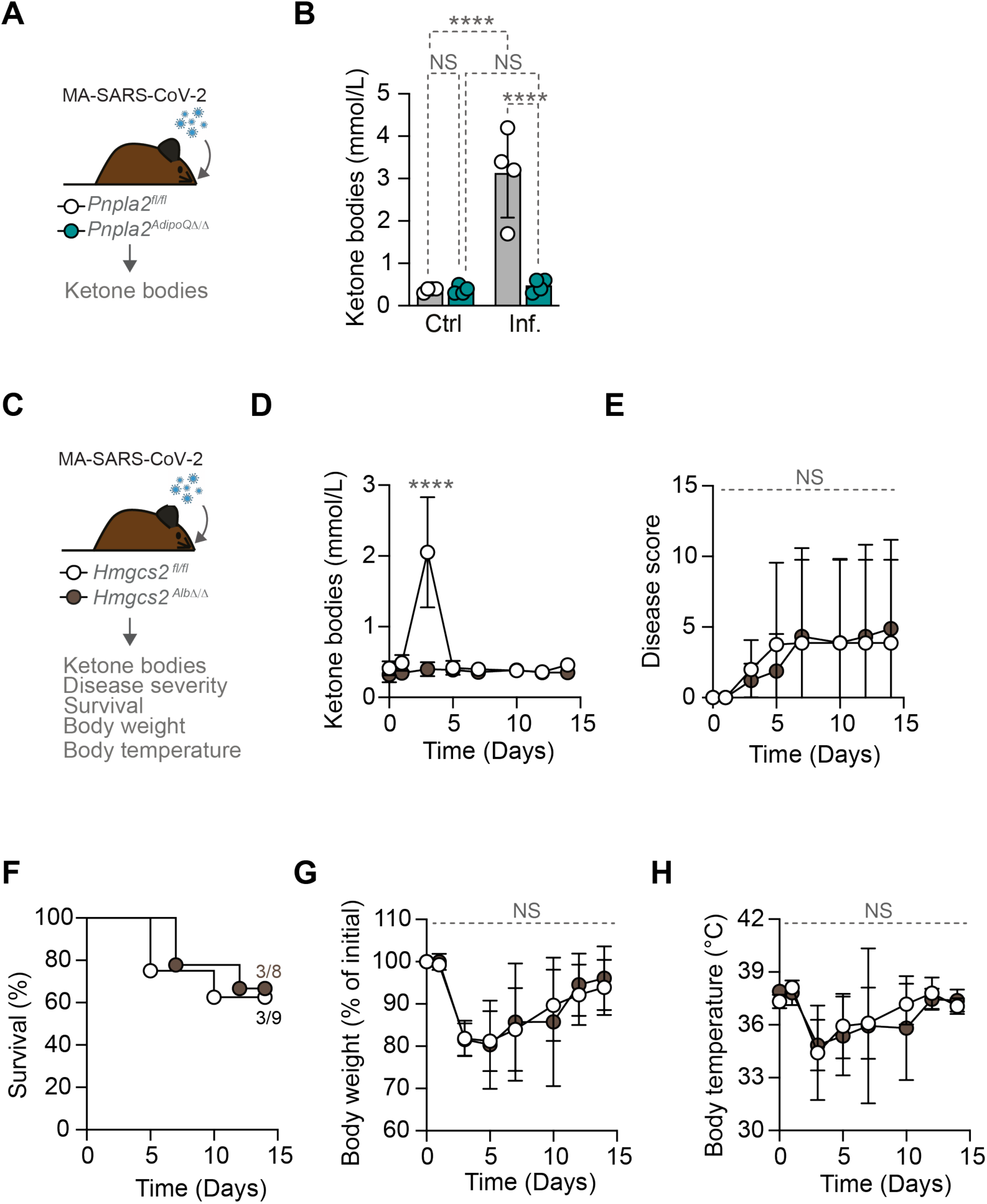
Hepatic ketogenesis is not essential to establish disease tolerance to MA-SARS-CoV-2 infection. (**A**) Young *Pnpla2^fl/fl^* and *Pnpla2^AdipoQΔ/Δ^* mice were infected with MA-SARS-CoV-2 and monitored for (**B**) blood ketone bodies at 3 dpi (n=4 mice *per* group, females). (**C**) *Hmgcs2^fl/fl^*and *Hmgcs2^AlbΔ/Δ^* mice were infected with MA-SARS-CoV-2 and monitored for (**D**) blood ketone bodies, (**E**) disease severity, (**F**) survival, (**G**) body weight and (**H**) body temperature, over 14 dpi. (N=8-9 mice *per* group; males & females; numbers in (F) indicate surviving mice per total mice at day 14). Data shown as mean ± SD (B, D, E, F, G, H). *p* values determined by two-way ANOVA (B, D-H) or Log-rank test (F). NS, not significant; ****p < 0.0001. Data pooled from three independent experiments with a similar trend (D-H) or from one experiment (B).

**Supplementary Figure 8.**
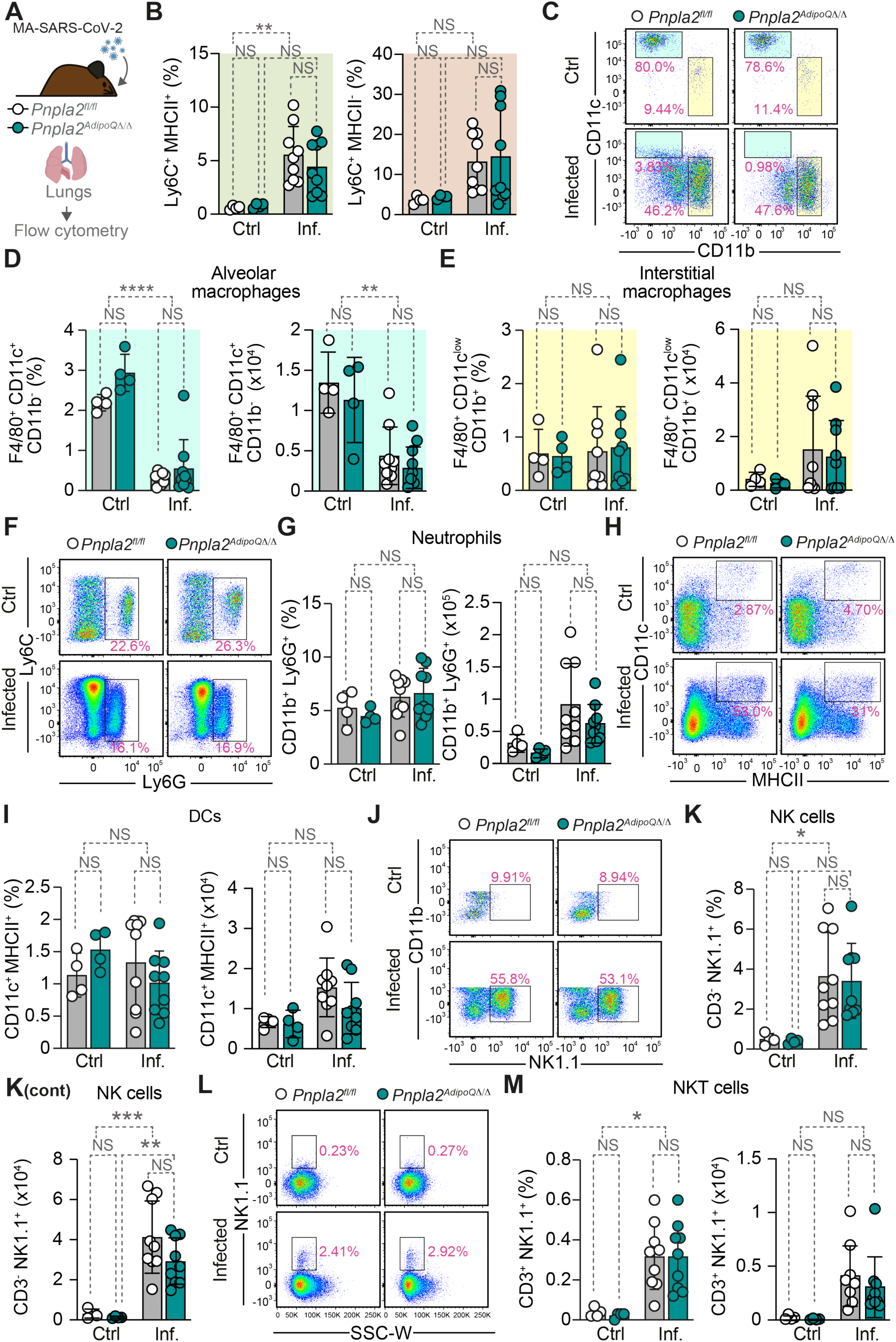
ATGL deletion selectively impairs monocyte recruitment without affecting other immune cell populations in infected lungs. (**A**) Young *Pnpla2^fl/fl^* and *Pnpla2^AdipoQΔ/Δ^*mice were infected with MA-SARS-CoV-2 and lung immune cell populations were analyzed by flow cytometry at 5 dpi. (**B**) Percentage of Ly6C^+^ MHCII^+^ and Ly6C^+^ MHCII^−^ monocytes. (**C**) Representative flow cytometry plots showing CD11b *vs.* CD11c gating for macrophage populations. Percentage and absolute numbers of (**D**) alveolar macrophages (F4/80^+^CD11c^+^CD11b^−^) and (**E**) interstitial macrophages (F4/80^+^CD11c^low^CD11b^+^). (**F**) Representative flow cytometry plots showing Ly6G *vs.* Ly6C gating for neutrophils. (**G**) Percentage and absolute numbers of neutrophils (CD11b^+^Ly6G^+^). (**H**) Representative flow cytometry plots showing MHCII *vs.* CD11c gating for dendritic cells. (**I**) Percentage and absolute numbers of dendritic cells (CD11c^+^MHCII^+^). (**J**) Representative flow cytometry plots showing NK1.1 *vs.* CD11b gating for NK cells. (**K**) Percentage and absolute numbers of NK cells (CD3^−^NK1.1^+^). (**L**) Representative flow cytometry plots showing SSC-W *vs.* NK1.1 gating for NKT cells. (**M**) Percentage and absolute numbers of NKT cells (CD3*^+^*NK1.1^+^). Data represented as mean ± SD (B, D, E, G, I, K, M). Circles represent individual mice (n=4-9 mice per group, males & females). *p* values determined using two-way ANOVA. NS, not significant; *p < 0.05; **p < 0.01; ***p < 0.001; ****p < 0.0001. Data pooled from two independent experiments with a similar trend.

**Supplementary Figure 9.**
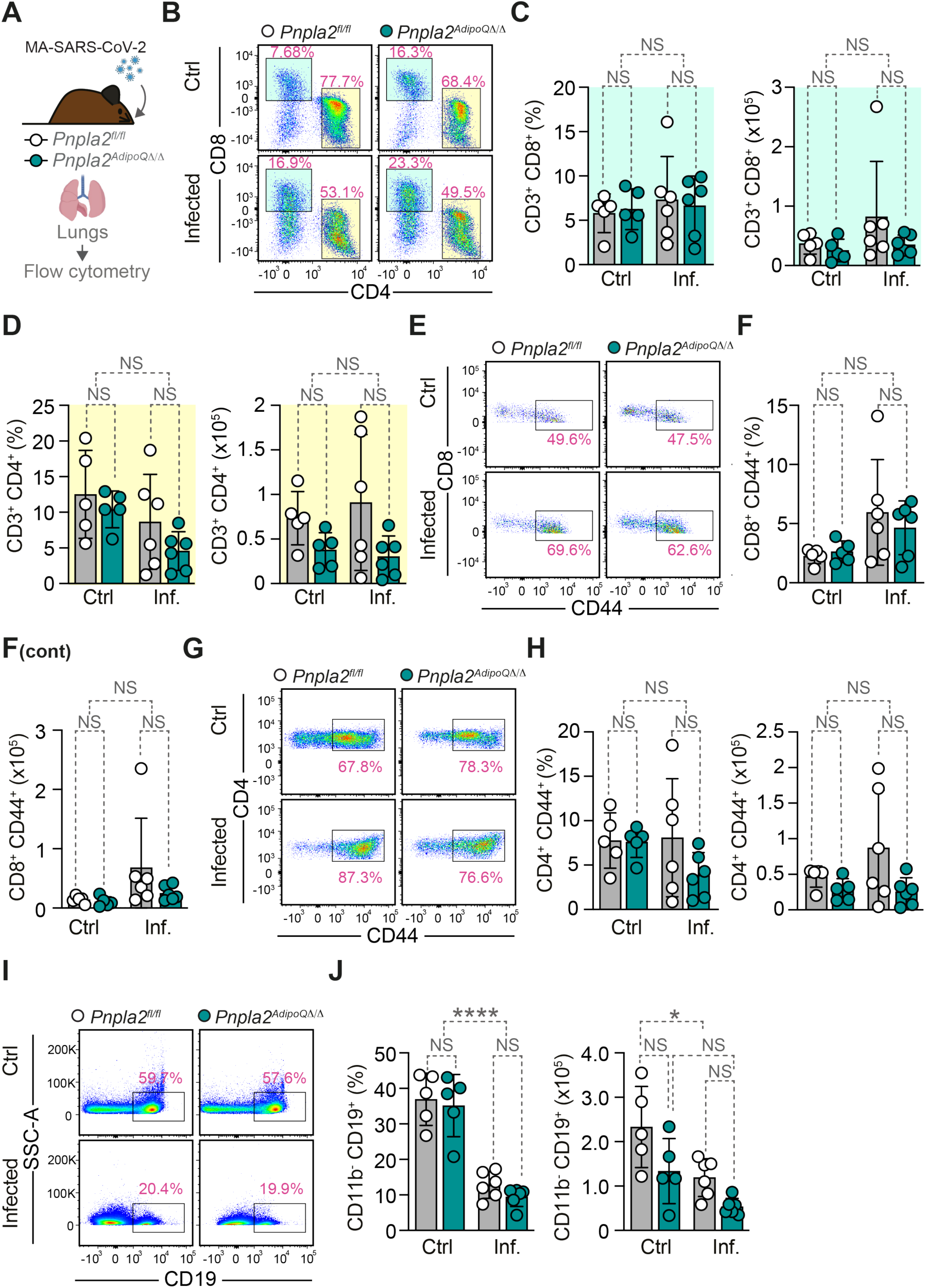
ATGL deletion does not impair adaptive immune response in infected lungs. (**A**) *Pnpla2^fl/fl^* and *Pnpla2^AdipoQΔ/Δ^* mice were infected with MA-SARS-CoV-2 and lung immune cell populations were analyzed by flow cytometry at 7 dpi. (**B**) Representative flow cytometry plots showing CD4 *vs.* CD8 gating for T cells population. **(C)** Percentage and absolute numbers of CD8^+^ T cells (CD3^+^CD8^+^) and (**D**) CD4^+^ T cells (CD3^+^CD4^+^). (**E)** Representative flow cytometry plots showing CD44 *vs.* CD8 gating for activated CD8 T cells population. (**F**) Percentage and absolute numbers of activated CD8 T cells (CD8^+^CD44^+^). (**G**) Representative flow cytometry plots showing CD44 *vs.* CD4 gating for activated CD4 T cells population. (**H**) Percentage and absolute numbers of activated CD4 T cells (CD4^+^CD44^+^). (**I**) Representative flow cytometry plots showing CD19 *vs.* SSC-A gating on CD11b^−^ for total B cells population. (**J**) Percentage and absolute numbers of B cells (CD11b^−^CD19^+^). Data in (C, D, F, H, J) represented as mean *±* SD. Circles represent individual mice (n=5-6 mice *per* group, males & females). *p* values determined using two-way ANOVA. NS, not significant; *p < 0.05; ****p < 0.0001. Data pooled from three independent experiments with a similar trend.

**Supplementary Figure 10.**
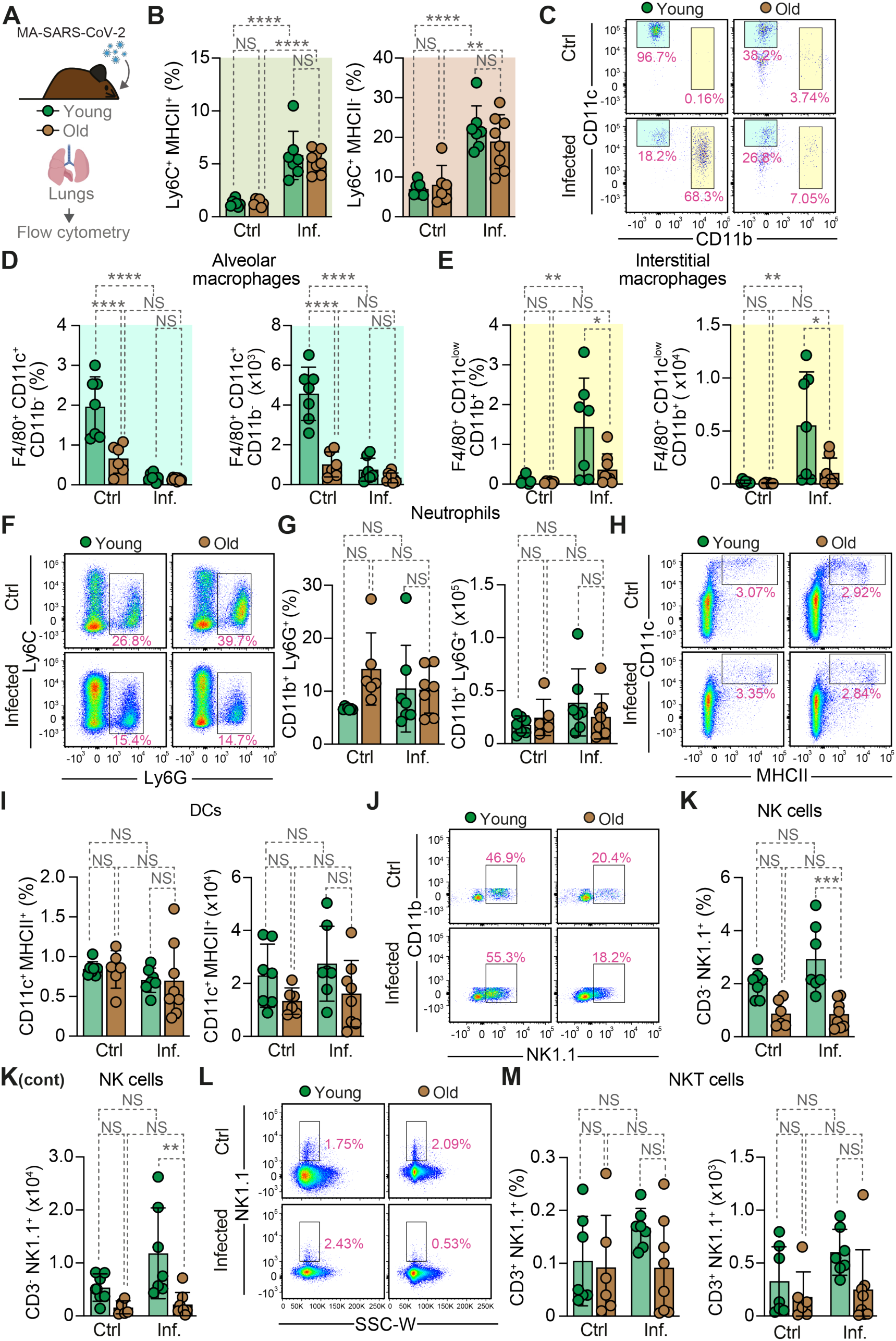
Age-dependent disease tolerance failure is associated with innate immune recruitment defects in the lung. (**A**) Young (8-12 weeks) and old (>18 months) C57BL/6J mice were infected with MA-SARS-CoV-2 and lung immune cell populations were analyzed by flow cytometry at 5 dpi. (**B**) Percentage of Ly6C^+^ MHCII^+^ and Ly6C^+^ MHCII^−^ monocytes. (**C**) Representative flow cytometry plots showing CD11b *vs.* CD11c gating for macrophage populations. Percentage and absolute numbers of (**D**) alveolar macrophages (F4/80^+^CD11c^+^CD11b^−^) and (**E**) interstitial macrophages (F4/80^+^CD11c^low^CD11b^+^). (**F**) Representative flow cytometry plots showing Ly6G *vs.* Ly6C gating for neutrophils. (**G**) Percentage and absolute numbers of neutrophils (CD11b^+^Ly6G^+^). (**H**) Representative flow cytometry plots showing MHCII *vs.* CD11c gating for dendritic cells. (**I**) Percentage and absolute numbers of dendritic cells (CD11c^+^MHCII^+^). (**J**) Representative flow cytometry plots showing NK1.1 *vs.* CD11b gating for NK cells. (**K**) Percentage and absolute numbers of NK cells (CD3^−^NK1.1^+^). (**L**) Representative flow cytometry plots showing SSC-W *vs.* NK1.1 gating for NKT cells. (**M**) Percentage and absolute numbers of NKT cells (CD3*^+^*NK1.1^+^). Data represented as mean *±* SD (B, D, E, G, I, K, M). Circles represent individual mice (n=6-8 mice *per* group, males). *p* values were determined using two-way ANOVA. NS, not significant; *p < 0.05; **p < 0.01; ***p < 0.001; ****p < 0.0001. Data pooled from two independent experiments with a similar trend.

**Supplementary Figure 11.**
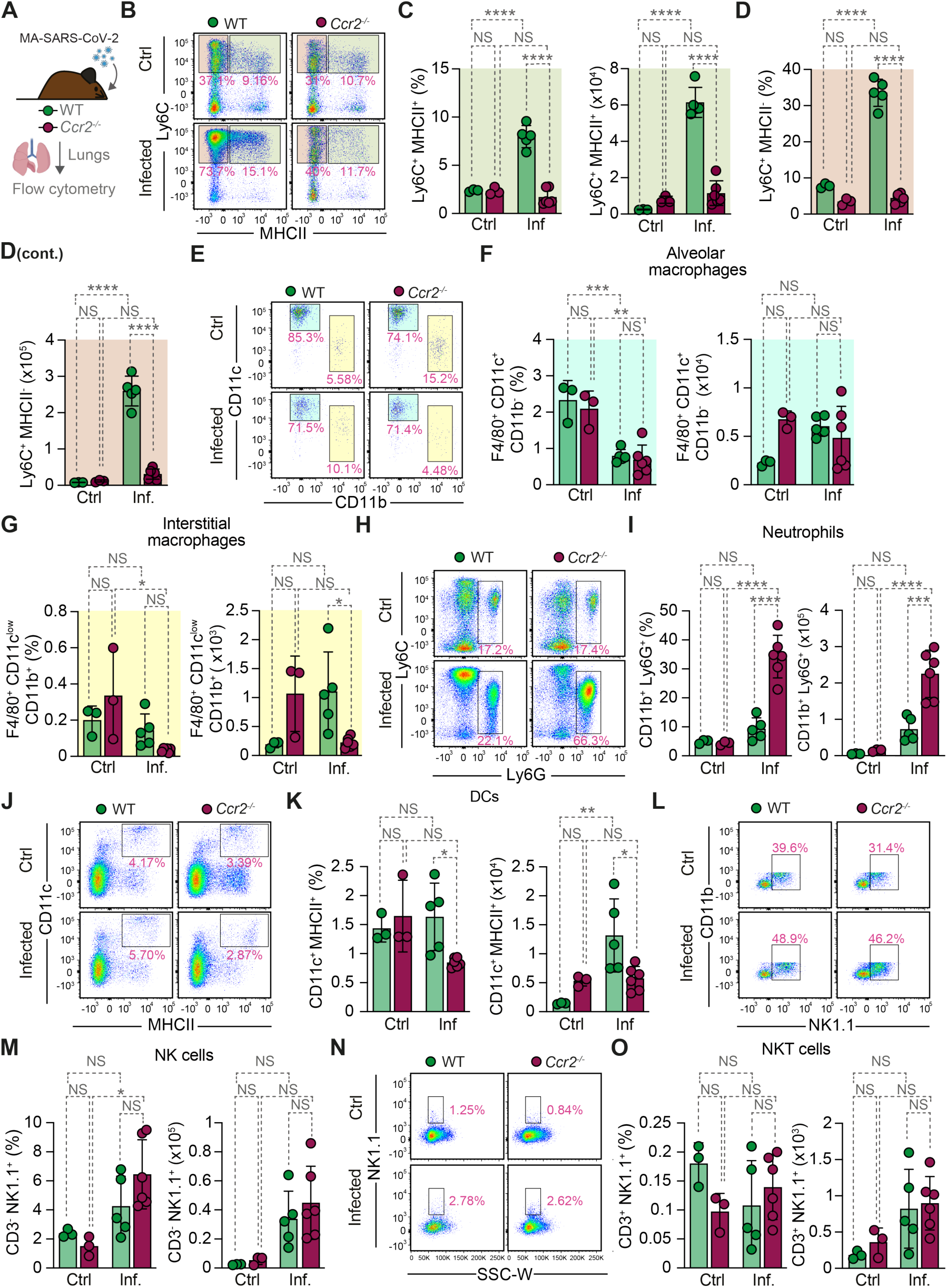
CCR2 deficiency impairs monocytes recruitment to the lungs during MA-SARS-CoV-2 infection. (A) WT and *Ccr2*^−/−^ mice were infected with MA-SARS-CoV-2 and lung immune cell populations were analyzed by flow cytometry at 5 dpi. (**B**) Representative flow cytometry plots showing MHCII *vs.* Ly6C gating on CD11b^+^ cells. (**C**) Percentage and absolute numbers of Ly6C^+^MHCII^+^ monocytes and (**D**) Ly6C^+^MHCII^−^ monocytes. (**E**) Representative flow cytometry plots showing CD11b *vs.* CD11c gating for macrophage populations. (**F**) Percentage and absolute numbers of alveolar macrophages (F4/80^+^CD11c^+^CD11b^−^) and (**G**) interstitial macrophages (F4/80^+^CD11c^low^CD11b^+^). **(H)** Representative flow cytometry plots showing Ly6G *vs.* Ly6C gating for neutrophils. (**I**) Percentage and absolute numbers of neutrophils (CD11b+Ly6G+). **(J)** Representative flow cytometry plots showing MHCII *vs.* CD11c gating for dendritic cells. (**K**) Percentage and absolute numbers of dendritic cells (CD11c+MHCII+). (**L**) Representative flow cytometry plots showing NK1.1 *vs.* CD11b gating for NK cells. (**M**) Percentage and absolute numbers of NK cells (CD3^−^NK1.1^+^). (**N**) Representative flow cytometry plots showing SSC-W *vs.* NK1.1 gating for NKT cells. (**O**) Percentage and absolute numbers of NKT cells (CD3^+^NK1.1^+^). Data are shown as mean ± SD (C, D, F, G, I, K, M, O). Circles represent individual mice (n=3-6 *per* group, males & females). *p* values were determined using two-way ANOVA. NS, not significant; *p < 0.05; **p < 0.01; ***p < 0.001; ****p < 0.0001. Data pooled from two independent experiments with a similar trend.

**Supplementary Figure 12.**
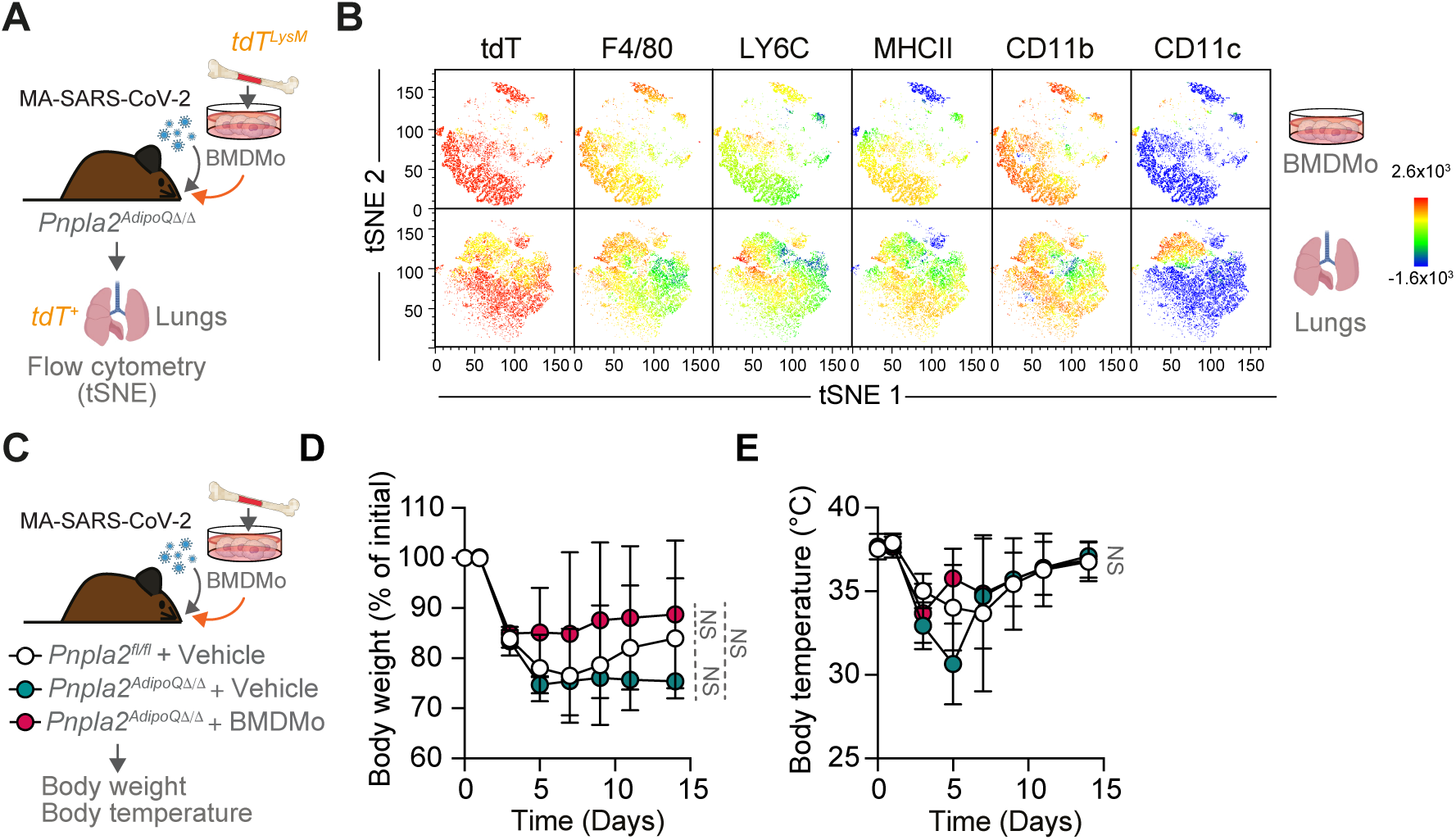
Adoptive transfer of BM derived monocytes rescues *Pnpla2^AdipoQΔ/Δ^* mice from severe SARS-CoV-2 infection. (**A**) *Pnpla2^AdipoQΔ/Δ^* mice were infected with MA-SARS-CoV-2 and received adoptive transfer of bone marrow-derived monocytes (BMDMo) from *LysM^Cre/WT^ tdT^R^*^26^*(tdT^LysM^)* mice (N=4 mice, males). **(B)** tSNE plots showing marker expression profiles in control *tdT^LysM^* BMDMo as well as tdTomato⁺ cells recovered from the lungs of *Pnpla2^AdipoQΔ/Δ^* mice. Each plot displays the expression of the designated marker across the tSNE projection. tSNE was used to generate projections from 16000 cells (downsampled) for BMDMo prior to adoptive transfer, and from 3900 to 4000 tdTomato^+^ cells/mice for BMDMo recovered from the lungs following adoptive transfer (pooled from 4 mice). (**C**) *Pnpla2^fl/fl^* and *Pnpla2^AdipoQΔ/Δ^*mice were infected with MA-SARS-CoV-2. *Pnpla2^AdipoQΔ/Δ^* mice received either vehicle or adoptive transfer of BMDMo from C57BL6/J mice, monitored for (**D**) body weight and (**E**) body temperature over 14 dpi. (N=6-9 mice *per* group, males). Data shown as mean ± SD (D, E). *p* values determined by two-way ANOVA (D,E). NS, not significant; **p < 0.01. Data pooled from two independent experiments with a similar trend (D, E) or from one experiment (B).

**Supplementary Figure 13.**
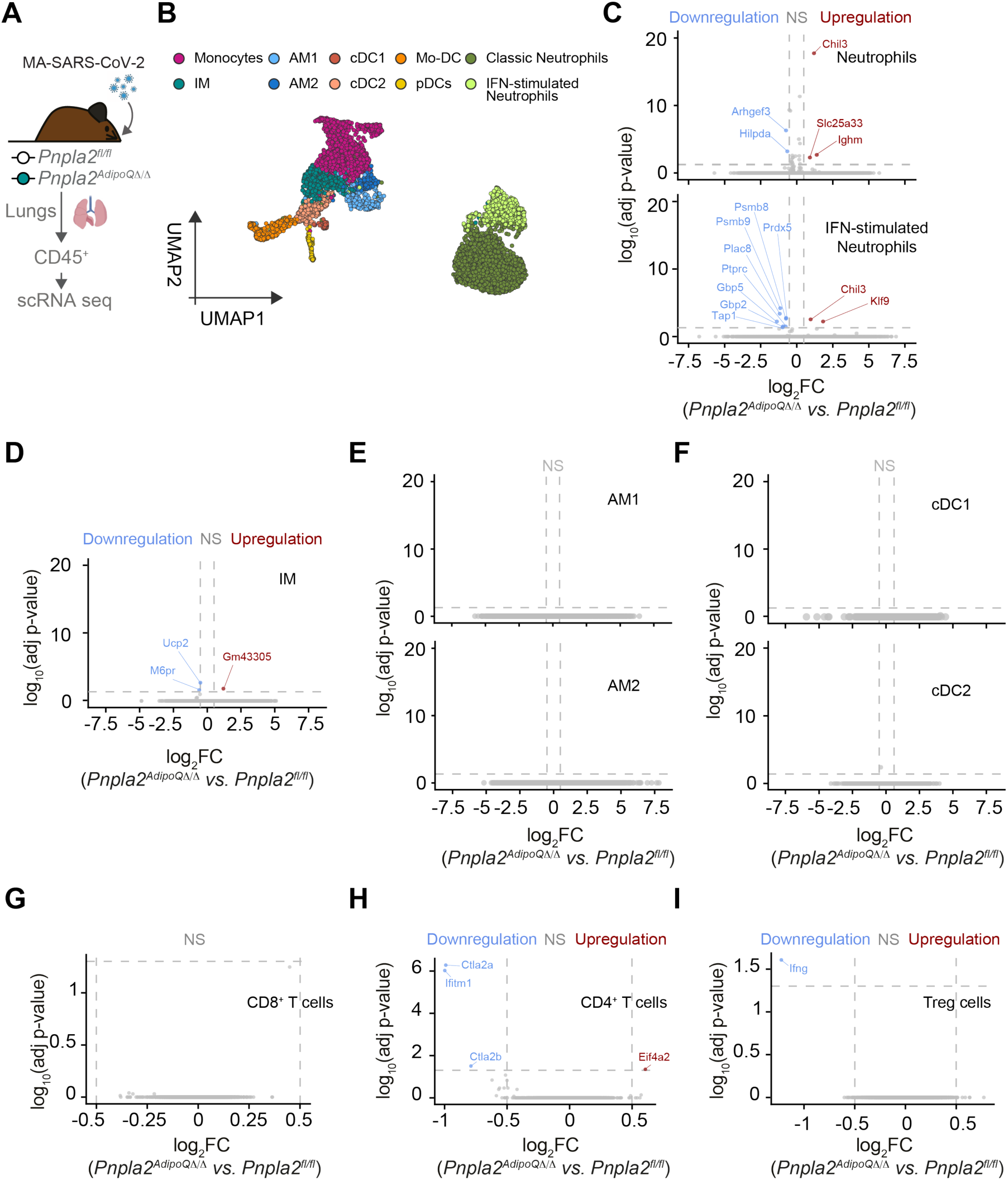
Adipose tissue lipolysis regulates monocyte transcriptional responses and cell-cell communication during MA-SARS-CoV-2 infection. (**A**) Young *Pnpla2^fl/fl^* and *Pnpla2^AdipoQΔ/Δ^*male mice were infected with MA-SARS-CoV-2 and lungs were harvested at 5 dpi. CD45^+^ cells were analysed for scRNA-seq (n=3 mice *per* group). (**B**) UMAP of myeloid cell from CD45^+^ lung cells, classified as monocytes, interstitial macrophages (IM), alveolar macrophages (AM1 and AM2), conventional dendritic cells (cDC1, cDC2), monocyte-derived dendritic cells (Mo-DC), plasmacytoid dendritic cells (pDC), classic neutrophils, and IFN-stimulated neutrophils. Volcano plots showing differential gene expression (log₂FC ≥ 0.5, *p*_adj_ ≤ 0.05) in **(C)** classic neutrophils (top) and IFN-stimulated neutrophils (bottom), (**D**) IM, (**E**) AM1 (top) and AM2 (bottom), (**F**) cDC1 (top) and cDC2 (bottom) (**G**) CD4^+^ T cells, (**H**) CD8^+^ T cells and (**I**) regulatory T cells (Treg) from *Pnpla2^AdipoQΔ/Δ^ vs. Pnpla2^fl/fl^* infected mice. Differentially expressed genes (DEG) are highlighted in red (upregulated) or blue (downregulated), non-DEG (NS) are shown in grey. Each dot represents an individual gene.

**Supplementary Figure 14.**
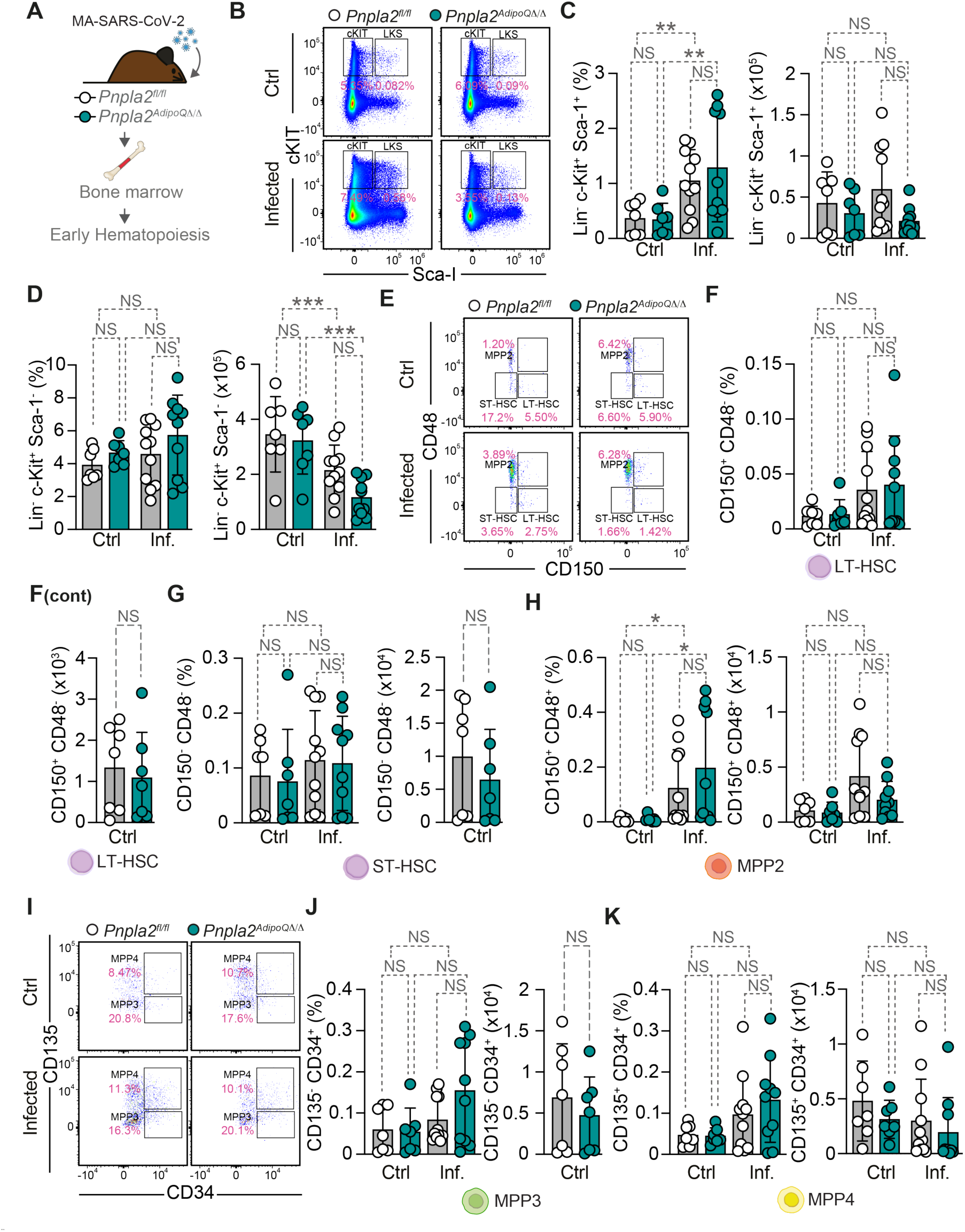
Characterization of early BM hematopoietic stem and progenitor cell populations during MA-SARS-CoV-2 infection upon ATGL deficiency. (**A**) Young *Pnpla2^fl/fl^* and *Pnpla2^AdipoQΔ/Δ^* mice were infected with MA-SARS-CoV-2 and BM was analyzed for early hematopoiesis at 5 dpi. (**B**) Representative flow cytometry plots showing c-KIT *vs*. Sca-1 gating on lineage-negative (Lin^−^) cells to identify LKS (Lin^−^c-Kit^+^Sca-1^+^) hematopoietic stem and progenitor cells. Percentage and absolute numbers of (**C**) LKS (Lin^−^ cKIT^+^ Sca-1^+^) and (**D**) c-KIT^+^ (Lin^−^ cKIT^+^ Sca-1^−^) cells. **(E)** Representative flow cytometry plots showing CD48 *vs.* CD150 gating on LKS cells to identify long-term hematopoietic stem cells (LT-HSC: CD150^+^CD48^−^), short-term HSC (ST-HSC: CD150^−^CD48^−^) and multipotent progenitor 2 (MPP2: CD150^+^ CD48^+^). Percentage and absolute numbers of (**F**) LT-HSC, (**G**) ST-HSC and (**H**) MPP2 cells. (**I**) Representative flow cytometry plots showing CD135 *vs*. CD34 gating on MPP cells to identify MPP3 (CD135^−^CD34^+^) and MPP4 (CD135^+^CD34^+^). Percentage and absolute numbers of **(J)** MPP3 and **(K)** MPP4 cells. Data shown as mean ± SD (C, D, F, G, H, J, K). Circles represent individual mice (n=7-11 *per* group, males & females). *p* values determined using two-way ANOVA. NS, not significant; *p < 0.05. Data pooled from three independent experiments with a similar trend.

**Supplementary Figure 15.**
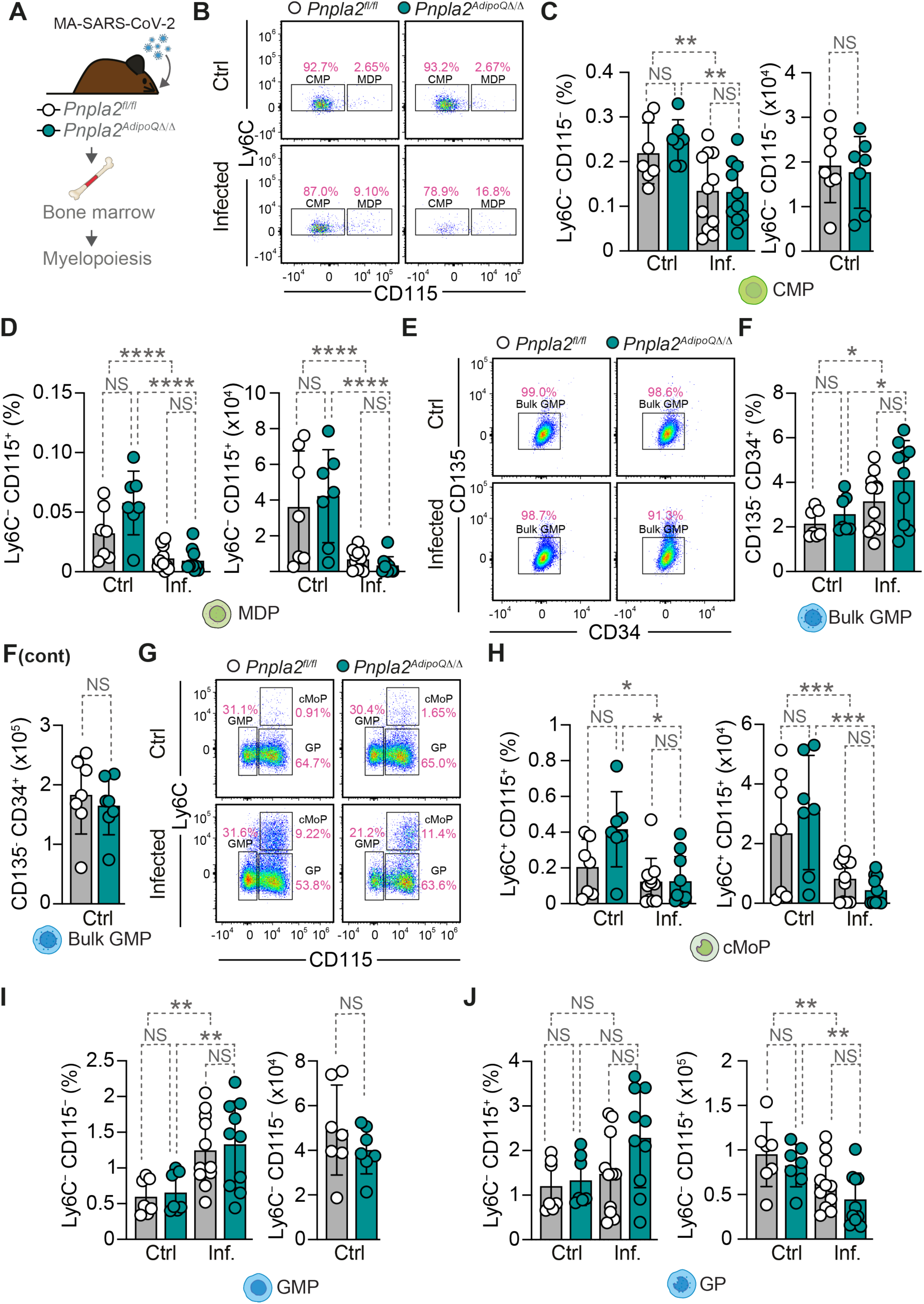
Characterization of myeloid progenitors during MA-SARS-CoV-2 infection upon ATGL deficiency. (**A**) Young *Pnpla2^fl/fl^* and *Pnpla2^AdipoQΔ/Δ^* mice were infected with MA-SARS-CoV-2 and BM was analyzed for myelopoiesis at 5 dpi. (**B**) Representative flow cytometry plots showing Ly6C *vs.* CD115 gating on Lin^−^c-Kit^+^Sca-1^−^ cells to identify common myeloid progenitors (CMP: Ly6C^−^CD115^−^) and monocyte-dendritic cell progenitors (MDP: Ly6C^−^CD115^+^). Percentage and absolute numbers of (**C**) CMP and (**D**) MDP cells. (**E**) Representative flow cytometry plots showing CD135 *vs.* CD34 gating on CD16/32^+^ to identify bulk granulocyte-monocyte progenitors (Bulk GMP: CD135^−^CD34^+^). Percentage and absolute numbers of (**F**) bulk GMP cells. (**G**) Representative flow cytometry plots showing Ly6C *vs.* CD115 gating on bulk GMP to identify granulocyte-monocyte progenitors (GMP: Ly6C^−^CD115^−^), common monocyte progenitors (cMoP: Ly6C^+^ CD115^+^) and granulocyte progenitors (GP: Ly6C^−^ CD115^+^). Percentage and absolute numbers of (**H**) cMoP, (**I**) GMP and (**J**) GP. Data are shown as mean ± SD (C, D, F, H, I, J). Circles represent individual mice (n=7-11 *per* group, males & females). *p* values were determined using two-way ANOVA. NS, not significant; *p < 0.05; **p < 0.01; ***p < 0.001, ****p < 0.0001. Data pooled from three independent experiments with a similar trend.

**Supplementary Figure 16.**
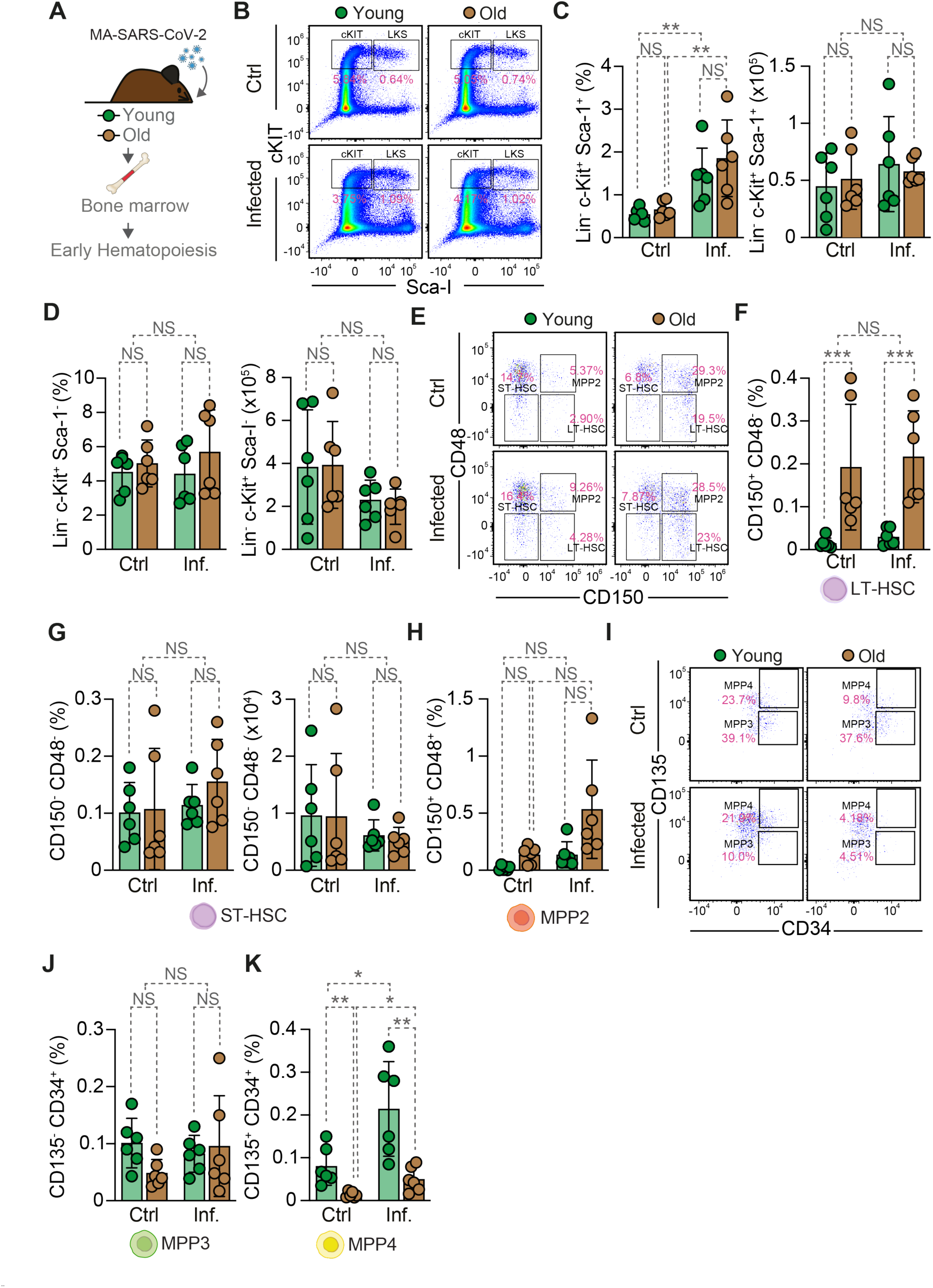
Aging selectively alters early hematopoietic stem and progenitor cell populations during MA-SARS-CoV-2 infection. (**A**) Young (3 months) and old (>20 months) mice were infected with MA-SARS-CoV-2 and BM was analyzed for early hematopoiesis at 5 dpi. (**B**) Representative flow cytometry plots showing c-KIT *vs.* Sca-1 gating on lineage-negative (Lin^−^) cells to identify LKS (Lin^−^c-Kit^+^Sca-1^+^) hematopoietic stem and progenitor cells. Percentage and absolute numbers of (**C**) LKS cells and (**D**) c-KIT^+^ (Lin^−^ cKIT^+^ Sca-1^−^) cells. (**E**) Representative flow cytometry plots showing CD48 *vs.* CD150 gating on LKS cells to identify long-term hematopoietic stem cells (LT-HSC: CD150^+^CD48^−^), short-term HSC (ST-HSC: CD150^−^CD48^−^), and multipotent progenitor 2 (MPP2: CD150^+^CD48^+^). Percentage and absolute numbers of (**F**) LT-HSC, (**G**) ST-HSC and (**H**) MPP2 cells. (**I**) Representative flow cytometry plots showing CD135 *vs.* CD34 gating on MPP cells to identify MPP3 (CD135^−^CD34^+^) and MPP4 (CD135^+^CD34^+^). Percentage and absolute numbers of (**J**) MPP3 and (**K**) MPP4 cells. Data are shown as mean ± SD (C, D, F, G, H, J, K). Circles represent individual mice (n=6 *per* group, males). *p* values were determined using two-way ANOVA. NS, not significant; *p < 0.05, **p < 0.01; ***p < 0.001. Data pooled from two independent experiments with a similar trend.

**Supplementary Figure 17.**
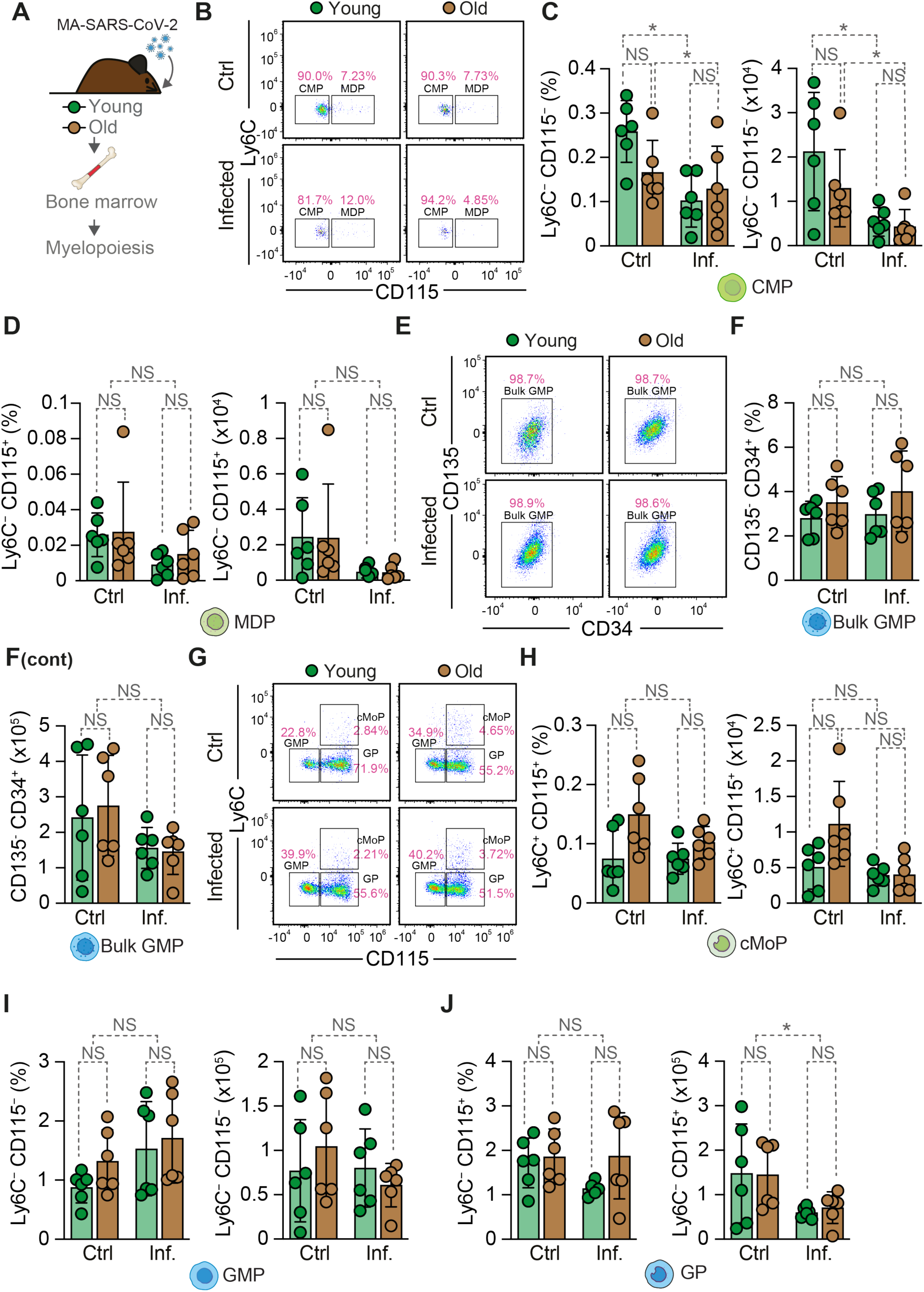
Age-dependent BM myeloid progenitor characterization during MA-SARS-CoV-2 infection. (**A**) Young (3 months) and old (>20 months) mice were infected with MA-SARS-CoV-2 and BM was analyzed for myelopoiesis at 5 dpi. (**B**) Representative flow cytometry plots showing Ly6C *vs*. CD115 gating on Lin^−^c-Kit^+^Sca-1^−^ cells to identify common myeloid progenitors (CMP: Ly6C^−^CD115^−^) and monocyte-dendritic cell progenitors (MDP: Ly6C^−^CD115^+^). Percentage and absolute numbers of (**C**) CMP and (**D**) MDP cells. (**E**) Representative flow cytometry plots showing CD135 *vs.* CD34 gating on CD16/32^+^ cells to identify bulk granulocyte-monocyte progenitors (Bulk GMP: CD135^−^CD34^+^). (**F**) Percentage and absolute numbers of bulk GMP cells. (**G**) Representative flow cytometry plots showing Ly6C *vs.* CD115 gating on bulk GMP to identify granulocyte-monocyte progenitors (GMP: Ly6C^−^CD115^−^), common monocyte progenitors (cMoP: Ly6C^+^CD115^+^), and granulocyte progenitors (GP: Ly6C^−^CD115^+^). Percentage and absolute numbers of (**H**) cMoP, (**I**) GMP and (**J**) GP. Data are shown as mean ± SD (C, D, F, H, I, J). Circles represent individual mice (n=6 per group, males). *p* values were determined using two-way ANOVA. NS, not significant; *p < 0.05. Data pooled from two independent experiments with a similar trend.

**Supplementary Figure 18.**
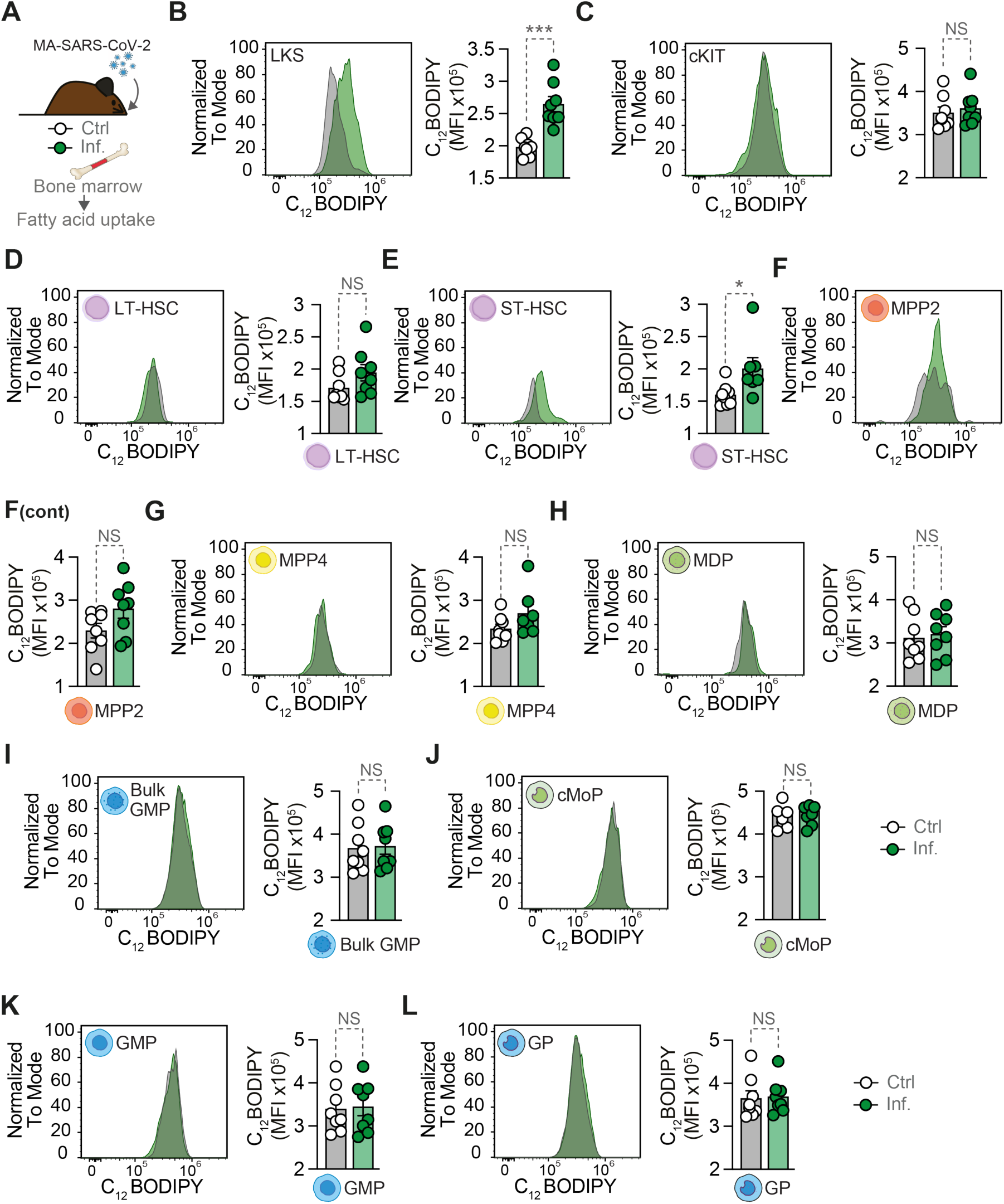
Assessment of FFA uptake by BM progenitor cells during MA-SARS-CoV-2 infection. (**A**) Control and infected C57BL/6J young mice were analyzed for FFA uptake by BM progenitor cells using BODIPY C_12_ at 5 dpi. (**B**) Representative flow cytometry histograms showing BODIPY C_12_ staining (fatty acid uptake) and quantification of BODIPY C_12_ mean fluorescence intensity (MFI) in LKS, (**C**) cKIT, (**D**) LT-HSC, (**E**) ST-HSC, (**F**) MPP2, (**G**) MPP4, (**H**) MDP, (**I**) Bulk GMP, (**J**) cMoP, (**K**) GMP and (**L**) GP populations from control and infected mice (n=8 mice *per* group, males). Data represented as mean *±* SEM (B-L). Circles represent individual mice (B-L). *p* values were determined using unpaired t-test (B-L). NS, not significant; *p < 0.05; ***p < 0.001. Data pooled from two independent experiments with a similar trend.

**Supplementary Figure 19.**
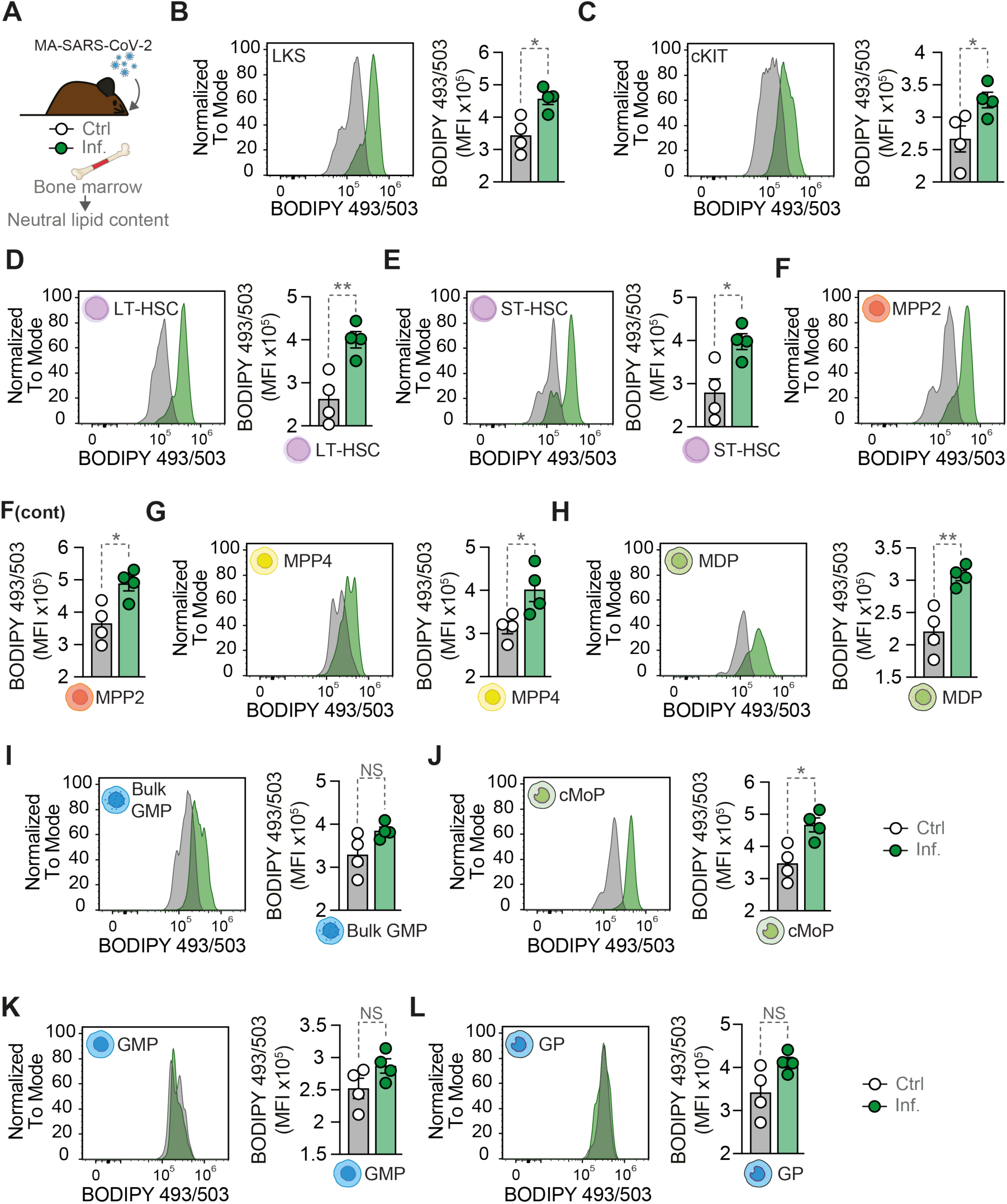
Profiling of intracellular neutral lipid content in BM progenitor cells upon MA-SARS-CoV-2 infection. (**A**) Control and infected C57BL/6J young mice were analyzed for lipid content in BM progenitor cells using BODIPY 493/503 at 5 dpi. (**B**) Representative flow cytometry histograms showing BODIPY 493/503 staining (lipid content) and quantification of BODIPY 493/503 mean fluorescence intensity (MFI) in LKS, (**C**) cKIT, (**D**) LT-HSC, (**E**) ST-HSC, (**F**) MPP2, (**G**) MPP4, (**H**) MDP, (**I**) Bulk GMP, (**J**) cMoP, (**K**) GMP and (**L**) GP populations from control and infected mice (n=4 mice *per* group from one out of two independent experiments with a similar trend, males). Data represented as mean *±* SEM (B-L). Circles represent individual mice (B-L). *p* values were determined using unpaired t-test (B-L). NS, not significant; *p < 0.05; **p < 0.01.

**Supplementary Figure 20.**
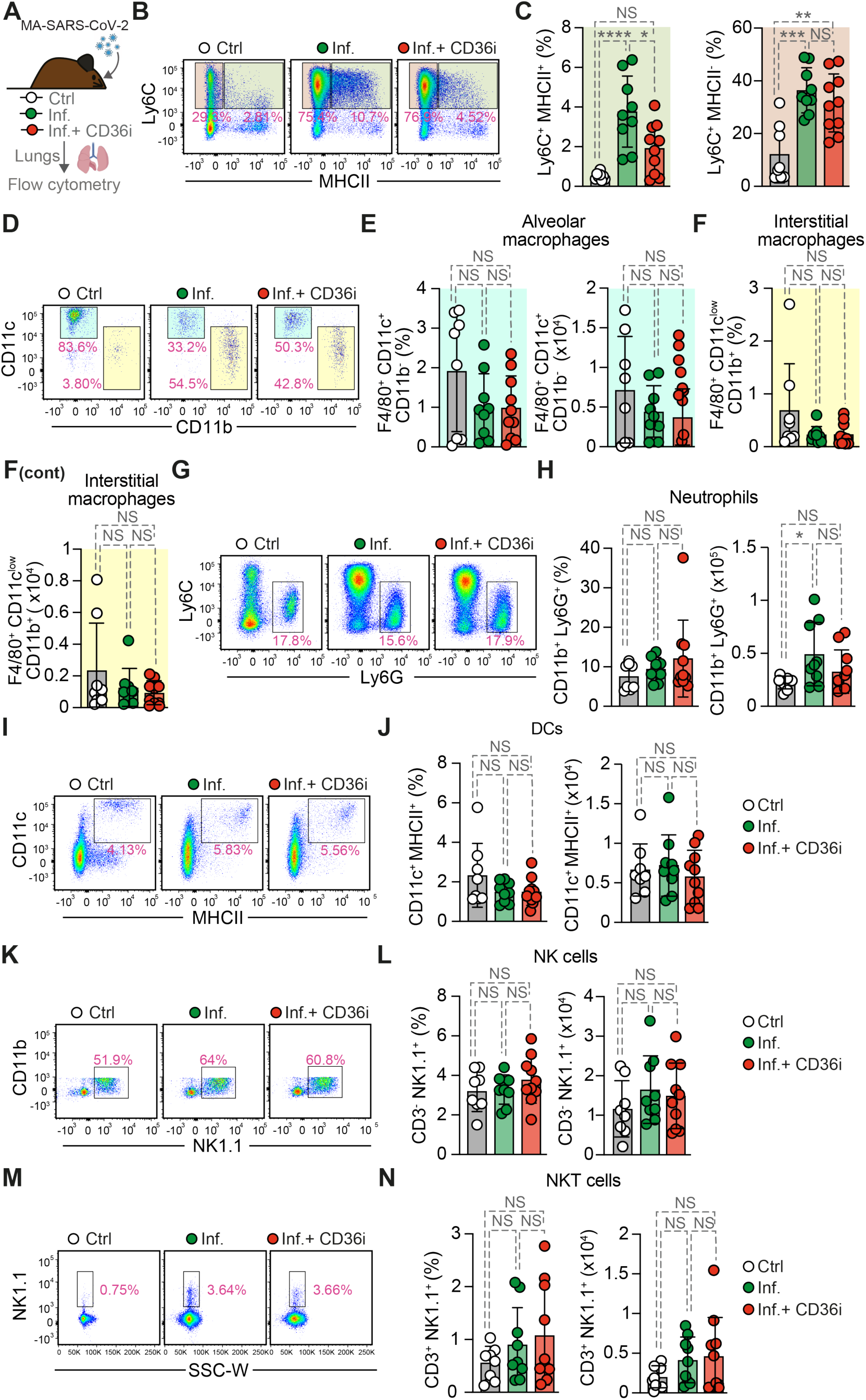
Pharmacologic CD36 inhibition specifically alters monocytes recruitment during MA-SARS-CoV-2 infection. (**A**) C57BL/6J mice were infected with MA-SARS-CoV-2 and treated with CD36 inhibitor, sulfo-N-succinimidyl oleate (CD36i), or vehicle and lung immune cell populations were analyzed by flow cytometry at 5 dpi. (**B**) Representative flow cytometry plots showing MHCII *vs.* Ly6C gating on CD11b^+^ cells. Percentage of (**C**) Ly6C^+^ MHCII^+^ and MHCII^−^ monocytes. (**D**) Representative flow cytometry plots showing CD11b *vs.* CD11c gating for macrophage populations. Percentage and absolute numbers of (**E**) alveolar macrophages (F4/80^+^CD11c^+^CD11b^−^) and (**F**) interstitial macrophages (F4/80^+^CD11c^low^CD11b^+^). (**G**) Representative flow cytometry plots showing Ly6G *vs.* Ly6C gating for neutrophils. (**H**) Percentage and absolute numbers of neutrophils (CD11b^+^Ly6G^+^). (**I**) Representative flow cytometry plots showing MHCII *vs.* CD11c gating for dendritic cells. (**J**) Percentage and absolute numbers of dendritic cells (CD11c^+^MHCII^+^). (**K**) Representative flow cytometry plots showing NK1.1 *vs.* CD11b gating for NK cells. (**L**) Percentage and absolute numbers of NK cells (CD3^−^NK1.1^+^). (**M**) Representative flow cytometry plots showing SSC-W *vs.* NK1.1 gating for NKT cells. (**N**) Percentage and absolute numbers of NKT cells (CD3^+^NK1.1^+^). Data are shown as mean ± SD (C, E, F, H, J, L, N). Circles represent individual mice (n=8-10 *per* group, males). *p* values were determined using one-way ANOVA. NS, not significant; *p < 0.05; **p < 0.01, ***p < 0.001, ****p < 0.0001. Data pooled from three independent experiments with a similar trend.

**Supplementary Figure 21.**
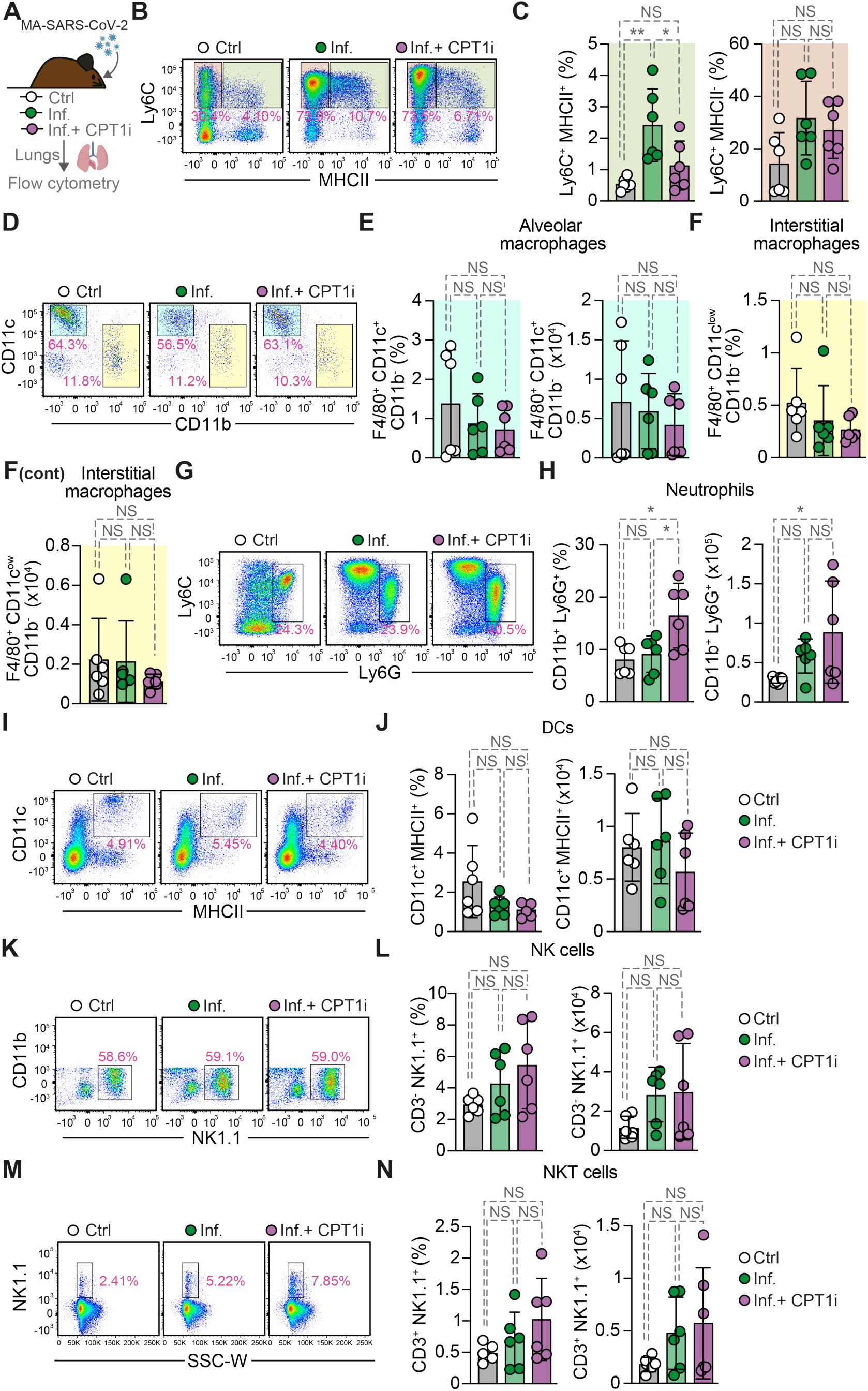
Pharmacologic CPT1 inhibition alters monocytes recruitment during MA-SARS-CoV-2 infection. (**A**) C57BL/6J mice were infected with MA-SARS-CoV-2 and treated with CPT1 inhibitor, etomoxir (CPT1i), or vehicle and lung immune cell populations were analyzed by flow cytometry at 5 dpi. (**B**) Representative flow cytometry plots showing MHCII *vs.* Ly6C gating on CD11b^+^ cells. Percentage of (**C**) Ly6C^+^ MHCII^+^ and MHCII^−^ monocytes. (**D**) Representative flow cytometry plots showing CD11b *vs.* CD11c gating for macrophage populations. Percentage and absolute numbers of (**E**) alveolar macrophages (F4/80^+^CD11c^+^CD11b^−^) and (**F**) interstitial macrophages (F4/80^+^CD11c^low^CD11b^+^). (**G**) Representative flow cytometry plots showing Ly6G *vs.* Ly6C gating for neutrophils. (**H**) Percentage and absolute numbers of neutrophils (CD11b^+^Ly6G^+^). (**I**) Representative flow cytometry plots showing MHCII *vs.* CD11c gating for dendritic cells. (**J**) Percentage and absolute numbers of dendritic cells (CD11c^+^MHCII^+^). (**K**) Representative flow cytometry plots showing NK1.1 *vs.* CD11b gating for NK cells. (**L**) Percentage and absolute numbers of NK cells (CD3^−^NK1.1^+^). (**M**) Representative flow cytometry plots showing SSC-W *vs.* NK1. gating for NKT cells. (**N**) Percentage and absolute numbers of NKT cells (CD3^+^NK1.1^+^). Data are shown as mean ± SD (C, E, F, H, J, L, N). Circles represent individual mice (n=6 *per* group, males). *p* values were determined using one-way ANOVA. NS, not significant; *p < 0.05; **p < 0.01. Data pooled from two independent experiments with a similar trend.

**Supplementary Figure 22.**
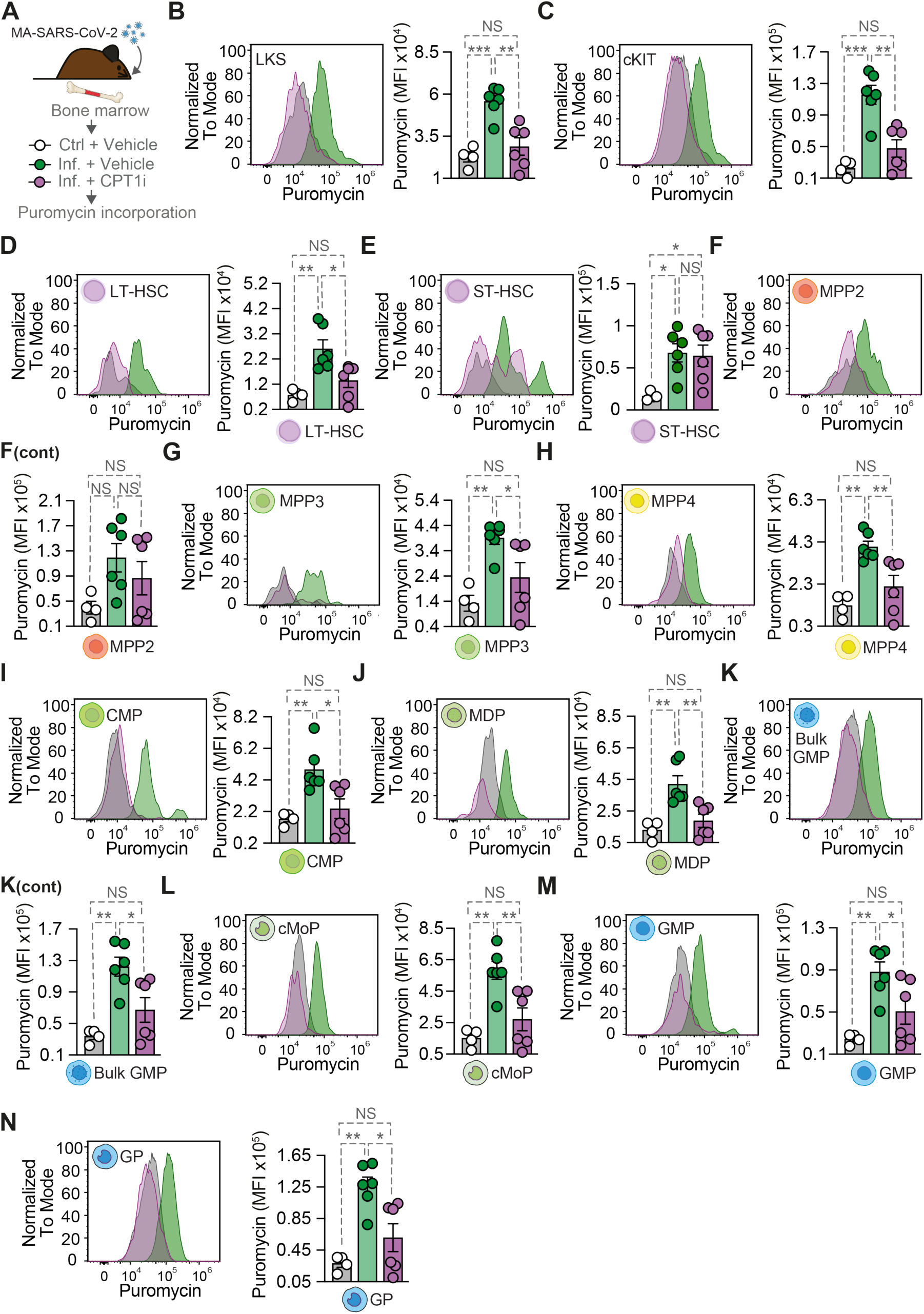
CPT1-dependent fatty acid oxidation fuels BM hematopoietic progenitors during MA-SARS-CoV-2 infection. **(A)** C57BL/6J mice were infected with MA-SARS-CoV-2 and BM progenitor cells were harvested from control and infected mice at 5 dpi. The cells were treated with vehicle or CPT1i (Inf + CPT1i) and analyzed for puromycin incorporation using flow cytometry. (**B**) Representative flow cytometry histograms showing puromycin incorporation levels and quantification of puromycin mean fluorescence intensity (MFI) in LKS, (**C**) cKIT, (**D**) LT-HSC, (**E**) ST-HSC, (**F**) MPP2, (**G**) MPP3, (**H**) MPP4, (**I**) CMP, (**J**) MDP, (**K**) Bulk GMP, cMoP, (**M**) GMP and (**N**) GP populations from control and infected mice treated with vehicle or CPT1i (n=4-6 mice *per* group, males). Data represented as mean *±* SEM (B-N). Circles represent individual mice (B-N). *p* values were determined using one-way ANOVA (B-N). NS, not significant; *p < 0.05; **p < 0.01, ***p < 0.001. Data pooled from two independent experiments with a similar trend.

## Acknowledgements

The authors are indebted to all the members of the Inflammation laboratory (GIMM) for helpful discussions and input, to GIMM core facilities, namely histopathology, bioimaging, flow cytometry, genomics, rodent and advanced data analysis. Luísa Figueiredo for providing the *Cd36^−/-^* mice and Moritz Treek for *Ccr2^−/-^* mice.

## Author contribution

Conceptualization: MPS, JZK, STR

Data Curation: DMF, SS, PF, BD, TP, LG

Formal analysis: DMF, SS, PF, TP, LG

Resources: LH, VM, DS, MJA

Investigation: STR, JZK, MM, SC, MA, JCL

Visualization: STR, JZK, MPS

Funding Support: MPS, EJ, JZK

Project Administration: MPS, JZK

Supervision: MPS, JZK

Writing original draft: MPS, STR

Writing – Reviewing & editing: MPS, STR, JZK

## Data and materials availability

All data are available in the main text or supplementary materials.

## SUPPLEMENTARY MATERIALS

Materials and Methods

Fig. S1 to S22

Supplementary figure legends

Table S1

